# Digging for meaningful connections: associations between root phenotypes and rhizosphere microbial diversity in maize

**DOI:** 10.64898/2026.06.03.725163

**Authors:** Elena Giuliano, Jagdeep Singh Sidhu, Ivan Lopez-Valdivia, Rafaela Feola Conz, Cody L. DePew, Jonathan P. Lynch, Johan Six, Martin Hartmann, Tania Galindo-Castañeda

## Abstract

Drought threatens food security globally. Adaptive root phenotypes and microbiomes can improve maize (*Zea mays* L.) water uptake and tolerance to drought. However, synergisms between root phenotypes and microbiomes remain underexplored. We aimed to investigate the association between varying root phenotypes and rhizosphere microbiomes under field-scale drought.

We grew 22 maize inbred lines in the field under optimal water availability and drought imposed by excluding rain with rainout shelters. We quantified grain yield and measured root architectural and anatomical phenotypes on root crown and cross-section images obtained by laser ablation tomography, respectively. We characterized rhizosphere prokaryotic and fungal communities with DNA metabarcoding of ribosomal markers.

Rhizosphere microbial diversity predominantly associated with root anatomy rather than root architecture. Cortical parenchyma wall width explained 13.1% of the variance of the prokaryotic β-diversity and correlated with grain yield under control conditions. Under the same conditions, number of cortical cell files and metaxylem vessels explained 1.4-2.1% of the variance of prokaryotic and fungal β-diversities. No effect of the root phenotypes was observed under drought. We found 248 significant correlations between microbial taxa abundances and root anatomical phenotypes, especially cortex-related phenotypes such as number of cell files and living cortical area. Overall, a greater number of correlations was found under control conditions.

We identified root phenotypes explaining a small but significant percentage of the variance of the microbial β-diversity, mostly under optimal water availability. We showed that especially root anatomy is associated with rhizosphere microbial diversity in field-grown maize.

## Introduction

Drought, together with extreme heat events, caused a global decline of 9-10% in the national production of cereals between 1964 and 2007. Moreover, severe drought conditions determined up to 90% yield losses for maize (*Zea mays* L.) during its reproductive stage globally (Farré and Faci 2009; Lesk et al. 2016; Leng and Hall 2019; Allakonon and Akponikpè 2022). The predicted increased intensity and duration of drought events will further challenge food production especially in drier areas of the globe (Edenhofer 2015; Maia et al. 2024; Pignède 2025). Harnessing beneficial root-microbe interactions has been proposed as a sustainable approach to improve crop water uptake while protecting the plants from drought. Under water deficit stressed plants initially increase the root/shoot ratio and the allocation of carbon belowground. This can influence rhizosphere microbial assemblies, which can positively affect nutrient cycling and stress tolerance (Williams and de Vries 2020; Wang et al. 2021; Ahmad et al. 2022; Parasar et al. 2024; Etesami and Chen 2025). However, the role that root anatomical and architectural phenotypes play in the establishment and maintenance of the interaction between plants and rhizosphere microbes has only recently started being explored (Galindo-Castañeda et al. 2022; Hartwig et al. 2025; Lattacher et al. 2025).

Root anatomy and architecture are known to be important for soil exploration, resource acquisition and microbial root colonization, and are variable across modern maize lines (Galindo-Castañeda et al. 2019; Lynch et al. 2021, 2022; Lynch 2022). Adaptive root anatomical and architectural phenotypes have been identified in maize under drought. For example, larger (or longer) root cortical cells (Chimungu et al. 2014a; Sidhu and Lynch 2024), thicker cortical parenchyma walls (Sidhu et al. 2024), multiseriate cortical sclerenchyma (Schneider et al. 2021), fewer cortical cell files (Chimungu et al. 2014b), and increased cortical aerenchyma (i.e., pockets of air in the root cortex) (Zhu et al. 2010; Chimungu et al. 2015) are associated with greater plant growth and yield under drought. Moreover, roots growing steeper displaying fewer nodal roots and longer and fewer lateral roots can better reach water accumulated in deep soil layers and reduce competition for resources (Lynch 2013; Zhan et al. 2015; Gao and Lynch 2016).

Non-pathogenic microbes are proposed to protect plants under drought (Jochum et al. 2019). For example, Actinobacteria are defined as “aridity-winners” (Marasco et al. 2021) because they are abundant under drought conditions, not only due to aridity resistance mechanisms but also because they can be positively affected by plant drought-triggered responses, such as changes in root iron homeostasis as observed in sorghum (Xu et al. 2021). Actinobacteria have been reported as potential plant growth-promoting (PGP) bacteria for their ability to increase nutrient availability in soil by contributing to nutrient cycling, to support plant growth by producing phytohormones (e.g., indole-3-acetic acid), and to stimulate antioxidative responses and stress-response gene expression in plants exposed to drought (Boubekri et al. 2022; Narsing Rao et al. 2022; Ebrahimi-Zarandi et al. 2023). Some *Bacillus* spp. can stimulate shoot and root growth (Vardharajula et al. 2011), for example by increasing root length, root surface area and root tip number when inoculated into maize seedlings subjected to drought (Jochum et al. 2019). Some *Pseudomonas* strains release extracellular polymeric substances which can protect the roots and the rhizosphere environment from desiccation (Costa et al. 2018; Ali et al. 2024). Moreover, arbuscular mycorrhizal fungi (AMF) can improve soil exploration and nutrient uptake under drought and can influence the cellular transport of water through the induced expression of aquaporins in plants (Quiroga et al. 2017; Tang et al. 2022).

Recent studies have reported the link between root phenotypes - such as root cortical and stele areas, lateral root branching density and length, root diameter - and rhizosphere microbes (Pérez-Jaramillo et al. 2017; Yu et al. 2018; Galindo-Castañeda et al. 2019, 2025; Zai et al. 2021; Wang et al. 2024; Giuliano et al. 2026). Others have hypothesized possible mechanisms driving the interaction between root phenotypes and microbes (Lynch et al. 2021; Galindo-Castañeda et al. 2022). For example, root anatomical phenotypes, such as cortical aerenchyma and living cortical area, were linked to changes in rhizosphere microbial community composition and had significant associations with the colonization of maize roots by *Fusarium* and arbuscular mycorrhizal symbionts under low phosphorus availability (Galindo-Castañeda et al. 2019, 2023). As hypothesized by Galindo-Castañeda et al. (2022), the presence of cortical aerenchyma could affect rhizosphere microbiomes by reducing transport through cortical cells, thereby changing root exudation levels. Moreover, Yu et al. (2021) found that three root zones along the longitudinal axis of maize roots differing for developmental stage and anatomy harbor different microbial communities. Root architecture can also shape rhizosphere microbiomes, as phenotypes such as root angle and number of axial roots, lateral roots and root hairs can influence nutrient distribution and microbial niches (Galindo-Castañeda et al. 2022; Quattrone et al. 2024).

Despite recent advances in our knowledge regarding the interaction of root anatomical and architectural phenotypes with rhizosphere microbes, their interplay under drought and agriculturally relevant growth conditions are unclear. Furthermore, to the best of our knowledge, the relative importance of root anatomy compared to root architecture for microbial associations under drought has not been studied in crops. Therefore, identifying not one phenotype but a combination of phenotypes, or integrated root phenotypes, under stress (Klein et al. 2020) might offer a valuable perspective on understanding the relative importance of root anatomical and architectural phenotypes for microbial associations. The present study addresses these knowledge gaps by investigating whether the natural variation for root anatomical and architectural phenotypes across 22 maize inbred lines (IBM population, B73 x Mo17; Lee et al. 2002) grown in the field under water deficit and unstressed conditions would be associated with plant performance and rhizosphere microbial diversity. Specifically, we hypothesized that (i) root anatomy and architecture significantly affect the structure of rhizosphere microbial communities under varying water availability; (ii) individual root phenotypes are associated with the relative abundance of microbial taxa depending on water availability and yield performance under stress; (iii) contrasting yield performance among genotypes are associated with differences in root phenotypes and microbiomes.

## Materials and methods

### Plant material

This study investigated 22 intermated recombinant inbred lines (RILs of the IBM population, B73 x Mo17) of maize (*Zea mays* L.) selected from the IBM panel of the University of Wisconsin (Zhu et al. 2005; Sidhu et al. 2024) for their potential performance differences under drought. The IBM lines are genetically distinct lines which originate from the same two parents. Even though IBM lines were not developed specifically for root phenotypes, they have been extensively used for root phenotyping studies to prevent genetics-dependent confounding factors that would have been present if genetically distant or locally adapted genotypes were used (Zhu et al. 2010; Saengwilai et al. 2014; Galindo-Castañeda et al. 2018).

### Set-up of the field experiment

The experiment ran from May to September 2021 in the Russell E. Larson Research and Education Center of The Pennsylvania State University, USA (Rock Springs (PA), 40°42’40.915’’N, 77°, 57’11.120’’W, 366 m.a.s.l.), which is characterized by a Hagerstown silt loam, mixed, fine, mesic Typic Hapludalf soil according to USDA (Haplic Luvisol, World Reference Base classification, WRB), whose main physiochemical properties are reported in Fig. S1. The experiment was performed at two field locations situated 29 m apart (Fig. S2). As described by Sidhu et al. (2024), drought was imposed by deploying rainout shelters (ROS) during rainfall events to exclude precipitation starting 20 days after planting. Under control (unstressed) conditions, rainfall was complemented by drip irrigation based on visual assessment of soil moisture and plant water status by leaf rolling. The successful establishment of water deficit conditions was confirmed by a significant reduction of bulk soil gravimetric water content (-25.2%, Fig. S1) and grain yield (-42.6%, Fig. S3a), together with a genotype-dependent decrease of plant shoot biomass (Fig. S4). As described by Sidhu et al. (2024), the first location (termed Short ROS) followed a split-plot designed consisting of four 22.9 m long and 7.3 m wide blocks for each treatment (drought and control conditions) in which 6 out of 22 genotypes (IBM007, IBM017, IBM200, IBM313, IBM317, IBM365) were arranged in three-row sub-plots and randomized. In this case, plants of the middle row were sampled to diminish edge effects. The second location (termed Long ROS) followed a modified split-plot design consisting of two 44 m long and 9 m wide adjacent fields (one for each treatment) containing 4 blocks in which 16 out of 22 genotypes (IBM071, IBM090, IBM097, IBM106, IBM153, IBM167, IBM181, IBM182, IBM199, IBM201, IBM230, IBM301, IBM344, IBM345, IBM351, IBM352) were randomly assigned to one-row plots. In both locations the plots had a length of 4.6 m, a row spacing of 76 cm and a plant spacing of 18 cm with a planting density of approximately 73,099 plants per hectare. All plots were fertilized with 157 kg ha^−1^ N (46-0-0 urea) according to local recommendations. Pre-emergence herbicides (Acuron, 5.9 l ha^−1^) and pesticides were applied to control diseases and weeds. We combined the data from both field locations (Short ROS and Long ROS) to increase the analytical power and the plant response and phenotypic variability by using a greater total number of genotypes. We have considered field-dependent physiochemical differences (Fig. S1, Table S1) in the data normalization and statistical analysis as described in the Statistical analyses section and in Methods S1.

### Bulk soil sampling

Bulk soil was sampled to characterize physiochemical parameters and resident bulk soil microbiomes (method description in Methods S2), as previously described by Giuliano et al. (2026) but adapted to the field design of this study. Specifically, four weed-free locations between three-row plots in the inner corners and one location in the middle of each block were chosen for the Short ROS fields. Five successive points were sampled horizontally between two one-row plots in the middle of each block of the Long ROS fields. A minimum distance of about 30 cm from the closest maize plant was maintained. A 20 cm-deep sterilized (bleach and 70% ethanol) soil auger with a 2 cm diameter was used for sampling. Five samples per block were collected over two consecutive days (2 weeks after rhizosphere sampling), pooled in sterile bags for a total of sixteen bulk soil samples (2 field locations x 2 treatments x 4 replicates), and kept on dry ice until -80°C storage and shipment to ETH Zurich in sterile bags (Nasco, Whirl-Pak^®^, USA) on dry ice.

### Plant performance measurements

Maize plants were harvested at flowering (after about 70 days of drought) by excavating 15-20 cm-deep root crown according to the shovelomics method (Trachsel et al. 2011) over 3 days during the third week of August 2021. Plant shoot was excised and dried (∼60°C for ∼72 h) to measure biomass. Yield was recorded when the growing season ended.

### Rhizosphere soil sampling

Following the protocol of Galindo-Castañeda et al. (2023) and Giuliano et al. (2026), we packed the excavated 15-20 cm-deep root crown of one plant out of three (2 treatments x 4 replicates x 22 genotypes = 176 samples) in paper bags to avoid cross-contamination and sampled it for anatomy and rhizosphere microbiome characterization. Four to six root segments of the outermost belowground whorl (i.e., roots from the youngest belowground node, the easiest to reach during sampling) were excised with ethanol-sterilized tools. Brace roots were avoided. After manual shaking to eliminate excess soil, -80°C storage and shipment to ETH Zurich in sterile bags (Nasco, Whirl-Pak_®_, USA) on dry ice, roots were thawed on ice and a representative portion was washed in 40 ml of autoclaved phosphate buffer (6.75 g of KH_2_PO_4_, 8.75 g K_2_HPO_4_, 200 μl Tween 20 in 1 l water) by using a vortex for 2 minutes to isolate the rhizosphere soil following the same protocol described by Giuliano et al. (2026) adapted from Lundberg et al. (2012), Simmons et al. (2018) and McPherson et al. (2018). The soil and root debris were removed by filtering the solution through an autoclaved 1 mm stainless-steel mesh (Huayue, China). After centrifuging (4643 g, 4°C, 10 min; Refrigerated Centrifuge Sigma 4-16KS, Sigma, Germany) and discarding the supernatant to obtain a drained soil pellet, DNA was extracted from it on the same day. Bulk soil (3 g) underwent the same procedure for comparability (Methods S2).

### Characterization of root anatomy and architecture

The root architecture of the whole root crown was measured with ImageJ (Schneider et al. 2012; Sidhu and Schneider 2024) through a tailored plugin in ObjectJ (Vischer et al. 2015). As explained by Giuliano et al. (2026), two root segments about ∼4 cm long were collected at about 3 cm from the base of the stem using the same roots selected to sample rhizosphere soil (n = 176). These roots segments were washed and stored in 75% ethanol (*v/v*, in H_2_O) before slicing them with laser ablation tomography (protocol by Strock et al. 2019) to obtain one to four cross-sectional images per sample subsequently processed with *RootScan* v.2.4 (Burton et al. 2012). Cortical parenchyma wall width (CPW, i.e., thickness of the walls of parenchyma cells in the cortex) was measured on root cross-section images of only six selected genotypes (three plants per genotype and treatment) by using ImageJ with the ObjectJ plugin as described by Sidhu et al. (2024). CPW was measured on single cell files and then averaged across all cell files to obtain one measure per sample. Since CPW requires a more laborious measurement at a finer resolution, we considered the same six genotypes used in a parallel study focused only on CPW (Sidhu et al. 2024). All root phenotypes measured in this study are listed in Table 1.

**Table 1.**
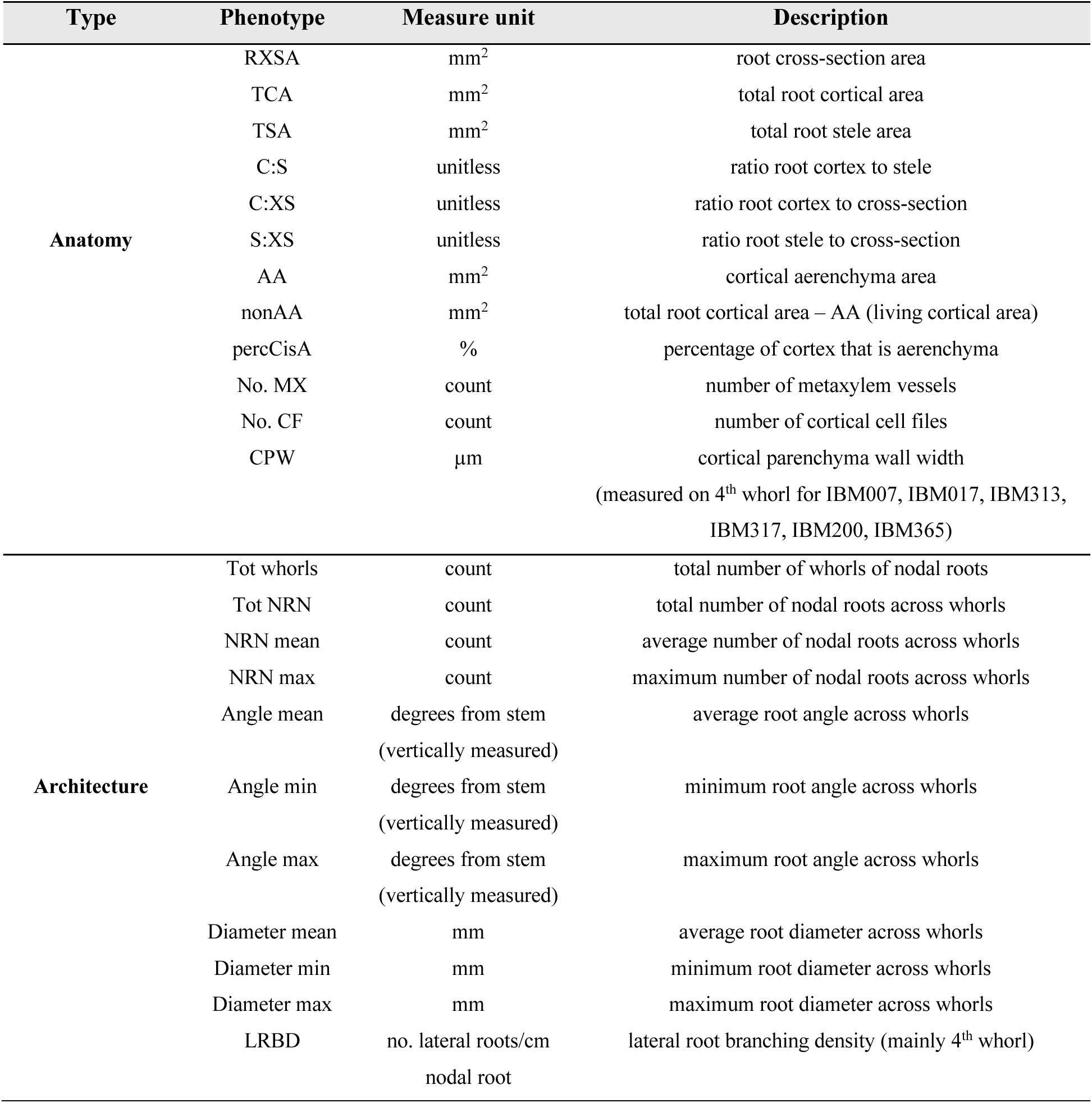
List and details of the anatomical and architectural phenotypes measured.

### Nucleic acid extraction from rhizosphere and bulk soil

As described for a previous experiment (Giuliano et al. 2026), DNA was extracted from moist bulk and rhizosphere soil (250 mg ± 2 mg) following manufacturer’s instructions (DNeasy® PowerSoil® Pro Kit, QIAGEN, Hilden, Germany) with the support of QIAcube Connect (Qiagen). Two 40 s cycles of bead beating and lysis at 5.5 m s^−1^ (FastPrep-24™ 5G, MP Biomedical, Santa Ana, USA) were added to the standard protocol. Extracted DNA was spectrophotometrically checked for concentration and purity (QIAexpert system, Qiagen) and normalized to 10 ng μl^−1^ by using the QIAgility instrument (Qiagen).

### DNA metabarcoding of ribosomal markers

As reported by Giuliano et al. (2026), PCR amplification of ribosomal markers was carried out in a total of 25 μl reaction mix (40 ng of DNA template, 1x GoTaq® Colorless Master Mix (Promega, Madison, WI, United States), 0.4 μM of primers (Microsynth, Balgach, Switzerland), 0.5 mM (prokaryotes) or 1 mM (fungi) of MgCl2 (Promega), prepared with the QIAgility system. For the V3-V4 region of the 16S rRNA gene (prokaryotes), primers 341F (5’-CCTAYGGGDBGCWSCAG-3’) and 806R (5’- GGACTACNVGGGTHTCTAAT-3’) were used (Frey et al. 2016). For the ITS2 region of the rrn operon (fungi), primers 5.8S-Fun (5’-AACTTTYRRCAAYGGATCWCT-3’) and ITS4-Fun (5’- AGCCTCCGCTTATTGATATGCTTAART-3’) were used (Taylor et al. 2016). The PCR was carried out with the following cycling conditions (C1000 Touch and S1000 Thermocyclers, Bio-Rad Laboratories, Hercules, USA): first denaturation at 95°C for 3 minutes, 30 (prokaryotes) or 35 (fungi) cycles of denaturation at 95°C for 40 seconds, annealing at 58°C (prokaryotes) or 55°C (fungi) for 40 seconds, elongation at 72°C for 1 minute and a final elongation at 72°C for 10 minutes. After checking with gel electrophoresis (QIAxcel Advanced, Qiagen) and pooling the three technical replicates produced for each sample, an indexing PCR followed by purification, concentration measurement, and pooling was performed at the Functional Genomic Center Zurich (FGCZ, Switzerland). Lastly, the Illumina Miseq platform (Illumina, San Diego, CA, USA) with the v3 chemistry (PE300) was used to sequence the products of the indexing PCR.

### Bioinformatics

Amplicon sequencing data were processed with a customized bioinformatic pipeline mostly built with VSEARCH (Rognes et al. 2016). Briefly, after checking sequences for PhiX 174 residual fragments with Bowtie2 (Langmead and Salzberg 2012) and trimming the primers with CUTADAPT accepting 1 mismatch (Martin 2011), paired-end reads were merged (*fastq_mergepairs* function), quality filtered allowing a maximum expected error of one (*fastq_filter* function) (Edgar and Flyvbjerg 2015), dereplicated (*derep_fulllength* function, VSEARCH), and delineated into amplicon sequence variants (ASVs) with the UNOISE method (Edgar 2016a) (*cluster_unoise* function, VSEARCH, alpha = 2 and minsize = 4 to remove singletons, doubletons and tripletons). Chimera identification and removal with the UCHIME2 algorithm (Edgar 2016b) (*uchime3_denovo* function, VSEARCH) and biological target verification with Metaxa2 (Bengtsson-Palme et al. 2015) for prokaryotes and ITSx (Bengtsson-Palme et al. 2013) for fungi were performed before mapping the quality-filtered reads against the verified ASV sequences (maxhits 1, maxaccepts 0, maxrejects 32, minimum identity 97%) with the *usearch_global* function (VSEARCH). The SILVA v138 database (Quast et al. 2013) and the UNITE v.9.0 database (Nilsson et al. 2019) were applied on verified prokaryotic and fungal ASVs, respectively, to taxonomically classify them by using a confidence cutoff of 0.8 with the SINTAX algorithm (Edgar 2016c) in VSEARCH.

### Statistical analyses

R Studio v.2026.01.2+418 (R v.4.5.2, R Core Team, 2025) was used to perform the analyses. P-values (P) < 0.05 were considered significant. A q-value threshold of 0.05 was used for all analyses except for the correlations between root phenotypes and microbial taxa (q-value < 0.1). All permutation-based statistical and correlational methods used 9999 permutations (i.e., PERMANOVA, PERMDISP, Mantel test). Information regarding sample number and missing values (NAs) for the main analyses are reported in Table S2. We applied analytical and statistical approaches analogous to those described by Giuliano et al. (2026), adapted here for the current study. A detailed description is reported below.

### Plant and soil data curation

Relative yield change was calculated as [(yield Drought – yield Control) / yield Control) × 100] by first pairing the yield data of corresponding biological replicates for each treatment and then averaging by genotype. We used the relative yield change to evaluate the successful establishment of the experimental drought and measure its impact on plant performance. Moreover, calculating the relative change with respect to the control conditions allowed us to reduce biases dependent on the fact that, in the LongROS, grain yield positively correlated with plant vigor under control conditions (Pearson, r = 0.32, P < 0.05) and grain yield under drought (Pearson, r = 0.67, P < 0.001) and greater vigor under control conditions corresponded to a greater relative yield change (Pearson, r = -0.37, P < 0.05). After data curation and gap filling where possible, yield relative change and root phenotypic data were normalized (z-scores, function *scale*, centering and scaling) by field, while bulk soil data were normalized for the entire dataset (Methods S1). Z-scores were calculated by subtracting the column mean from each sample value and then dividing by the sample standard deviation. This approach was used to make variables with different value ranges and units comparable by having a mean of zero and a standard deviation of one for each column.

### Statistical analyses on yield, root phenotypic and bulk soil data

For yield and root phenotypes, Linear Mixed-effects Models (LMMs, function *lme*, package *nlme* 3.1-168; Zuur et al. 2009; Pinheiro 2022) were used to test the effect of treatment and genotype including field or block as random effect (Methods S3 for model equations). Estimated marginal means (EMMs, function *emmeans,* package *emmeans* v1.11.2) and 95% confidence intervals built on the LMM model were used to evaluate the genotype effect on percentage yield change. Genotypes with EMM-based 95% confidence intervals excluding zero were considered significantly different form the zero baseline (yield under control conditions), while the opposite applied if their 95% confidence intervals included zero. A pairwise test (function *pairs*) with Tukey p-value adjustment was performed on root phenotype EMMs to identify specific treatment x genotype differences (Methods S3). Pearson correlations were run between root phenotypes and grain yield with the function *cor.test* (package *stats* v4.5.2). For bulk soil parameters, a non-parametric multivariate analysis of variance based on permutations (PERMANOVA, function *adonis2*, package *vegan* v2.7-2; Anderson 2001) followed by PERMDISP to test homogeneity of variance (functions *betadisper* and *permutest*, package *vegan* v2.7.2; Anderson 2006; Anderson et al. 2006) was used to test the effect of treatment × field (Methods S3). P-values were adjusted for multiple testing using the false-discovery rate (FDR) method (Benjamini and Hochberg 1995; function *p.adjust*). To test the relationship between cortical thickness, cell size and file number, we calculated the root radius as square root of the root cross-sectional area divided by pi (π, 3.141593), and the stele radius as square root of the total stele area divided by pi (π). We obtained the cortical radius (cortical thickness) by subtracting the stele radius from the root radius. Subsequently, we divided the cortical thickness by the cell file number to estimate the average radial size of cortical cells across cell files. These estimated phenotypic measurements were not included in the main analyses as they were used only to evaluate the relative contribution of radial cell size and cell file number to cortical thickness. We used the function *chart.Correlation* (package *PerformanceAnalytics* v2.0.8) to plot the correlations with the statistics between all pair-wise combinations of total cortical area, cortical thickness, cell file number and radial cell size (Pearson, *cor.test*).

### Clustering based on relative yield change and root phenotypes

Partitioning Around Medoids (PAM) k-medoids clustering (package *cluster* v2.1.8.1; Maechler et al. 2019) was used to identify clusters based on relative yield change and root phenotypes, as similarly done by Martins et al. (2024) and Klein et al. (2020). Outliers or NAs were not removed or estimated to avoid reducing the number of samples or inserting biases. The within-sums-of-squares (WSS) plot (function *fviz_nbclust*, package *factoextra* v1.0.7) with the elbow method was used to identify the optimal number of clusters, whose robustness was checked with the Silhouette score (function *silhouette*, package *cluster* v2.1.8.1). LMMs on the original data were used to confirm statistical differences (P < 0.05) between yield-based clusters. For yield-based grouping, four clusters were identified and grouped by two based on their similarity. Genotypes having at least 50% of their available biological replicates in two paired clusters were considered being part of that super-group containing those two clusters. Genotypes with only one yield change value (IBM317, IBM097, IBM090) or not enough corresponding replicates to calculate it (IBM344, IBM352) were not included in the clustering. One cluster included lower-performing genotypes and the other one well-performing ones based on their high or low relative grain yield reduction under drought (Fig. S5, S6). Since the genotype IBM071 (classified as well-performing) had a significantly lower grain yield than IBM181 and IBM182 (classified as lower-performing) under control conditions (LMM, P < 0.05), it was removed from the well-performing cluster to avoid introducing confounding factors related to the original yield potential which are difficult to disentangle. To balance the two groups, IBM182 was removed as well having only two replicates. This selection resulted in a total of six genotypes with comparable grain yield and vigor (LMM, P > 0.05) under control conditions and displaying no links between their yield and vigor under optimal water availability and their performance under drought (Fig. S3b-d). Since finding the cause of the observed yield differences was not in the scope of this work, these performance groups were only used to explore trends in root phenotype-microbiome associations possibly linked to drought stress response. The anatomy-based k-medoids clusters (higher Silhouette score compared to architecture) were plotted together with their Principal Component Analysis (PCA, function *prcomp*, package *stats* v4.5.2).

### Microbial diversity calculation and effect of the main experimental variables

Sequencing depth was checked with rarefaction curves (*rarecurve* function, package *vegan* v2.7-2; Oksanen et al. 2024) (Fig. S7). As previously suggested by Schloss (2024) and implemented by Longepierre et al. (2021), ASV count tables were 100-fold iteratively subsampled to standardize for sampling and sequencing depth differences. Shannon index (α-diversity, function *diversity*, package *vegan* v2.7-2) and Bray-Curtis dissimilarity (β-diversity, function *vegdist*, package *vegan* v2.7-2) were calculated as average across the 100 subsampled matrixes of ASV count data. Unconstrained variability across samples was displayed through Principal Coordinate Analysis (PCoA, function *cmdscale*, package *stats* v4.5.2). Canonical analysis of Principal Coordinates (CAP) was used to constrain the β-diversity by performance groups (function *CAPdiscrim*, package *BiodiversityR* v2.17-4). For the taxon level analysis, the averaged 100 subsampled matrixes of ASV counts were converted into relative abundances by sample and summed up by genus. PERMANOVA (function *adonis2,* package *vegan* v2.7-2), PERMDISP (functions *betadisper* and *permutest,* package *vegan* v2.7-2) and pairwise PERMANOVA (function *pairwise.adonis2,* package *pairwiseAdonis* v0.4.1) were used to test the effect of treatment, genotype and their interaction on α- and β-diversity and on genera relative abundances (Methods S3 for model equations). For consistency with statistical analyses applied to yield and root phenotypes and because of its effect on microbial communities, field was used as *strata* in PERMANOVA. P-values were adjusted with the FDR method (function *qvalue,* package *qvalue* 2.40.0; Storey 2002). The plant beneficial and phytopathogenic bacteria databases provided by Li P. et al. (2023) and literature search were used to identify potential functions for each microbial genus.

### Links between root phenotypes and microbiomes at community and genus level

PERMANOVA, Mantel test (function *mantel*, package *vegan* v2.7-2, Spearman method) and Spearman correlations (*cor.test* function, package *stats* v4.5.2) were used to investigate the link between plant yield, root phenotypes and microbiomes (Methods S4 for additional analyses performed). At the community level, PERMANOVA was used to test the effect of treatment, genotype and root phenotypes on microbial β-diversity. Mantel test was used to correlate microbial β-diversity (Bray-Curtis dissimilarity) with root phenotypes (Euclidean distance). At the genus level, the 100 averaged subsampled matrices of ASV data previously converted into relative abundances and aggregated by genus were filtered for sparsity by retaining only the genera with relative abundance greater than zero in at least 75% of the samples. The dataset was split by treatment, and each subset was z-transformed by field (function *scale*, centering and scaling) and further split by performance group to run Spearman correlations between root phenotypes and microbial taxa relative abundances. P-values were adjusted for each phenotype across all microbial genera (function *qvalue,* package *qvalue* 2.40.0; Storey 2002).

## Results

### Variation in root phenotypes under drought, with linkages to yield

Variation among replicates and plasticity in response to drought was observed for both anatomy and architecture across all 22 maize genotypes. Maize genotypes displayed significant (LMM, P < 0.05) differences in all anatomical phenotypes, except for number of cell files (No. CF), in dependence of the treatment (Fig. 1a, Fig. S8, Table S3). Total number of whorls, nodal root number, minimum angle and average diameter displayed higher plasticity and were significantly (LMM, P < 0.05) reduced under drought as shown by the orange color of the heatmap cells in correspondence of drought in Fig. 1b (Table S3).

**Fig. 1.**
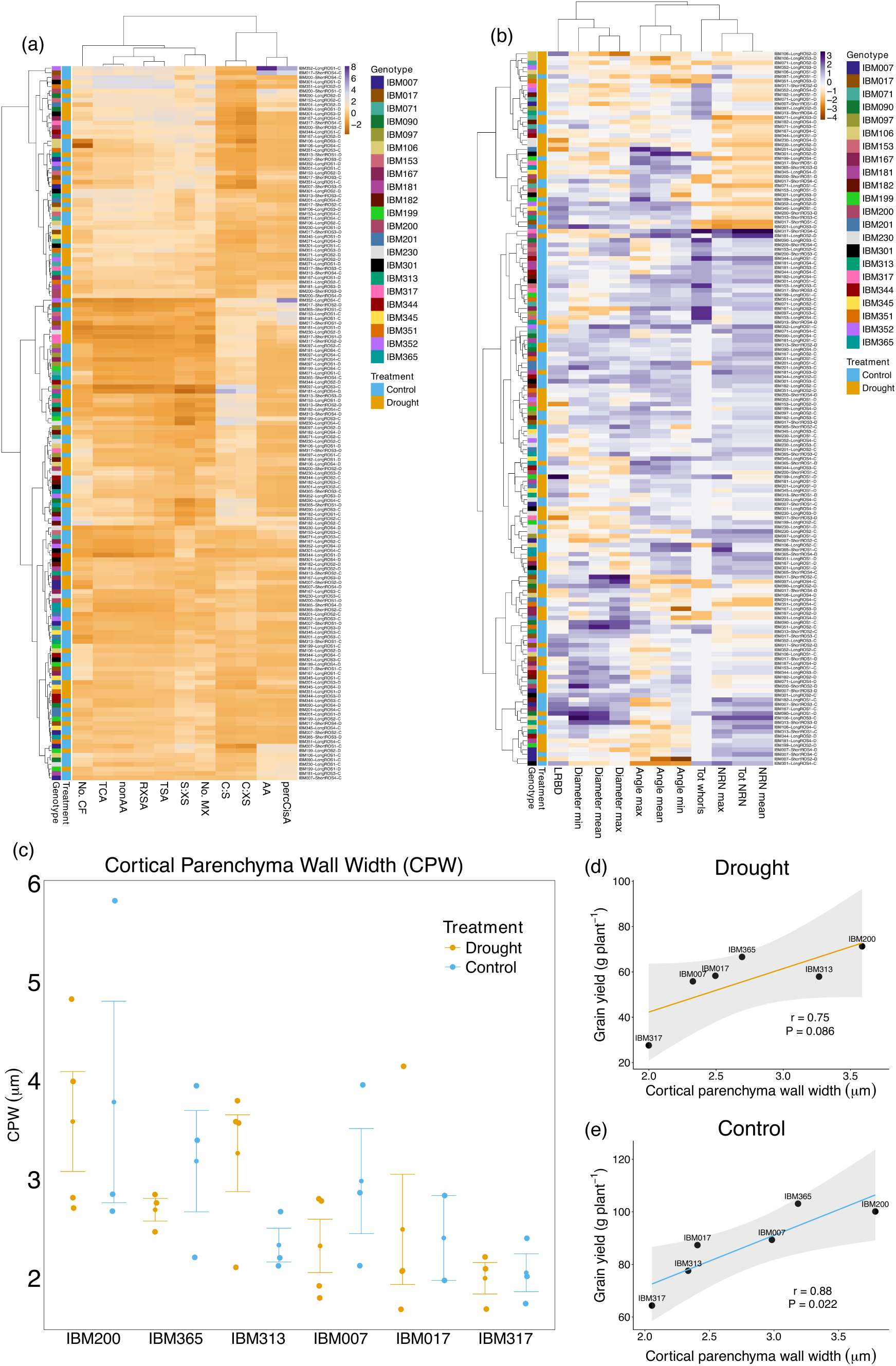
Natural variability of root anatomical and architectural phenotypes and correlations with yield. Heatmaps showing the hierarchical clustering of samples (sample information including genotype, field, replicate and treatment are displayed on the y-axis, right side) based on (a) z-transformed anatomical (x-axis) or (b) architectural (x-axis) measurements across treatments, genotypes and replicates (y-axis). Expression of the anatomical phenotype cortical parenchyma wall width (CPW) across six genotypes and two treatments (c). Scatter plots reporting the positive correlation (linear model) between CPW and grain yield under drought (d) and control conditions (e). Each dot represents the average CPW and grain yield of corresponding biological replicates (n=1 to 4). The Pearson correlation coefficient (r) and p-value (P) are reported.

Regarding links with yield, root cross-section area (RXSA), total stele area (TSA), root cortex to cross-section area (C:XS), living cortical area (nonAA) and number of metaxylem vessels (No. MX) were significantly associated with grain yield under drought, explaining between 2-7% of the variance compared to 59% explained by the genotype (PERMANOVA, P < 0.05, Table S4). Moreover, cortical parenchyma wall width (CPW) ranged from 1.68 to 4.83 µm under drought and from 1.74 to 5.82 µm under control conditions across six genotypes (Fig. 1c), and was positively correlated with grain yield under control conditions (Fig. 1d,e).

### Effect of drought and maize genotype on rhizosphere microbial diversity

Drought significantly reduced rhizosphere prokaryotic and fungal α-diversity (Shannon index at ASV level, Table 2, Fig. S9) and affected bulk soil prokaryotic and fungal β-diversities (PERMANOVA, P < 0.001). A genotype-dependent drought effect was observed for rhizosphere prokaryotic and fungal β-diversities (Bray-Curtis dissimilarity at ASV level, Fig. 2a,b) and 15 prokaryotic genera, but not fungal genera, were affected by genotype (Table S5). Genotype and drought alone explained a significant proportion of the variance for prokaryotic (12-22%) and fungal (20-19%) community structures (Table 2). Moreover, a field effect on bulk soil and rhizosphere microbial β-diversity was observed (PERMANOVA, P = 0.0001; Fig. 2a,b; Fig. S10).

**Fig. 2.**
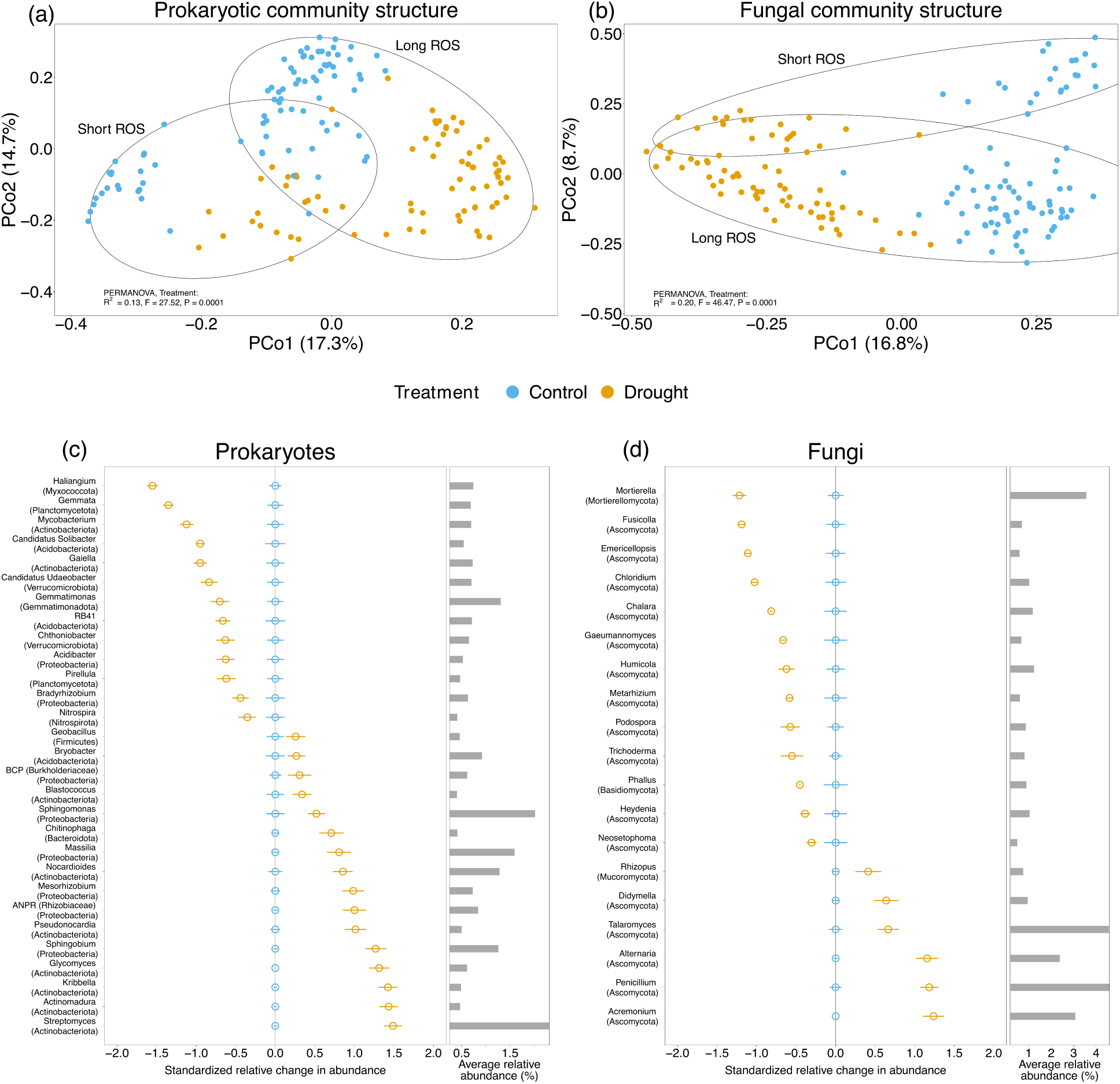
Effect of drought on rhizosphere prokaryotic and fungal communities. Principal Coordinates Analysis (PCoA) on Bray-Curtis dissimilarities (ASVs) showing the separation between treatments and field for prokaryotic (a) and fungal (b) β-diversities. The percent variation explained by each axis is reported in brackets, and the output of the permutational multivariate analysis of variance (PERMANOVA) is shown in the plot. The most abundant (mean relative abundance across samples > 0.3 %) prokaryotic (c) and fungal (d) genera, followed by the respective phylum, significantly affected (PERMANOVA, q < 0.05) by drought are reported with their relative change in abundance compared to control conditions (zero line with standard errors). Displayed abundance levels were calculated as (z-transformed relative abundance under drought) – (z-transformed relative abundance under control conditions) and reported with their standard errors. The right-hand sides of the plots illustrate average relative abundances per genus across all rhizosphere samples and treatments.

**Table 2.** Impact of drought, maize genotype and their interaction on prokaryotic and fungal α-diversity (Shannon index) and β-diversity (Bray-Curtis dissimilarities).

| Prokaryotes |  |  |  |  |  |  |
| --- | --- | --- | --- | --- | --- | --- |
| Experimental variables | α-diversity | | | β-diversity | | |
|  | R<sup>2</sup> | F | P | R<sup>2</sup> | F | P |
| Treatment | 0.500 | 165.2 | **0.0001\*** | 0.126 | 27.5 | **0.0001** |
| Genotype | 0.080 | 1.3 | 0.1945 | 0.217 | 2.2 | **0.0111** |
| Treatment x Genotype | 0.069 | 1.1 | 0.3841 | 0.124 | 1.3 | **0.0014** |
| Fungi |  |  |  |  |  |  |
| Experimental variables | α-diversity | | | β-diversity | | |
|  | R<sup>2</sup> | F | P | R<sup>2</sup> | F | P |
| Treatment | 0.272 | 56.4 | **0.0001** | 0.200 | 46.5 | **0.0001** |
| Genotype | 0.077 | 0.8 | 0.8691 | 0.185 | 2.05 | **0.0042** |
| Treatment x Genotype | 0.091 | 0.9 | 0.5873 | 0.115 | 1.3 | **0.0120** |

<sup>a</sup>Heteroscedasticity was tested with PERMDISP, and significant results are highlighted with an asterisk (\*). Statistically significant results (PERMANOVA, $P < 0.05$ ) are reported in bold.

At the taxon level, drought significantly affected (PERMANOVA, q < 0.05) 58.8% and 32.5% of the 765 and 588 taxonomically classified prokaryotic and fungal genera, respectively. The most abundant genera affected by drought belonged to the phyla Actinobacteriota, Proteobacteria and Ascomycota (Fig. 2c,d). Moreover, two arbuscular mycorrhizal fungi (AMF, *Glomus aggregatum* and *Glomus indicum*) significantly decreased in abundance under drought (Fig. S11).

### Effect of root phenotypes on rhizosphere microbial diversity

Both anatomical and architectural phenotypes significantly influenced microbial β-diversity across all 22 maize genotypes. No correlations between the observed patterns in prokaryotic and fungal β-diversities and root phenotypes were found (Mantel test, P > 0.05), except for the six genotypes on which cortical parenchyma wall width (CPW) was measured for which patterns of prokaryotic β-diversity correlated with patterns of CPW only under control conditions (Mantel test, ρ = 0.4, P < 0.05). Nevertheless, when testing the effect of the root phenotypes on microbial β-diversity for all genotypes and treatments, root cross-section area (RXSA) and average root angle (Angle mean) or RXSA only significantly affected prokaryotic and fungal β-diversities, respectively. These phenotypes explained a smaller portion of the variance (0.7-0.9%) than the treatment, the genotype and their interaction (12-22%). To assess even small effects of the root phenotypes on microbial β-diversity excluding the treatment effect, we split the data into drought and control conditions. No significant effects of root phenotypes were found under drought. However, under control conditions, number of metaxylem vessels (No. MX) and number of cell files (No. CF) explained 1.4-2.1% of the variance of both prokaryotic and fungal β-diversities, while the minimum root diameter (Diameter min) explained 1.4% of the variance of the fungal community structure in comparison with up to 42% of the variance explained by the genotype (PERMANOVA, P < 0.05, Table S6). Cell parenchyma wall width (CPW) explained 13.1% of the variance of prokaryotic β-diversity across six inbred lines under control conditions only (PERMANOVA, P < 0.001). Also in this case, the genotype explained a higher portion of the variance (37.3%).

### Correlations between root phenotypes and relative abundance of microbial genera

Two performance groups (i.e., three well-performing and three lower-performing genotypes) were identified based on grain yield change-dependent k-medoids clustering (see Materials and Methods, Fig. S5, S6 and Notes S1 for more details on the yield data and clustering). Although only aerenchyma significantly changed between these groups (i.e., greater aerenchyma in the lower- vs. well-performance genotypes) (Fig. S12, S13, Notes S2), rhizosphere prokaryotic and fungal community structures differentiated between well- and lower-performing genotypes (Fig. S14). Therefore, we run correlations between root phenotypes and microbial relative abundance not only across all 22 genotypes together but also for the subset of six genotypes clustered in the two performance groups.

Overall, significant (Spearman, q < 0.1) correlations were more numerous under control conditions and root anatomical phenotypes revealed a greater number of correlations (n=248) compared to architectural phenotypes (n=58) (Tables S7, S8, S9, S10). When investigating the correlations only by performance group, well-performing genotypes displayed a smaller number of correlations under drought (n=4) compared to control conditions (n=105), while the opposite pattern was observed for lower-performing genotypes (Tables S9, S10).

In all cases, root angle (Angle mean, max, and min) and root diameter (Diameter mean, max, and min) were the architectural phenotypes most often correlating either positively or negatively with prokaryotic taxa abundances. Overall, the greatest number of significant correlations was found between the relative abundance of prokaryotes and fungi and root cortex-related anatomical phenotypes, such as number of cell files (No. CF), root cross-section area (RXSA), total cortex area (TCA) and living cortical area (nonAA). Specifically, *number of cortical cell files (No. CF)*, which correlated positively with TCA and estimated cortical thickness but negatively with estimated radial cortical cell size (Fig. S15, S16), was associated with 112 prokaryotic and 18 fungal relative abundances under control conditions across all 22 genotypes. Apart from the frequency with which this phenotype was observed, number of cell files negatively correlated with potentially plant beneficial prokaryotic and fungal taxa, such as *Bradyrhizobium*, *Burkholderia-Caballeronia-Paraburkholderia*, *Massilia*, *Mesorhizobium*, *Sphingomonas*, *Rhizophagus*, *Gongronella* (Fig. 3, Supplementary Tables S7, S8). Moreover, *living cortical area (nonAA)*, which is closely related to root cross-section area (RXSA) and total cortical area (TCA), negatively correlated with potential PGP-prokaryotic genera, such as *Bradyrhizobium*, *Deinococcus* and *Pseudolabrys*, in the well-performing genotypes under control conditions (Fig. 3, Table S9). Furthermore, *cortical parenchyma wall width (CPW)* positively correlated with the fungal genus *Saitozyma* and the prokaryotic genus *Azospirillum* across six genotypes under control conditions (Fig. 3).

**Fig. 3.**
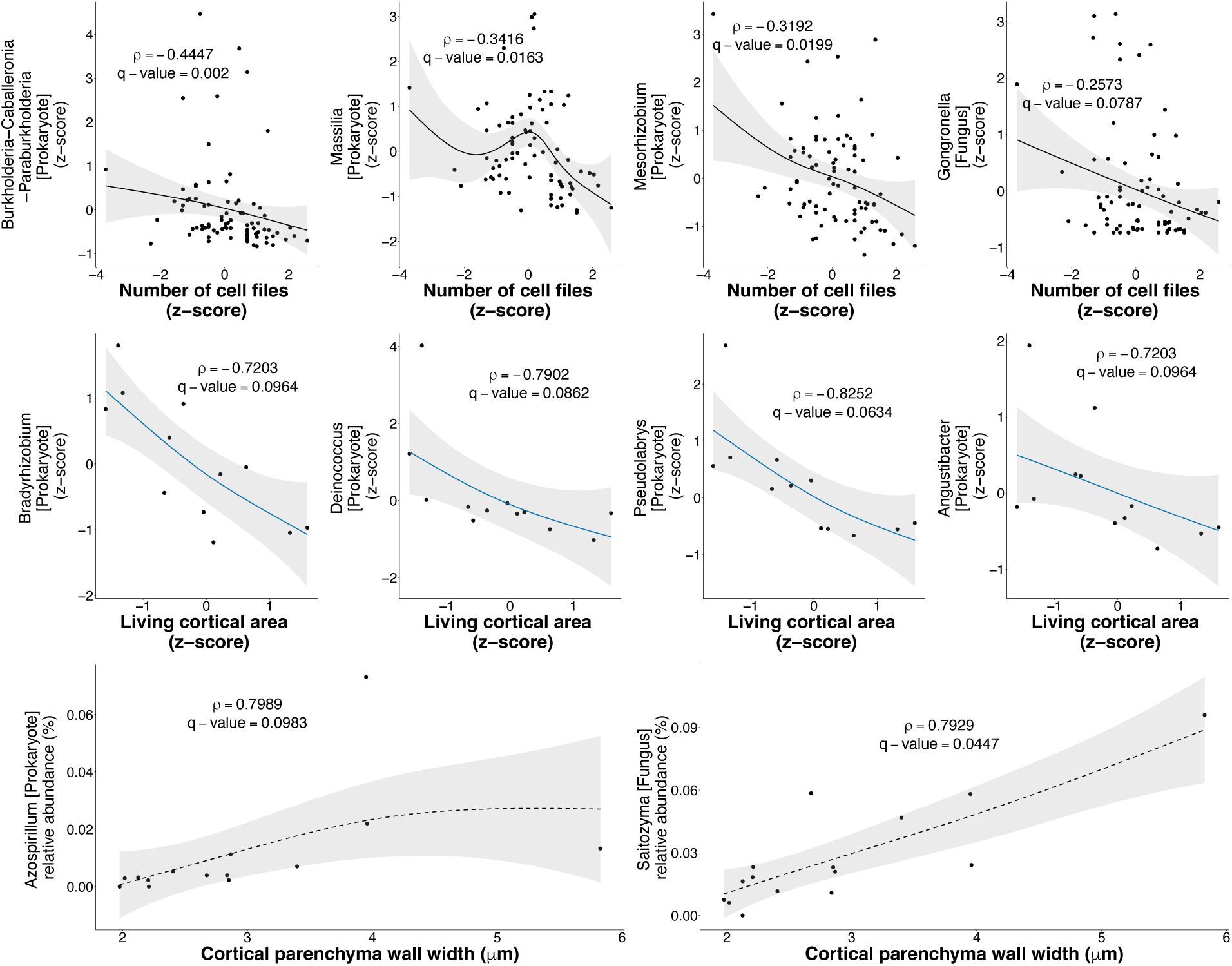
Examples of correlations between root cortex-related anatomical phenotypes and relative abundances of microbial genera under control conditions. Significant (Spearman, q < 0.1) correlations between number of cell files (No. CF), living cortical area (nonAA) and cortical parenchyma wall width (CPW) with the relative abundance of potentially beneficial prokaryotic and fungal taxa are illustrated. Each dot in the scatter plot is a sample available in both the anatomical and microbial datasets. Z-transformed root phenotypic and microbial relative abundance data were plotted, except for the last two plots where raw CPW data and rarefied ASV counts transformed into relative abundances by genus were used. Generalized additive model (GAM), 95% confidence intervals, correlation coefficients (ρ) and q-values are reported in each plot. The continuous black and blue lines and the black dashed lines indicate whether the correlations were calculated on the 22 genotypes, on the well-performing genotypes only and on the six genotypes in which CPW was measured, respectively.

In the lower-performing genotypes under drought, instead, a smaller root diameter, reduced stele area (TSA, S:XS) and number of metaxylem vessels (No. MX) associated with increased abundance of potentially beneficial and non-pathogenic prokaryotic taxa (Table S9). For example, the prokaryotic genus *Acidibacter* positively correlated with cortex to cross-section ratio (C:XS) but negatively with stele to cross-section ratio (S:XS) and number of metaxylem vessels (No. MX).

## Discussion

In the present study we investigated patterns in root anatomy and architecture and rhizosphere microbiomes across 22 field-grown maize inbred lines with varying yield performances under drought. We could address our hypotheses by finding that: (i) number of metaxylem vessels and cell files, cortical parenchyma wall width and root diameter explained 1.4-13.1% of the variance of the β-diversity of prokaryotes and fungi under control conditions but not under drought; (ii) root anatomical and architectural phenotypes displayed 248 and 58 significant correlations with relative abundances of microbial taxa, respectively, whereas correlations were more numerous under control conditions; (iii) yield performance dissimilarities associated with differences in cortical aerenchyma and microbial β-diversity. Therefore, this study represents an advancement in the investigation of the association between root phenotypes and rhizosphere microbiomes.

### Root phenotypes differed among genotypes and treatments

The observed variability in the expression of root anatomical and architectural phenotypes is well-known among maize inbred lines (Burton et al. 2013; Lynch 2013; Lammerts van Bueren and Struik 2017). The genotypes used belong to the IBM population consisting in recombinant inbred lines (RILs), which are genetically distinct genotypes originated from the same two parents (Zhu et al. 2005). This characteristic made these lines suitable for identifying specific associations between adaptive root phenotypes and microbes without introducing genetic biases that would have been present in the case of a comparison between locally adapted genotypes with contrasting genetic backgrounds (Saengwilai et al. 2014).

In addition to the known root phenotypic variability, the IBM genotypes also displayed plasticity in response to drought. Specifically, number of crown whorls and nodal roots, and root diameter were significantly reduced by drought, which are adaptive responses improving rooting depth and root available surface for water acquisition, as previously observed in maize by Gao and Lynch (2016) and Wasaya et al. (2018). Number of cell files (No. CF) was the sole anatomical phenotype whose expression remained stable across genotypes and treatments (Figure 1a, Table S3). Compared to the more plastic root diameter which often correlated with microbial relative abundances under drought in the two performance groups (Table S9), number of cell files displayed the greatest number of associations with rhizosphere microbes but mostly under control conditions (Table S6, S7). This might be an indication that the role of phenotypic plasticity should be explored further in the associations with rhizosphere microbes.

### Drought and maize genotypes shaped rhizosphere microbial diversity

Drought affected both microbial α- and β-diversity but only the effect on β-diversity was modulated by genotype. Our results support previous studies reporting a reduction in prokaryotic and fungal α-diversity (Gao et al. 2023) and changes in prokaryotic and fungal β-diversity due to drought (Meisner et al. 2018; Gao et al. 2023; Li et al. 2025). However, studies focused on maize or other plants (e.g., cowpea, rice, oak) reported an increase in rhizosphere prokaryotic α-diversity (Wu et al. 2024; Halo et al. 2025), with fungi being particularly more resistant to drought compared to prokaryotes (Preece et al. 2019; Tian et al. 2024). Regarding the microbial genera affected, some potential plant growth-promoting (PGP)-bacterial genera, such as *Bradyrhizobium*, *Nitrospira* and *Gemmatimonas* (Antoun et al. 1998; Ayiti et al. 2022; Ma et al. 2024), and fungal genera, such as *Mortierella*, *Emericellopsis* and *Humicola* (Ozimek and Hanaka 2021; Li et al. 2024; Ma et al. 2024), decreased in abundance under drought, as previously indicated for some of them in other crops by Loiko and Islam (2024) and Liu et al. (2025). On the contrary, potentially beneficial drought tolerant bacterial (e.g., *Burkholderia-Caballeronia-Paraburkholderia*, *Massilia*, *Mesorhizobium*, *Sphingobium*, *Sphingomonas*, *Streptomyces*) and fungal (e.g., *Rhizopus, Penicillium*) genera with potential to improve plant tolerance to water limitation (Chaudhary et al. 2018; Luo et al. 2020; Tiziani et al. 2022; Fan et al. 2023; Kihara et al. 2024; Taha et al. 2025) increased in abundance under drought compared to control conditions. Moreover, potential fungal pathogens, such as *Didymella* and *Alternaria* (Corlett 1981; Thomma 2003; Sánchez-Arizpe et al. 2023), increased in abundance under drought as also reported in different crops (Dikilitas et al. 2016). However, it is important to acknowledge the limitation of the metabarcoding approach which does not allow to discern pathogenic taxa and does not provide information on the function of the microbial genera. Therefore, these results should be corroborated by the functional characterization of these genera.

The different maize genotypes explained 22% and 19% of the variance of prokaryotic and fungal β-diversities, respectively. Previous studies have reported similar degrees of variance (i.e., 13-19%) of prokaryotic and fungal community structures explained by 20 to 27 maize inbred genotypes grown either in the field or in the greenhouse (Peiffer et al. 2013; Favela et al. 2021). When investigating 27 up to 500 genotypes or comparing maize lines across a genetic transect, a smaller genotype effect (less taxa affected or 5-7% variance of β-diversity explained) was observed (Walters et al. 2018; Brisson et al. 2019; Schmidt et al. 2020; Meier et al. 2022; Mukhtar et al. 2024; He et al. 2024). A genotype-dependent effect on prokaryotic and fungal diversity was also observed at a smaller scale by using three to six maize genotypes in the greenhouse or in the field (Li et al. 2022; Wen et al. 2025). Therefore, even by using a relatively small number of replicates (n=4) of genetically close lines and considering the influence of field location which is common in microbiome studies (Longepierre et al. 2021; Hartmann and Six 2022), we could observe enough variability across genotypes.

### Root anatomy, more than architecture, affected rhizosphere microbial diversity

Interestingly, we found a greater number of anatomical phenotypes influencing microbial β-diversity (root cross section area, number of metaxylem vessels and cell files) and correlating with microbial taxa relative abundances (n=248) compared to architecture (n=58). This indicate that specific anatomical phenotypes might play a larger role in forming associations with microbes compared to the architectural ones under these field conditions. However, we must acknowledge that characterizing rhizosphere microbiomes and root anatomy on the same root samples or whorls might have resulted in the identification of a greater number of associations between anatomical rather than architectural phenotypes and microbes because of the direct correspondence between samples or locations in the root system. Nevertheless, cortical cell parenchyma wall width (CPW) was measured on the 4^th^ whorl instead of the outermost one and averaged across plant replicates; therefore, this explanation might not apply to all phenotypes. For this reason, future experiments should carefully consider sampling the same whorl or root location for all phenotypic and microbial samples to exclude possible confounding factors.

Root cortex-related anatomical phenotypes were more often associating with rhizosphere microbial diversity, especially under control conditions across all 22 genotypes. Specifically, *number of cell files (No. CF)* explained up to 1.5% of the variance of prokaryotic and fungal community structures and displayed the greatest number of correlations (n=132) of all investigated phenotypes. Interestingly, it negatively correlated with some of the most abundant prokaryotic taxa that increased in abundance under drought (e.g., *Allorhizobium-Neorhizobium-Pararhizobium-Rhizobium*, *Burkholderia-Caballeronia-Paraburkholderia*, *Massilia*, *Mesorhizobium*, *Sphingomonas*). We hypothesize that number of cell files may have influenced the amount of living tissue in the cortex as, especially in combination with cell size and number and aerenchyma, this trait can influence both the endophytic microbe colonization and the travel distance for exudates (Galindo-Castañeda et al. 2019, 2022). Moreover, since a reduced number of cell files is an adaptive trait under drought (Chimungu et al. 2014b), the observed negative correlations between this phenotype and microbial taxa might indicate that its adaptive nature can coincide with the increased abundance of beneficial bacteria. Nevertheless, tradeoffs in the expression of this phenotype can be important as a reduced number of cell files could facilitate water transport in the root but reduces the space available for mycorrhizal colonization which can be beneficial under water deficit (Freschet et al. 2021). Cell file number displayed also a greater positive correlation with total cortical area and estimated cortical thickness across all 22 genotypes within each treatment compared to estimated radial cortical cell size. Moreover, a negative correlation between cell file number and estimated radial cortical cell size was observed (Fig. S15, S16). Therefore, number of cell files might have a greater significant role in determining cortical thickness and overall root diameter in concert with cortical cell size, as reported by Sidhu and Lynch (2024). For this reason, number of cell files, its relationship with other root cortical phenotypes and their direct (e.g., apoplastic space available for fungal colonization) or indirect (e.g., changes in root metabolic costs and carbon rhizodeposition) relative contribution to rhizosphere and endosphere microbial associations merits attention in future studies.

Number of metaxylem vessels (No. MX) and root diameter (Diameter min) also explained a small but significant portion of the variance of microbial community structure across all 22 genotypes under control conditions. Number of metaxylem vessels (No. MX), together with metaxylem wall thickness, affect hydraulic conductivity and root mechanical resistance (Hochholdinger 2009; Li H. et al. 2023; Klein et al. 2024). Therefore, it might be interesting to study the association of these phenotypes with microbes, especially with endosphere microbial communities and root pathogens under drought. Root diameter could affect microbial community assemblies by changing root exudation rate and root surface available for microbial colonization. Indeed, a small diameter in combination with fewer cortical cells ensures a shorter symplastic and apoplastic path for molecule transport towards the rhizosphere and can increase the available root surface for microbial colonization in the rhizosphere (Galindo-Castañeda et al. 2022; Agarwal et al. 2024).

Another anatomical phenotype of interest is the *living cortical area (nonAA)*. This phenotype usually positively correlates with total cortical area (TCA) and root cross-section area (RXSA) (Vanhees et al. 2020), which indeed displayed similar trends in the correlational analysis by negatively correlating with potentially PGP taxa, such as *Bradyrhizobium, Deinococcus* and *Pseudolabrys*, in well-performing genotypes under control conditions. For example, a reduced living cortical area not only reduces the root maintenance costs but can also decrease the availability of apoplastic space for microbial colonization and can reduce the symplastic and apoplastic release of carbon into the rhizosphere depending on the root diameter, the cell size and the cell wall thickness (Dreyer et al. 2010; Sharda & Koide 2010; Galindo-Castañeda et al. 2019, 2022; Lynch et al. 2023). Moreover, since living cortical area was significantly associated to grain yield under drought across all 22 genotypes, investigating how its levels of expression associate with microbes and therefore to plant yield might be of importance in future manipulative studies.

Therefore, compared to studies reporting the influence of individual phenotypes or groups of a few anatomical and architectural phenotypes on the microbial diversity (Dreyer et al. 2009; Pérez-Jaramillo et al. 2017; Rüger et al. 2021, 2023; Galindo-Castañeda et al. 2023), our results highlight the importance of exploring the integration of multiple cortex-related phenotypes and the need of carefully considering tradeoffs of their expression for beneficial and pathogenic microbial interactions.

### Root phenotypes and microbes associated more often under optimal water availability rather than under drought

Root anatomical and architectural phenotypes significantly associated with microbial β-diversity and correlated with microbial taxa relative abundances more frequently under optimal conditions (n=256 out of 306 total significant correlations), such as high water and N availability (Giuliano et al. 2026). This might indicate that high water availability in soil improves microbial diversity and could reduce competition between roots and microbes for water and other nutrients in the rhizosphere since maize plants invest more energy in aboveground growth under optimal conditions (Eziz et al. 2017; Gao et al. 2023). Another explanation can be the previously shown self-sufficiency of adaptive root phenotypes under drought (Lynch 2013), which might not necessarily require plant-driven recruitment of beneficial microbiomes. Vice versa, these results might suggest that root phenotypes play a less relevant role in plant-microbe associations under drought, especially in these closely related maize genotypes. Moreover, a slightly greater variability in phenotypic responses among genotypes under control conditions (Fig. S8) might have further contributed to the greater number of established associations.

As we previously observed under nitrogen limitation (Giuliano et al. 2026), significant correlations among root anatomical and architectural phenotypes and relative abundances of prokaryotic taxa were more numerous in the lower-performing than in the well-performing genotypes under drought, while the opposite was observed under control conditions. Even though this study does not aim to and cannot investigate the causal links between root phenotype-microbiome associations and plant performance groups, we believe that it is important to discuss the significant correlations found in the lower-performing genotypes under drought to hypothesize possible association mechanisms establishing under stress conditions. In this study, number of metaxylem vessels (No. MX) more often negatively correlated with non-pathogenic prokaryotic genera but also positively with potentially beneficial bacteria under stress (e.g., *Deinococcus*, *Aridibacter*). Therefore, an increase in number of metaxylem vessels, which can be adaptive under drought (Klein et al. 2020), could be beneficial for the associations with certain taxa but not with others. We also observed a greater number of negative correlations between root diameter (Diameter mean, max) and prokaryotic taxa abundances. Indeed, thicker nodal root diameter can improve soil exploration under drought, but it is hypothesized to decrease interactions with nitrogen-cycling microbes due to lower available surface area (Lynch et al. 2023). Finer roots, instead, can lower transpiration rates and improve water absorption through a higher surface area relative to the volume (Klein et al. 2020; Yan et al. 2022). Plants expressing these phenotypes might also display a better carbon economy, allowing for greater carbon rhizodeposition which facilitates microbial associations (Wang et al. 2021; Galindo-Castañeda et al. 2024). Therefore, both finer roots and a smaller number of metaxylem vessels can be of interest for further investigation under drought stress.

### Cortical parenchyma wall width might be a key anatomical phenotype for microbial associations

In this study, we discovered that the *cortical parenchyma wall width (CPW)* explained 13.1% of the variance of prokaryotic β-diversity under control conditions in a subset of six maize genotypes. Previous studies have focused on the thickening of cell walls in different cell types, such as endodermal and hypodermal cells and the outer cortical cells, as a useful trait to sustain the structure of tissues and improve adaptation to different stresses, such as salinity or soil compaction (Degenhardt and Gimmler 2000; Schneider et al. 2021). Sidhu et al. (2024) proposed that increased cortical parenchyma wall width can decrease metabolic costs of soil exploration, especially under terminal drought, possibly due to reduced cytosol:wall volume ratio in the cortex as suggested by RootSlice simulations. Moreover, using the same dataset for the six genotypes employed in the present study, along with additional data, Sidhu et al. (2024) reported a positive correlation between CPW and yield with overall improved drought resistance in the field.

We hypothesize that the level of expression of cortical parenchyma wall width can influence root exudation and the apoplastic space available for microbial colonization, for example for arbuscular mycorrhizal fungi. Also, the positive correlation of this phenotype with two non-pathogenic and potentially beneficial or N-fixing microbial taxa (i.e., *Saitozyma* and *Azospirillum*) (Cassán et al. 2020; Ramos-Garza et al. 2023) is promising. However, CPW studies should be complemented by measurements of the cytosolic space and by the empirical quantification of the CPW effect on root exudation rate, and therefore on microbial assemblies, for example by employing genotypes expressing contrasting levels of CPW or developing plant mutants defective in producing thick parenchyma cell walls. Furthermore, the relative contribution of this phenotype and of the associated microbiomes to plant yield should be further explored with manipulative experiments. Lastly, having investigated six genotypes with known contrasting expression of CPW might have contributed to the greater effect of CPW observed on prokaryotic β-diversity compared to other phenotypes investigated across 22 genotypes. This observation supports the importance of selecting genotypes based on specific root phenotypes rather than only on yield capability (Lynch et al. 2023).

### Yield-based plant performance associated with differences in root anatomy and rhizosphere microbiomes

In this study, most root anatomical and architectural phenotypes did not differentiate between the performance groups identified based on relative yield change (Fig. S5, S8, S12, S13, Notes S2). Only the well-performing IBM313 displayed higher cortical parenchyma wall width (CPW) under drought compared to control conditions (Fig. 1c) and aerenchyma was the only phenotype significantly increasing in the lower-performing genotypes no matter the treatment (Fig. S12). Aerenchyma is an adaptive trait induced by drought stress (Zhu et al. 2010). Therefore, genotypes experiencing greater stress overall (i.e., lower-performing genotypes) might have had greater aerenchyma expression then the well-performing genotypes in this study. These results indicate that root phenotypes alone do not explain yield performance in this study, but that other factors, such as rhizosphere microbes might play a role. Indeed, we observed a separation between prokaryotic and fungal community structures based on the performance groups (Fig. S14, Notes S2), which support the importance of understanding rhizosphere microbial associations under drought.

Nevertheless, considering all 22 genotypes, root cortex- and stele related phenotypes significantly explained 2-7% of the variance of yield under drought. This result agrees with previous studies highlighting the influence of these same anatomical phenotypes, such as stele and living cortical area, and number of metaxylem vessels, on maize yield under drought (Zhu et al. 2010; Chimungu et al. 2014b; Klein et al. 2020). Therefore, investigating the association of a complex array of root anatomical and architectural phenotypes with rhizosphere microbiomes is challenging and requires finetuning, especially when it comes to quantifying the relative contribution of root phenotypes and microbiomes to plant performance.

## Conclusion

Our results show that root anatomical phenotypes are more predominantly associated with rhizosphere microbiomes at community and taxon levels than architectural phenotypes across 22 maize inbred lines grown in the field with and without drought stress. Specifically, cortical parenchyma wall width, number of metaxylem vessels and cell files, and root diameter explained a small but significant percentage of the variance of the prokaryotic and fungal β-diversity under control conditions but not under drought. Moreover, we found a greater number of correlations between root phenotypes and microbial taxa relative abundances for anatomical phenotypes (n=248) compared to architectural phenotypes (n=58) and, overall, under optimal water availability. We identified promising correlations between potentially beneficial microbial taxa and cortex-related anatomical phenotypes, such as number of cell files, living cortical area and cortical parenchyma wall width which can be investigated further in manipulative experiments.

We are aware that the observed moderate effects of root phenotypes on microbial diversity based on statistical tests and correlations might not necessarily translate into a strong impact at the ecological level. Without in depth knowledge of the functional significance of these associations and due to the complexity of the root-microbe-soil system, it is still challenging to assign ecological relevance. However, these associations are a starting point for future studies exploring the potential to modify a limited but significant portion of the microbial diversity at a finer scale through differential root phenotypic expression under specific environmental conditions. For example, a consistent and potentially beneficial change in the rhizosphere microbial diversity associated with a specific level of expression of a certain root phenotype could be harnessed through repeated selection cycles through plant breeding programs (Vieira et al. 2025). Moreover, the fact that more associations between root anatomy and microbial diversity were found in control rather than drought conditions might suggest that adaptive root phenotypes do not necessarily play a role in microbial associations under stress. Also, more studies like the one presented here could be pursued using a wider range of genotypes, for example coming from maize diversity panels, to reveal the extent to which root phenotypes and microbial associations are genetically controlled. So far, quantitative genetics of plant-microbe associations has not fully integrated root anatomy and architecture. However, taking our results into account, interesting associations might be discovered if root anatomy, and especially cortex-related phenotypes, would be considered in such studies. Our findings highlight the need for future studies focusing on understanding the mechanism behind the observed associations between root anatomy and rhizosphere microbial diversity and their implication for sustainable agriculture.

## Data availability

The plant and soil data and the R scripts are available on Zenodo (https://doi.org/10.5281/zenodo.20511548). The DNA amplicon sequencing data were deposited in the European Nucleotide Archive (ENA, accession number PRJEB104640).

## Acknowledgments

We thank the support of Molly Hanlon and the field manager (Austin Kirt and his team) of the Rock Spring research facility, and Dr. Shawn Kaeppler (University of Wisconsin) for supplying the IBM genotypes. We are grateful to Connor Hoffman, Spencer Riccio, Samuel Walker, Douglas Watford, and Michael Frantz for assisting with the harvest and the sample collection, to Gabriella Cavell and Rebecca Groff for producing additional root-cross section pictures via laser ablation and to Argeo Ulrich and Alesia Boiko for helping with the *RootScan* picture processing. We are thankful to Britta Jahn-Humphrey for the assistance in the lab, Lian Tengxiang, Moritz Bach, Cong Zhang and Manon Longepierre for helping with the rhizosphere soil processing, to Maria Domenica Moccia for the sequencing at the Functional Genomic Center Zurich (FGCZ) and to Alexandre Strigens for the input during the start of the project. DeepL was used to improve the structure of some sentences. Microsoft Copilot was used to support R-based data handling, statistical analyses and partial literature searching, and to check for the applicability and interpretation of certain analyses.

## Author Contributions

Elena Giuliano, Tania Galindo-Castañeda, Martin Hartmann, Jonathan P. Lynch and Jagdeep Singh Sidhu designed the study. Elena Giuliano collected root, rhizosphere, and bulk soil samples with the support of Jagdeep Singh Sidhu and Ivan Lopez-Valdivia; she ablated the root samples and processed the pictures with *RootScan*, performed physiochemical analyses and DNA metabarcoding on the soil samples, analyzed and interpreted the data and wrote the manuscript. Martin Hartmann, Tania Galindo-Castañeda and Jonathan P. Lynch supported the method development and the data analysis and interpretation. With the support of Jonathan P. Lynch, Jagdeep Singh Sidhu designed the field experiment and managed and harvested it together with Ivan Lopez-Valdivia. Jagdeep Singh Sidhu handled the plant biomass, yield and root architecture data collection. Cody DePew organized the laser ablation of additional root pictures. Rafaela Feola Conz supported the soil sample processing, the molecular lab work, and analyzed part of the root pictures with *RootScan*. Johan Six and Martin Hartmann provided lab spaces and materials. All authors gave feedback and approved to the manuscript.

## Status of the manuscript

this manuscript and the accompanying supplementary information constitute the updated version of the second chapter of Elena Giuliano’s PhD thesis titled “Unearthing synergies: associations between root phenotypes and rhizosphere microbiomes in maize under resource limitation” (ETH Zurich Research Collection 2026, https://doi.org/10.3929/ethz-c-000799814). This preprint version incorporates major revisions following peer review, as this manuscript is currently under review at the journal Plant and Soil.

## References

Agarwal P, Vibhandik R, Agrahari R, et al (2024) Role of Root Exudates on the Soil Microbial Diversity and Biogeochemistry of Heavy Metals. Appl Biochem Biotechnol 196:2673–2693. 10.1007/S12010-023-04465-2

Ahmad HM, Fiaz S, Hafeez S, et al (2022) Plant Growth-Promoting Rhizobacteria Eliminate the Effect of Drought Stress in Plants: A Review. Front Plant Sci 13:875774. 10.3389/fpls.2022.875774

Ali N, Abbas SAAA, Sharif L, et al (2024) Microbial extracellular polymeric substance and impacts on soil aggregation. Bacterial Secondary Metabolites: Synthesis and Applications in Agroecosystem 221–237. 10.1016/B978-0-323-95251-4.00021-1

Allakonon MGB, Akponikpè PBI (2022) Relationship of Maize Yield to Climatic and Environmental Factors under Deficit Irrigation: A Quantitative Review. International Journal of Agronomy 2022:2408439. 10.1155/2022/2408439

Anderson MJ (2001) A new method for non-parametric multivariate analysis of variance. Austral Ecol 26:32–46. 10.1111/J.1442-9993.2001.01070.PP.X

Anderson MJ (2006) Distance-Based Tests for Homogeneity of Multivariate Dispersions. Biometrics 62:245–253. 10.1111/J.1541-0420.2005.00440.X

Anderson MJ, Ellingsen KE, McArdle BH (2006) Multivariate dispersion as a measure of beta diversity. Ecol Lett 9:683–693. 10.1111/J.1461-0248.2006.00926.X

Antoun R, Beauchamp CJ, Goussard N, et al (1998) Potential of Rhizobium and Bradyrhizobium species as plant growth promoting rhizobacteria on non-legumes: Effect on radishes (Raphanus sativus L.). Molecular Microbial Ecology of the Soil 204:57–67. 10.1007/978-94-017-2321-3_5

Ayiti OE, Ayangbenro AS, Babalola OO (2022) 16S Amplicon Sequencing of Nitrifying Bacteria and Archaea Inhabiting Maize Rhizosphere and the Influencing Environmental Factors. Agriculture (Switzerland) 12:1328. 10.3390/AGRICULTURE12091328/S1

Bengtsson-Palme J, Hartmann M, Eriksson KM, et al (2015) metaxa2: improved identification and taxonomic classification of small and large subunit rRNA in metagenomic data. Mol Ecol Resour 15:1403–1414. 10.1111/1755-0998.12399

Bengtsson-Palme J, Ryberg M, Hartmann M, et al (2013) Improved software detection and extraction of ITS1 and ITS2 from ribosomal ITS sequences of fungi and other eukaryotes for analysis of environmental sequencing data. Methods Ecol Evol 4:914–919. 10.1111/2041-210X.12073

Benjamini Y, Hochberg Y (1995) Controlling the False Discovery Rate: A Practical and Powerful Approach to Multiple Testing. Journal of the royal statistical society. Series B (Methodological) 57: 289–300. 10.1111/j.2517-6161.1995.tb02031.x

Boubekri K, Soumare A, Mardad I, et al (2022) Multifunctional role of Actinobacteria in agricultural production sustainability: A review. Microbiol Res 261:127059. 10.1016/J.MICRES.2022.127059

Brisson VL, Schmidt JE, Northen TR, et al (2019) Impacts of Maize Domestication and Breeding on Rhizosphere Microbial Community Recruitment from a Nutrient Depleted Agricultural Soil. Scientific Reports 2019 9:1 9:1–14. 10.1038/s41598-019-52148-y

Burton AL, Lynch JP, Brown KM (2013) Spatial distribution and phenotypic variation in root cortical aerenchyma of maize (*Zea mays* L.). Plant Soil 367:263–274. http://www.jstor.org/stable/42952892

Burton AL, Williams M, Lynch JP, Brown KM (2012) RootScan: Software for high-throughput analysis of root anatomical traits. Plant Soil 357:189–203. 10.1007/s11104-012-1138-2

Cassán F, Coniglio A, López G, et al (2020) Everything you must know about *Azospirillum* and its impact on agriculture and beyond. Biology and Fertility of Soils 2020 56:4 56:461–479. 10.1007/S00374-020-01463-Y

Chaudhary S, Shankar A, Singh A, Prasad V (2018) Usefulness of *Penicillium* in Enhancing Plants Resistance to Abiotic Stresses: An Overview. New and Future Developments in Microbial Biotechnology and Bioengineering: Penicillium System Properties and Applications 277–284. 10.1016/B978-0-444-63501-3.00017-X

Chimungu JG, Brown KM, Lynch JP (2014a) Large root cortical cell size improves drought tolerance in maize. Plant Physiol 166:2166–2178. 10.1104/pp.114.250449

Chimungu JG, Brown KM, Lynch JP (2014b) Reduced root cortical cell file number improves drought tolerance in maize. Plant Physiol 166:1943–1955. 10.1104/pp.114.249037

Chimungu JG, Maliro MFA, Nalivata PC, et al (2015) Utility of root cortical aerenchyma under water limited conditions in tropical maize (*Zea mays* L.). Field Crops Res 171:86–98. 10.1016/J.FCR.2014.10.009

Corlett M (1981) A taxonomic survey of some species of *Didymella* and *Didymella*-like species. Can J Bot 59, 2016–2042. 10.1139/b81-264

Costa OYA, Raaijmakers JM, Kuramae EE (2018) Microbial extracellular polymeric substances: Ecological function and impact on soil aggregation. Front Microbiol 9:337094. 10.3389/fmicb.2018.01636

Degenhardt B, Gimmler H (2000) Cell wall adaptations to multiple environmental stresses in maize roots. J Exp Bot 51:595–603. 10.1093/JEXBOT/51.344.595

Dikilitas M, Karakas S, Hashem A, et al (2016) Oxidative stress and plant responses to pathogens under drought conditions. Water Stress and Crop Plants: A Sustainable Approach 1–2:102– 123. 10.1002/9781119054450.ch8

Dreyer B, Morte A, López JÁ, Honrubia M (2009) Comparative study of mycorrhizal susceptibility and anatomy of four palm species. Mycorrhiza 2009 20:2 20:103–115. 10.1007/S00572-009-0266-X

Ebrahimi-Zarandi M, Etesami H, Glick BR (2023) Fostering plant resilience to drought with Actinobacteria: Unveiling perennial allies in drought stress tolerance. Plant Stress 10:100242. 10.1016/J.STRESS.2023.100242

Edenhofer Ottmar (2015) Climate change 2014 : mitigation of climate change: Working Group III contribution to the Fifth assessment report of the Intergovernmental Panel on Climate Change. Cambridge University Press.

Edgar RC (2016a) UNOISE2: improved error-correction for Illumina 16S and ITS amplicon sequencing. bioRxiv 081257. 10.1101/081257

Edgar RC (2016b) UCHIME2: improved chimera prediction for amplicon sequencing. bioRxiv 074252. 10.1101/074252

Edgar RC (2016c) SINTAX: a simple non-Bayesian taxonomy classifier for 16S and ITS sequences. bioRxiv 074161. 10.1101/074161

Edgar RC, Flyvbjerg H (2015) Error filtering, pair assembly and error correction for next-generation sequencing reads. Bioinformatics 31:3476–3482. 10.1093/BIOINFORMATICS/BTV401

Etesami H, Chen Y (2025) Adapting to arid conditions: the interplay of plant roots, microbial communities, and exudates in the face of drought challenges. Sustainable Agriculture under Drought Stress: Integrated Soil, Water and Nutrient Management 471–487. 10.1016/B978-0-443-23956-4.00028-4

Eziz A, Yan Z, Tian D, et al (2017) Drought effect on plant biomass allocation: A meta-analysis. Ecol Evol 7:11002–11010. 10.1002/ece3.3630

Fan W, Tang F, Wang J, et al (2023) Drought-induced recruitment of specific root-associated bacteria enhances adaptation of alfalfa to drought stress. Front Microbiol 14:1114400. 10.3389/fmicb.2023.1114400

Farré I, Faci JM (2009) Deficit irrigation in maize for reducing agricultural water use in a Mediterranean environment. Agric Water Manag 96:383–394. 10.1016/J.AGWAT.2008.07.002

Favela A, O. Bohn M, D. Kent A (2021) Maize germplasm chronosequence shows crop breeding history impacts recruitment of the rhizosphere microbiome. ISME J 15:2454–2464. 10.1038/S41396-021-00923-Z

Freschet GT, Pagès L, Iversen CM, et al (2021) A starting guide to root ecology: strengthening ecological concepts and standardising root classification, sampling, processing and trait measurements. New Phytologist 232:973–1122. 10.1111/nph.17572

Frey B, Rime T, Phillips M, et al (2016) Microbial diversity in European alpine permafrost and active layers. FEMS Microbiol Ecol 92:18. 10.1093/FEMSEC/FIW018

Galindo-Castañeda T, Brown KM, Kuldau GA, et al (2019) Root cortical anatomy is associated with differential pathogenic and symbiotic fungal colonization in maize. Plant Cell Environ 42:2999–3014. 10.1111/PCE.13615

Galindo-Castañeda T, Brown KM, Lynch JP (2018) Reduced root cortical burden improves growth and grain yield under low phosphorus availability in maize. Plant Cell Environ 41:1579–1592. 10.1111/PCE.13197

Galindo-Castañeda T, Hartmann M, Lynch JP (2024) Location: root architecture structures rhizosphere microbial associations. J Exp Bot 75:594–604. 10.1093/JXB/ERAD421

Galindo-Castañeda T, Kost E, Giuliano E, et al (2025) Locating the microbes along the maize root system under nitrogen limitation: a root phenotypic approach. Ann Bot. 10.1093/AOB/MCAF185

Galindo-Castañeda T, Lynch JP, Six J, Hartmann M (2022) Improving Soil Resource Uptake by Plants Through Capitalizing on Synergies Between Root Architecture and Anatomy and Root-Associated Microorganisms. Front Plant Sci 13:827369. 10.3389/fpls.2022.827369

Galindo-Castañeda T, Rojas C, Karaöz U, et al (2023) Influence of root cortical aerenchyma on the rhizosphere microbiome of field-grown maize. bioRxiv 2023.01.31.525837. 10.1101/2023.01.31.525837

Gao Y, Lynch JP (2016) Reduced crown root number improves water acquisition under water deficit stress in maize (*Zea mays* L.). J Exp Bot 67:4545–4557. 10.1093/JXB/ERW243

Gao Y, Zhao Y, Li P, Qi X (2023) Responses of the maize rhizosphere soil environment to drought-flood abrupt alternation stress. Front Microbiol 14:1295376. 10.3389/fmicb.2023.1295376

Giuliano E, Sidhu SJ, Lopez-Valdivia I, et al (2026) Shovelomics meets microbiomics: root phenotype-microbiome associations and links with maize yield under nitrogen limitation. Rhizosphere 101241. 10.1016/J.RHISPH.2025.101241

Halo BA, Aljabri YAS, Yaish MW (2025) Drought-induced microbial dynamics in cowpea rhizosphere: Exploring bacterial diversity and bioinoculant prospects. PLoS One 20:e0320197. 10.1371/JOURNAL.PONE.0320197

Hartmann M, Six J (2022) Soil structure and microbiome functions in agroecosystems. Nature Reviews Earth & Environment 2022 4:1 4:4–18. 10.1038/s43017-022-00366-w

Hartwig RP, Santangeli M, Würsig H, et al (2025) Drought response of the maize plant–soil–microbiome system is influenced by plant size and presence of root hairs. Ann Bot 1–18. 10.1093/AOB/MCAF033

He X, Wang D, Jiang Y, et al (2024) Heritable microbiome variation is correlated with source environment in locally adapted maize varieties. Nature Plants 2024 10:4 10:598–617. 10.1038/s41477-024-01654-7

Hochholdinger F (2009) The Maize Root System: Morphology, Anatomy, and Genetics. Handbook of Maize: Its Biology 145–160. 10.1007/978-0-387-79418-1_8

Oksanen J, Simpson G, Blanchet F, et al (2024). vegan: Community Ecology Package - R package version 2.6-8. https://CRAN.R-project.org/package=vegan.

Pinheiro J, Bates D, DebRoy S, at al (2022) Package ‘nlme’. Linear and nonlinear mixed effects models v3.1-160 (version 3.1-160). https://svn.r-project.org/R-packages/trunk/nlme/

Jochum MD, McWilliams KL, Borrego EJ, et al (2019) Bioprospecting Plant Growth-Promoting Rhizobacteria That Mitigate Drought Stress in Grasses. Front Microbiol 10:466447. 10.3389/fmicb.2019.02106

Kihara S, Yamamoto K, Shiwa Y, et al (2024) The rhizosphere bacterial community of water yam (*Dioscorea alata* L.) under limited water conditions. Journal of Sustainable Agriculture and Environment 3:e70009. 10.1002/sae2.70009

Klein SP, Kaeppler SM, Brown KM, Lynch JP (2024) Integrating GWAS with a gene co-expression network better prioritizes candidate genes associated with root metaxylem phenes in maize. Plant Genome 17:e20489. 10.1002/tpg2.20489

Klein SP, Schneider HM, Perkins AC, et al (2020) Multiple Integrated Root Phenotypes Are Associated with Improved Drought Tolerance. Plant Physiol 183:1011–1025. 10.1104/PP.20.00211

Lammerts van Bueren ET, Struik PC (2017) Diverse concepts of breeding for nitrogen use efficiency. A review. Agronomy for Sustainable Development 2017 37:5 37:1–24. 10.1007/S13593-017-0457-3

Langmead B, Salzberg SL (2012) Fast gapped-read alignment with Bowtie 2. Nature Methods 2012 9:4 9:357–359. 10.1038/nmeth.1923

Lattacher A, Le Gall S, Rothfuss Y, et al (2025) Rooting for microbes: impact of root architecture on the microbial community and function in top- and subsoil. Plant Soil 513:333–351. 10.1007/s11104-024-07181-w

Lee M, Sharopova N, Beavis WD, et al (2002) Expanding the genetic map of maize with the intermated B73 x Mo17 (IBM) population. Plant Mol Biol 48:453–461. 10.1023/A:1014893521186

Leng G, Hall J (2019) Crop yield sensitivity of global major agricultural countries to droughts and the projected changes in the future. Science of the Total Environment 654:811–821. 10.1016/j.scitotenv.2018.10.434

Lesk C, Rowhani P, Ramankutty N (2016) Influence of extreme weather disasters on global crop production. Nature 2016 529:7584 529:84–87. 10.1038/nature16467

Li J, Zhou L, Chen G, et al (2025) Arbuscular mycorrhizal fungi enhance drought resistance and alter microbial communities in maize rhizosphere soil. Environ Technol Innov 37:103947. 10.1016/J.ETI.2024.103947

Li Y, An J, Guo J, et al (2024) Ridge planting increases the rhizosphere microbiome diversity and improves the yield of *Pinellia ternata* (Thunb.) Breit in North China. PLoS One 19:e0304898. 10.1371/JOURNAL.PONE.0304898

Li H, Xie J, Gao Y, et al (2023) IQ domain-containing protein ZmIQD27 modulates water transport in maize. Plant Physiol 193:1834–1848. 10.1093/PLPHYS/KIAD390

Li P, Tedersoo L, Crowther TW, et al (2023). Fossil-fuel-dependent scenarios could lead to a significant decline of global plant-beneficial bacteria abundance in soils by 2100. Nat Food 4, 996–1006. 10.1038/s43016-023-00869-9

Li Y, Qu Z, Xu W, et al (2022) Maize (*Zea mays* L.) genotypes induce the changes of rhizosphere microbial communities. Archives of Microbiology 2022 204:6 204:1–13. 10.1007/S00203-022-02934-6

Liu Y, Ren J, Yu B, et al (2025) Metagenomic and Metabolomic Perspectives on the Drought Tolerance of Broomcorn Millet (*Panicum miliaceum* L.). Microorganisms 13:1593. 10.3390/MICROORGANISMS13071593

Loiko N, Islam MN (2024) Plant–Soil Microbial Interaction: Differential Adaptations of Beneficial vs. Pathogenic Bacterial and Fungal Communities to Climate-Induced Drought. Agronomy 2024, Vol 14, Page 1949 14:1949. 10.3390/AGRONOMY14091949

Longepierre M, Widmer F, Keller T, et al (2021) Limited resilience of the soil microbiome to mechanical compaction within four growing seasons of agricultural management. ISME Communications 2021 1:1–13. 10.1038/s43705-021-00046-8

Lundberg DS, Lebeis SL, Paredes SH, et al (2012) Defining the core Arabidopsis thaliana root microbiome. Nature 488:86–90. 10.1038/NATURE11237

Luo Y, Zhou M, Zhao Q, et al (2020) Complete genome sequence of *Sphingomonas* sp. Cra20, a drought resistant and plant growth promoting rhizobacteria. Genomics 112:3648–3657. 10.1016/J.YGENO.2020.04.013

Lynch JP (2022) Harnessing root architecture to address global challenges. The Plant Journal 109:415–431. 10.1111/TPJ.15560

Lynch JP (2013) Steep, cheap and deep: an ideotype to optimize water and N acquisition by maize root systems. Ann Bot 112:347–357. 10.1093/AOB/MCS293

Lynch JP, Galindo-Castañeda T, Schneider HM, et al (2023) Root phenotypes for improved nitrogen capture. Plant and Soil 2023 502:1 502:31–85. 10.1007/S11104-023-06301-2

Lynch JP, Mooney SJ, Strock CF, Schneider HM (2022) Future roots for future soils. Plant Cell Environ 45:620–636. 10.1111/pce.14213

Lynch JP, Strock CF, Schneider HM, et al (2021) Root anatomy and soil resource capture. Plant and Soil 2021 466:1 466:21–63. 10.1007/S11104-021-05010-Y

Ma J, Liu D, Zhao P, et al (2024) Intercropping of tobacco and maize at seedling stage promotes crop growth through manipulating rhizosphere microenvironment. Front Plant Sci 15:1470229. 10.3389/fpls.2024.1470229

Maechler M Rousseeuw P, Struyf A, et al (2025) Cluster: Cluster Analysis Basics and Extensions. R package version 2.1.8.1. — For new features, see the “NEWS” and the “Changelog” file in the package source. https://CRAN.R-project.org/package=cluster

Maia SMF, Galvão C de O, Abbruzzini TF, et al (2024) Managing Drought Stress in Agro-Ecosystems of Latin America and the Caribbean Region. In Book: Soil and Drought: Basic Processes 157–180. 10.1201/b22954-7

Marasco R, Fusi M, Rolli E, et al (2021) Aridity modulates belowground bacterial community dynamics in olive tree. Environ Microbiol 23:6275–6291. 10.1111/1462-2920.15764

Martin M (2011) Cutadapt removes adapter sequences from high-throughput sequencing reads. EMBnet J 17:10–12. 10.14806/EJ.17.1.200

Martins BR, Radl V, Treder K, et al (2024) The rhizosphere microbiome of 51 potato cultivars with diverse plant growth characteristics. FEMS Microbiol Ecol 100:88. 10.1093/FEMSEC/FIAE088

McPherson MR, Wang P, Marsh EL, et al (2018) Isolation and analysis of microbial communities in soil, rhizosphere, and roots in perennial grass experiments. Journal of Visualized Experiments. 10.3791/57932

Meier MA, Xu G, Lopez-Guerrero MG, et al (2022) Association analyses of host genetics, root-colonizing microbes, and plant phenotypes under different nitrogen conditions in maize. Elife 11. 10.7554/ELIFE.75790

Meisner A, Jacquiod S, Snoek BL, et al (2018) Drought legacy effects on the composition of soil fungal and prokaryote communities. Front Microbiol 9:323518. 10.3389/fmicb.2018.00294

Mukhtar H, Hao J, Xu G, et al (2024) Nitrogen input differentially shapes the rhizosphere microbiome diversity and composition across diverse maize lines. Biol Fertil Soils 61:1–12. 10.1007/s00374-024-01863-4

Narsing Rao MP, Lohmaneeratana K, Bunyoo C, Thamchaipenet A (2022) Actinobacteria–Plant Interactions in Alleviating Abiotic Stress. Plants 11:2976. 10.3390/PLANTS11212976

Nilsson RH, Larsson KH, Taylor AFS, et al (2019) The UNITE database for molecular identification of fungi: handling dark taxa and parallel taxonomic classifications. Nucleic Acids Res 47:D259–D264. 10.1093/NAR/GKY1022

Ozimek E, Hanaka A (2021) *Mortierella* Species as the Plant Growth-Promoting Fungi Present in the Agricultural Soils. Agriculture 11:7. 10.3390/AGRICULTURE11010007

Parasar BJ, Sharma I, Agarwala N (2024) Root exudation drives abiotic stress tolerance in plants by recruiting beneficial microbes. Applied Soil Ecology 198:105351. 10.1016/J.APSOIL.2024.105351

Peiffer JA, Spor A, Koren O, et al (2013) Diversity and heritability of the maize rhizosphere microbiome under field conditions. Proc Natl Acad Sci U S A 110:6548–6553. 10.1073/pnas.1302837110

Pérez-Jaramillo JE, Carrión VJ, Bosse M, et al (2017) Linking rhizosphere microbiome composition of wild and domesticated Phaseolus vulgaris to genotypic and root phenotypic traits. The ISME Journal 2017 11:10 11:2244–2257. 10.1038/ismej.2017.85

Pignède E (2025) Who carries the burden of climate change? Heterogeneous impact of droughts in sub-Saharan Africa. Am J Agric Econ 107:925–957. 10.1111/AJAE.12507

Preece C, Verbruggen E, Liu L, et al (2019) Effects of past and current drought on the composition and diversity of soil microbial communities. Soil Biol Biochem 131:28–39. 10.1016/J.SOILBIO.2018.12.022

Quast C, Pruesse E, Yilmaz P, et al (2013) The SILVA ribosomal RNA gene database project: improved data processing and web-based tools. Nucleic Acids Res 41:D590–D596. 10.1093/NAR/GKS1219

Quattrone A, Lopez-Guerrero M, Yadav P, et al (2024) Interactions between root hairs and the soil microbial community affect the growth of maize seedlings. Plant Cell Environ 47:611–628. 10.1111/pce.14755

Quiroga G, Erice G, Aroca R, et al (2017) Enhanced drought stress tolerance by the arbuscular mycorrhizal symbiosis in a drought-sensitive maize cultivar is related to a broader and differential regulation of host plant aquaporins than in a drought-tolerant cultivar. Front Plant Sci 8:268043. 10.3389/fpls.2017.01056

Ramos-Garza J, Aguirre-Noyola JL, Bustamante-Brito R, et al (2023) Mycobiota of Mexican Maize Landraces with Auxin-Producing Yeasts That Improve Plant Growth and Root Development. Plants 12:1328. 10.3390/PLANTS12061328/S1

Rognes T, Flouri T, Nichols B, et al (2016) VSEARCH: A versatile open source tool for metagenomics. PeerJ 2016:e2584. 10.7717/PEERJ.2584/FIG-7

Rüger L, Feng K, Dumack K, et al (2021) Assembly Patterns of the Rhizosphere Microbiome Along the Longitudinal Root Axis of Maize (*Zea mays* L.). Front Microbiol 12:614501. 10.3389/fmicb.2021.614501

Rüger L, Ganther M, Freudenthal J, et al (2023) Root cap is an important determinant of rhizosphere microbiome assembly. New Phytologist 239:1434–1448. 10.1111/nph.19002

Saengwilai P, Tian X, Lynch JP (2014) Low Crown Root Number Enhances Nitrogen Acquisition from Low-Nitrogen Soils in Maize. Plant Physiol 166:581–589. 10.1104/PP.113.232603

Sánchez-Arizpe A, Luis Arispe-Vázquez J, Alejandro Cadena-Zamudio D, et al (2023) Toxigenic fungi collected in maize fields from four states of Mexico. 463 AJCS 17:1835–2707. 10.21475/ajcs.23.17.05.p3846

Schloss PD (2024) Rarefaction is currently the best approach to control for uneven sequencing effort in amplicon sequence analyses. mSphere 9. 10.1128/msphere.00354-23

Schmidt JE, Mazza Rodrigues JL, Brisson VL, et al (2020) Impacts of directed evolution and soil management legacy on the maize rhizobiome. Soil Biol Biochem 145:107794. 10.1016/J.SOILBIO.2020.107794

Schneider CA, Rasband WS, Eliceiri KW (2012) NIH Image to ImageJ: 25 years of image analysis. Nat Methods 9:671–675. 10.1038/NMETH.2089

Schneider HM, Strock CF, Hanlon MT, et al (2021) Multiseriate cortical sclerenchyma enhance root penetration in compacted soils. Proceedings of the National Academy of Sciences 118:e2012087118. 10.1073/PNAS.2012087118

Sidhu JS, Lopez-Valdivia I, Strock CF, et al (2024) Cortical parenchyma wall width regulates root metabolic cost and maize performance under suboptimal water availability. J Exp Bot 75:5750–5767. 10.1093/JXB/ERAE191

Sidhu JS, Lynch JP (2024) Cortical cell size regulates root metabolic cost. The Plant Journal 118:1343–1357. 10.1111/TPJ.16672

Sidhu JS, Schneider HM (2024) Root Anatomy: Preparing, Imaging, and Analyzing Maize Root Cross-Sections. Cold Spring Harb Protoc. 10.1101/PDB.PROT108585

Simmons T, Caddell DF, Deng S, Coleman-Derr D (2018) Exploring the root microbiome: Extracting bacterial community data from the soil, rhizosphere, and root endosphere. Journal of Visualized Experiments. 10.3791/57561

Storey JD (2002) A Direct Approach to False Discovery Rates. Source: Journal of the Royal Statistical Society Series B (Statistical Methodology) 64:479–498. 10.1111/1467-9868.00346

Strock CF, Schneider HM, Galindo-Castañeda T, et al (2019) Laser ablation tomography for visualization of root colonization by edaphic organisms. J Exp Bot 70:5327–5342. 10.1093/JXB/ERZ271

Taha AS, Fathey HA, Mohamed AH, et al (2025) Mitigating drought stress and enhancing maize resistance through biopriming with *Rhizopus arrhizus*: insights into Morpho-Biochemical and molecular adjustments. BMC Plant Biology 2025 25:1 25:1–24. 10.1186/S12870-025-06793-3

Tang H, Hassan MU, Feng L, et al (2022) The Critical Role of Arbuscular Mycorrhizal Fungi to Improve Drought Tolerance and Nitrogen Use Efficiency in Crops. Front Plant Sci 13:919166. 10.3389/fpls.2022.919166

Taylor DL, Walters WA, Lennon NJ, et al (2016) Accurate estimation of fungal diversity and abundance through improved lineage-specific primers optimized for Illumina amplicon sequencing. Appl Environ Microbiol 82:7217–7226. 10.1128/AEM.02576-16

Thomma BPHJ (2003) Alternaria spp.: from general saprophyte to specific parasite. Mol Plant Pathol 4:225–236. 10.1046/J.1364-3703.2003.00173.X

Tian L, Han M, Liang K, et al (2024) Profiling of farmland microorganisms in maize and minor-grain crops under extreme drought conditions. Applied Soil Ecology 204:105743. 10.1016/J.APSOIL.2024.105743

Tiziani R, Miras-Moreno B, Malacrinò A, et al (2022) Drought, heat, and their combination impact the root exudation patterns and rhizosphere microbiome in maize roots. Environ Exp Bot 203:105071. 10.1016/J.ENVEXPBOT.2022.105071

Trachsel S, Kaeppler SM, Brown KM, Lynch JP (2011) Shovelomics: High throughput phenotyping of maize (*Zea mays* L.) root architecture in the field. Plant Soil 341:75–87. 10.1007/s11104-010-0623-8

Vanhees DJ, Loades KW, Glyn Bengough A, et al (2020) Root anatomical traits contribute to deeper rooting of maize under compacted field conditions. J Exp Bot 71:4243–4257. 10.1093/JXB/ERAA165

Vardharajula S, Ali SZ, Grover M, et al (2011) Drought-tolerant plant growth promoting Bacillus spp.: effect on growth, osmolytes, and antioxidant status of maize under drought stress. J Plant Interact 6:1–14. 10.1080/17429145.2010.535178

Vieira RA, Nogueira APO, Fritsche-Neto R (2025) Optimizing the selection of quantitative traits in plant breeding using simulation. Front Plant Sci 16:1495662. 10.3389/fpls.2025.1495662

Vischer NOE, Verheul J, Postma M, et al (2015) Cell age dependent concentration of *Escherichia coli* divisome proteins analyzed with ImageJ and ObjectJ. Front Microbiol 6:135279. 10.3389/fmicb.2015.00586

Walters WA, Jin Z, Youngblut N, et al (2018) Large-scale replicated field study of maize rhizosphere identifies heritable microbes. Proc Natl Acad Sci U S A 115:7368–7373. 10.1073/pnas.1800918115

Wang D, He X, Baer M, et al (2024) Lateral root enriched *Massilia* associated with plant flowering in maize. Microbiome 12:1–20. 10.1186/s40168-024-01839-4

Wang R, Cavagnaro TR, Jiang Y, et al (2021) Carbon allocation to the rhizosphere is affected by drought and nitrogen addition. Journal of Ecology 109:3699–3709. 10.1111/1365-2745.13746

Wasaya A, Zhang X, Fang Q, Yan Z (2018) Root Phenotyping for Drought Tolerance: A Review. Agronomy 2018, Vol 8, Page 241 8:241. 10.3390/AGRONOMY8110241

Wen X, Lu J, Zou J, et al (2025) Maize genotypes foster distinctive bacterial and fungal communities in the rhizosphere. Agric Ecosyst Environ 382:109505. 10.1016/J.AGEE.2025.109505

Williams A, de Vries FT (2020) Plant root exudation under drought: implications for ecosystem functioning. New Phytologist 225:1899–1905. 10.1111/NPH.16223

Wu C, Zhang X, Liu Y, et al (2024) Drought Stress Increases the Complexity of the Bacterial Network in the Rhizosphere and Endosphere of Rice (*Oryza sativa* L.). Agronomy 2024, Vol 14, Page 1662 14:1662. 10.3390/AGRONOMY14081662

Xu L, Dong Z, Chiniquy D, et al (2021) Genome-resolved metagenomics reveals role of iron metabolism in drought-induced rhizosphere microbiome dynamics. Nature Communications 12:3209. 10.1038/s41467-021-23553-7

Yan M, Zhang L, Ren Y, et al (2022) The Higher Water Absorption Capacity of Small Root System Improved the Yield and Water Use Efficiency of Maize. Plants 11:2300. 10.3390/PLANTS11172300/S1

Yu P, He X, Baer M, et al (2021) Plant flavones enrich rhizosphere *Oxalobacteraceae* to improve maize performance under nitrogen deprivation. Nature Plants 2021 7:4 7:481–499. 10.1038/s41477-021-00897-y

Yu P, Wang C, Baldauf JA, et al (2018) Root type and soil phosphate determine the taxonomic landscape of colonizing fungi and the transcriptome of field-grown maize roots. New Phytologist 217:1240–1253. 10.1111/nph.14893

Zai X, Luo W, Bai W, et al (2021) Effect of Root Diameter on the Selection and Network Interactions of Root-Associated Bacterial Microbiomes in *Robinia pseudoacacia* L. Microb Ecol 82:391–402. 10.1007/s00248-020-01678-4

Zhan A, Schneider H, Lynch JP (2015) Reduced lateral root branching density improves drought tolerance in maize. Plant Physiol 168:1603–1615. 10.1104/pp.15.00187

Zhu J, Brown KM, Lynch JP (2010) Root cortical aerenchyma improves the drought tolerance of maize (*Zea mays* L.). Plant Cell Environ 33:740–749. 10.1111/j.1365-3040.2009.02099.x

Zhu J, Kaeppler SM, Lynch JP (2005) Mapping of QTL controlling root hair length in maize (*Zea mays* L.) under phosphorus deficiency. Plant Soil 270:299–310. 10.1007/s11104-004-1697-y

Zuur AF, Ieno EN, Walker N, et al (2009) Mixed effects models and extensions in ecology with R. Statistics for Biology and Health. Springer. 10.1007/978-0-387-87458-6

