## Supplementary_Information for "Digging for meaningful connections: associations between root phenotypes and rhizosphere microbial diversity in maize"

[1]Institute of Agricultural Sciences, Department of Environmental System Science, ETH Zurich, 8092 Zurich, Switzerland; [2]Department of Plant Science, The Pennsylvania State University, University Park, PA 16802, USA; [3]Division of Plant Sciences and Technology, College of Agriculture, Food and Natural Resources, University of Missouri, Columbia, MO 65201, USA; [4]Department of Physiology and Cell Biology, Leibniz Institute for Plant Genetics and Crop Plant Research (IPK), OT Gatersleben, Seeland 06466, Germany.

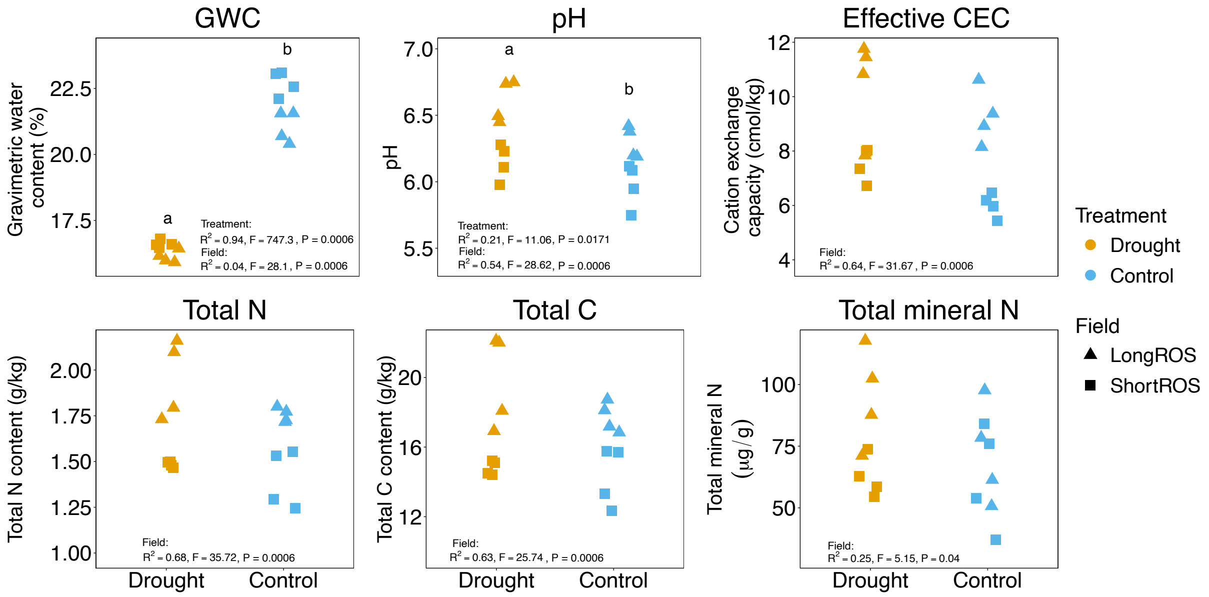


**Fig. S1 Physiochemical properties of bulk soil by treatment and field.** One sample per block in each field is reported. Gravimetric water content (GWC), pH (in water), effective cation exchange capacity (CEC), total nitrogen (N) and carbon (C) contents and total mineral N ($N\text{-}NO_3$ + $N\text{-}NH_4$) content are shown. The letters represent the significant effect of drought on GWC and pH. The PERMANOVA (permutational multivariate analysis of variance) results after FDR (false discovery rate according to Benjamini-Hochberg) p-value correction are reported in each plot. Field affected all measured parameters. After p-value correction, PERMUTEST results were not statistically significant for treatment and field.

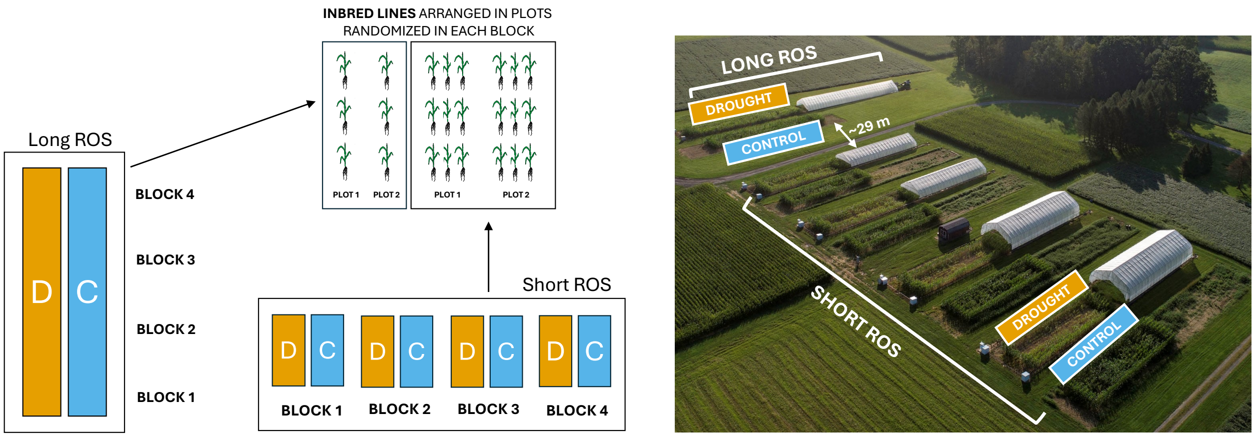


**Fig. S2 Design of the field experiment.** Split-plot design and modified split-plot design with four blocks (biological replicates) and two treatments (drought and control) were used for the Short and Long rain-out shelters (ROS), respectively. Short ROS and Long ROS were about 29 m distant from each other and were established in different years. Inbred lines (n=6) were arranged in 3-row plots in the Short ROS and 1-row plots in the Long ROS (n=16) randomized in each block. Blocks were represented by separated fields in the Short ROS and by different consecutive locations in the same field in the Long ROS. The drone picture on the right (credits: Austin Kirt) is from the 2020 field season but it represents well the 2021 field season reported in this study. The final illustration was realized with PowerPoint.

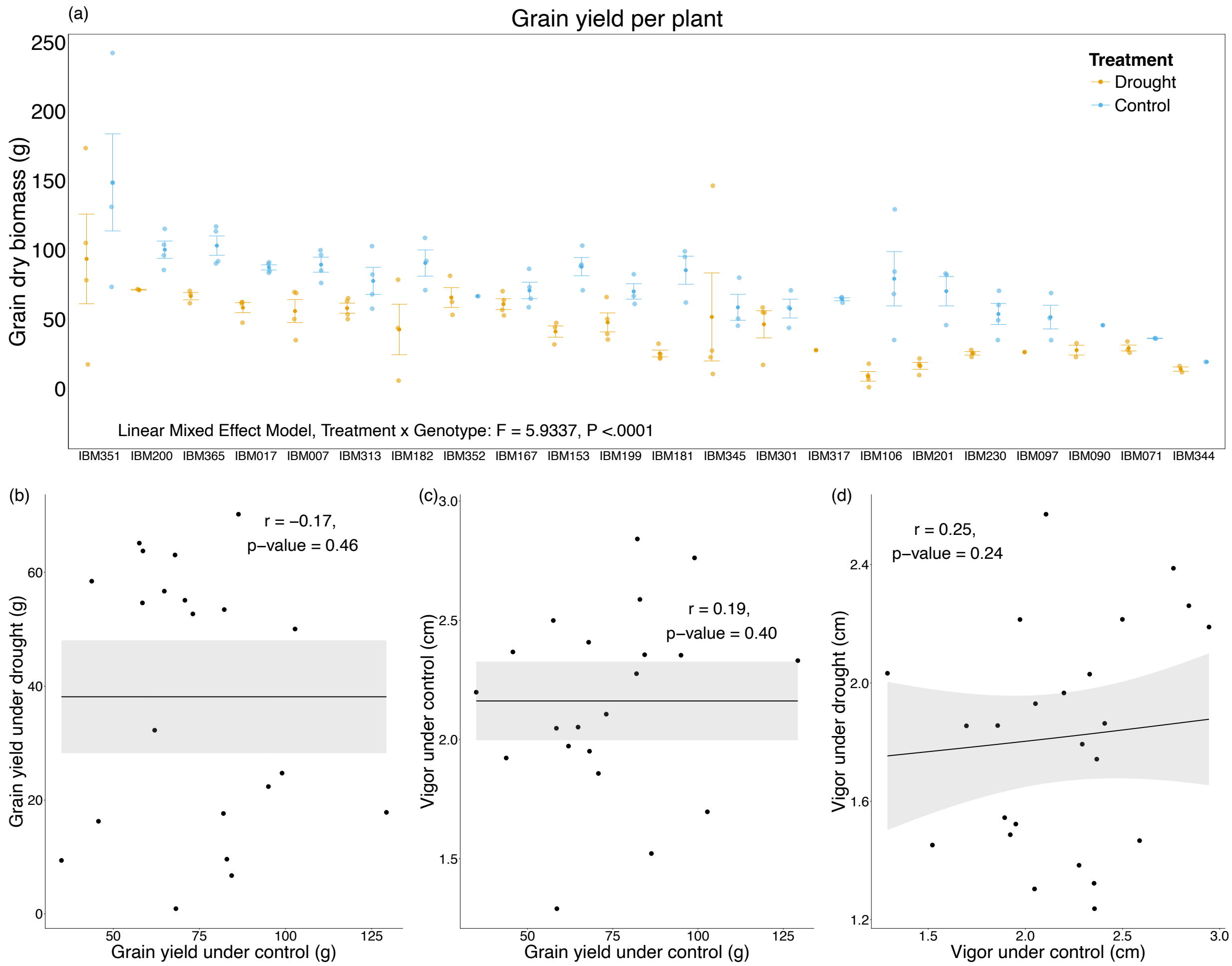


**Fig. S3 Effect of drought on grain yield and links with plant vigor.** (a) Dry grain biomass per plant (cumulative value for 3 plants divided by 3), standard error (SE) bars and reduction due to drought are shown. The statistically significant results of the interaction between drought and genotype are reported (linear mixed-effects model). The scatterplots show the absence of a significant correlation (Pearson (r), P ≥ 0.05) between grain yield under control and drought (b), grain yield and vigor under control (c) and vigor under control and drought (d) for the six genotypes included in the yield-based performance groups.

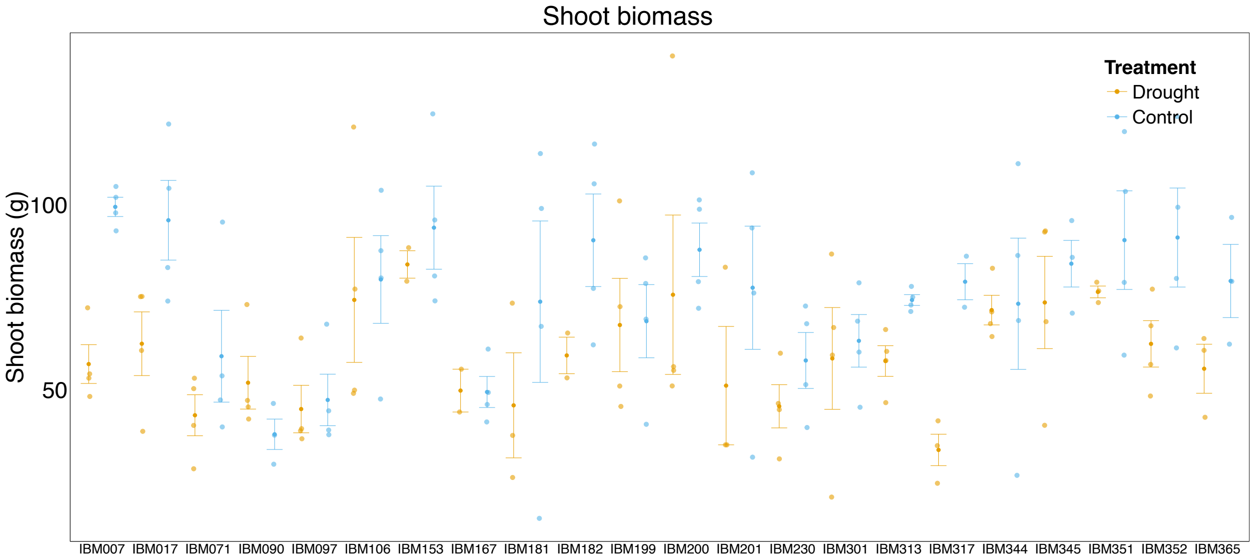


**Fig. S4 Dry shoot biomass by genotype and treatment.** Dry biomass of stems and leaves are reported for every biological replicate (data points) with the standard error (SE) bars. Eight individual samples for which leaves and cob biomass were recorded together are not reported.

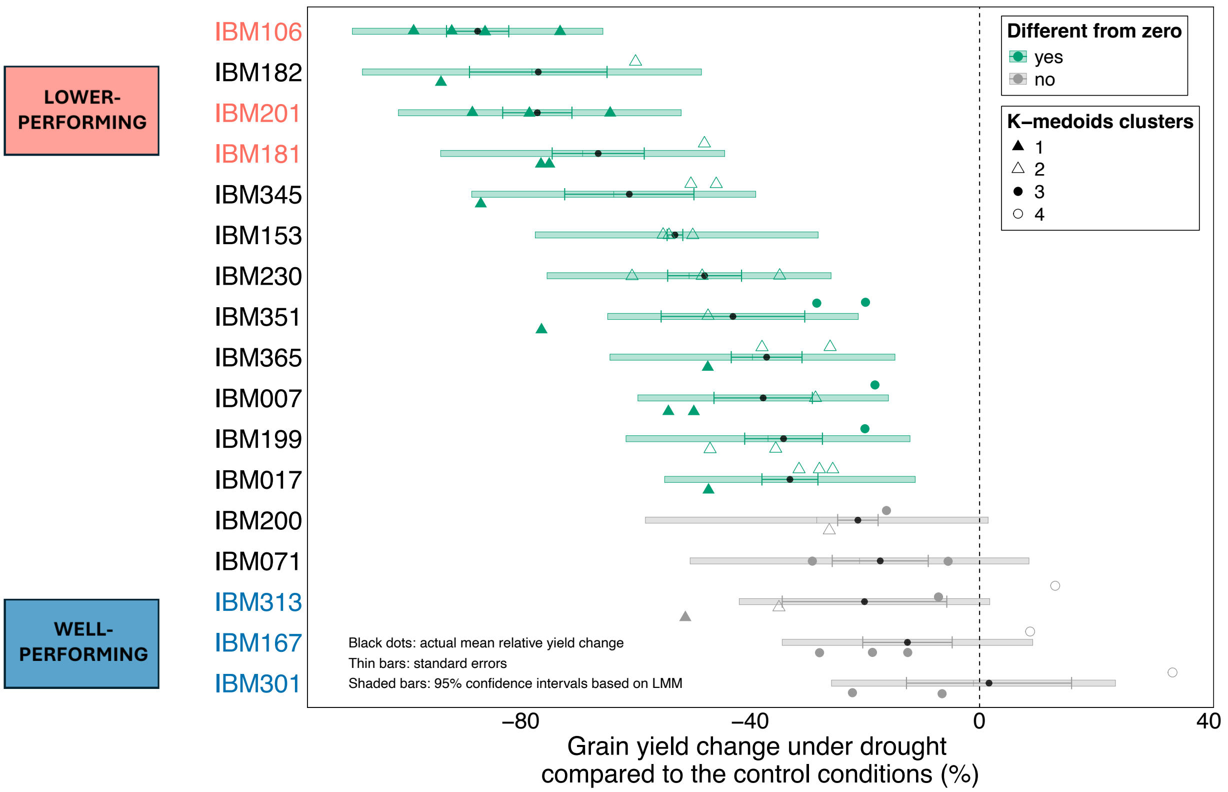


**Fig. S5 Maize inbred line responses to drought stress.** Relative change of grain yield under drought compared to control conditions (zero line) for 17 out of 22 genotypes including k-medoids clustering for each biological replicate. The statistical difference from the control conditions is based on the 95% confidence intervals (thick, shaded bars) obtained by the estimated marginal means calculated on the output of the linear mixed-effects model used to identify genotype effect on relative yield change. Data points and thin error bars represent the actual relative change data and standard errors, respectively. Groups of lower-performing (significant yield reduction under drought) and well-performing (comparable yield under drought and control conditions) genotypes are reported in red and blue, respectively.

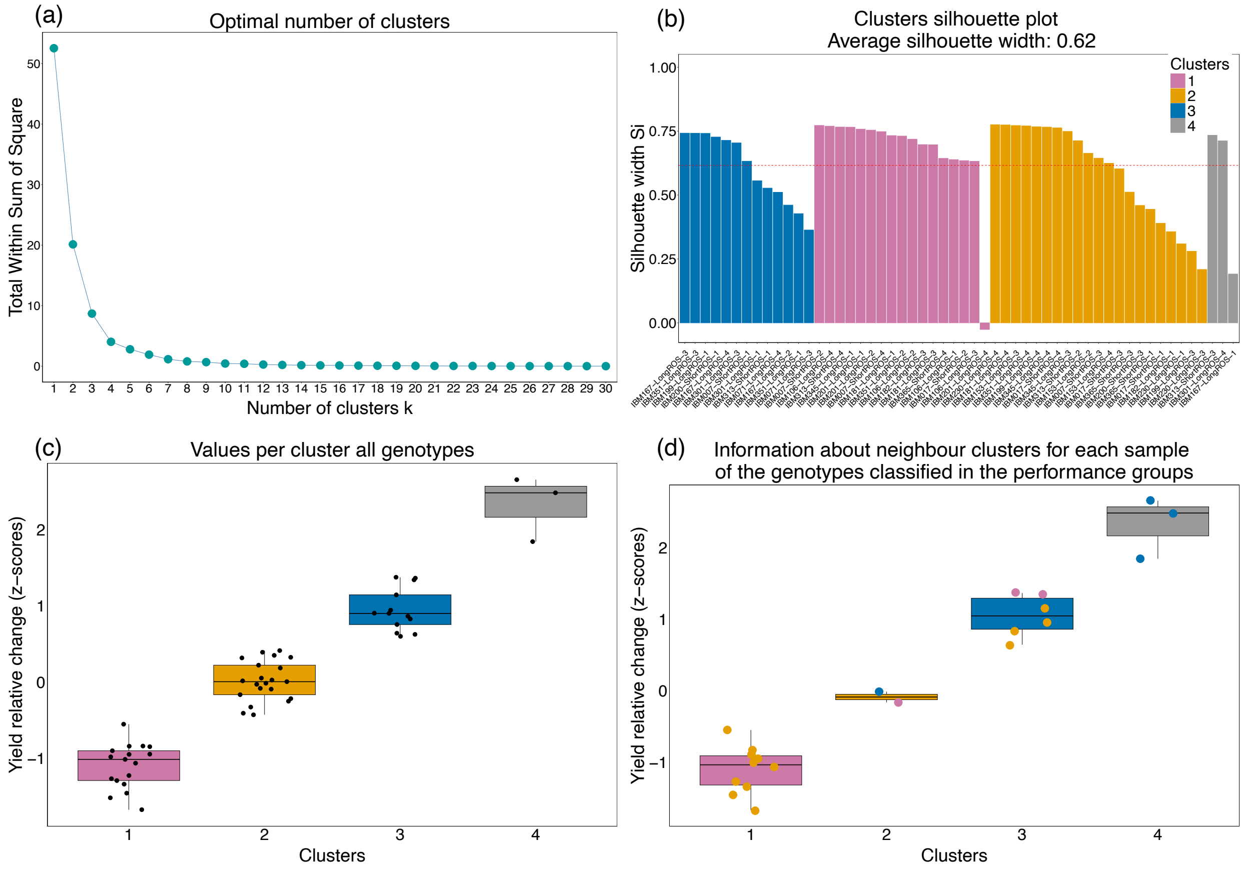


**Fig. S6 Characterization of k-medoids clusters based on relative yield change.** (a) Within-sum-of-squares plot used to select the best number of clusters (n = 4) using the elbow method; (b) robustness of the clustering indicated by the average silhouette width (0.62) and distribution of each sample into each cluster; (c) z-transformed relative yield change values for each cluster for all genotypes; (d) same z-transformed relative yield change values only for the genotypes selected in the two performance groups. In panel d the color of the dots represents the closest (“neighbour”) cluster identified by the silhouette analysis for each sample. This confirms the clear separation between cluster 1 and 4, while showing the closeness of cluster 4 to cluster 3, with the latter including also genotypes with higher variability in yield reduction that tend to associate to clusters 1 and 2 as second clustering option. The R script by Klein et al. (2020) was used for these analyses and plots, as also applied by Giuliano et al. (2026).

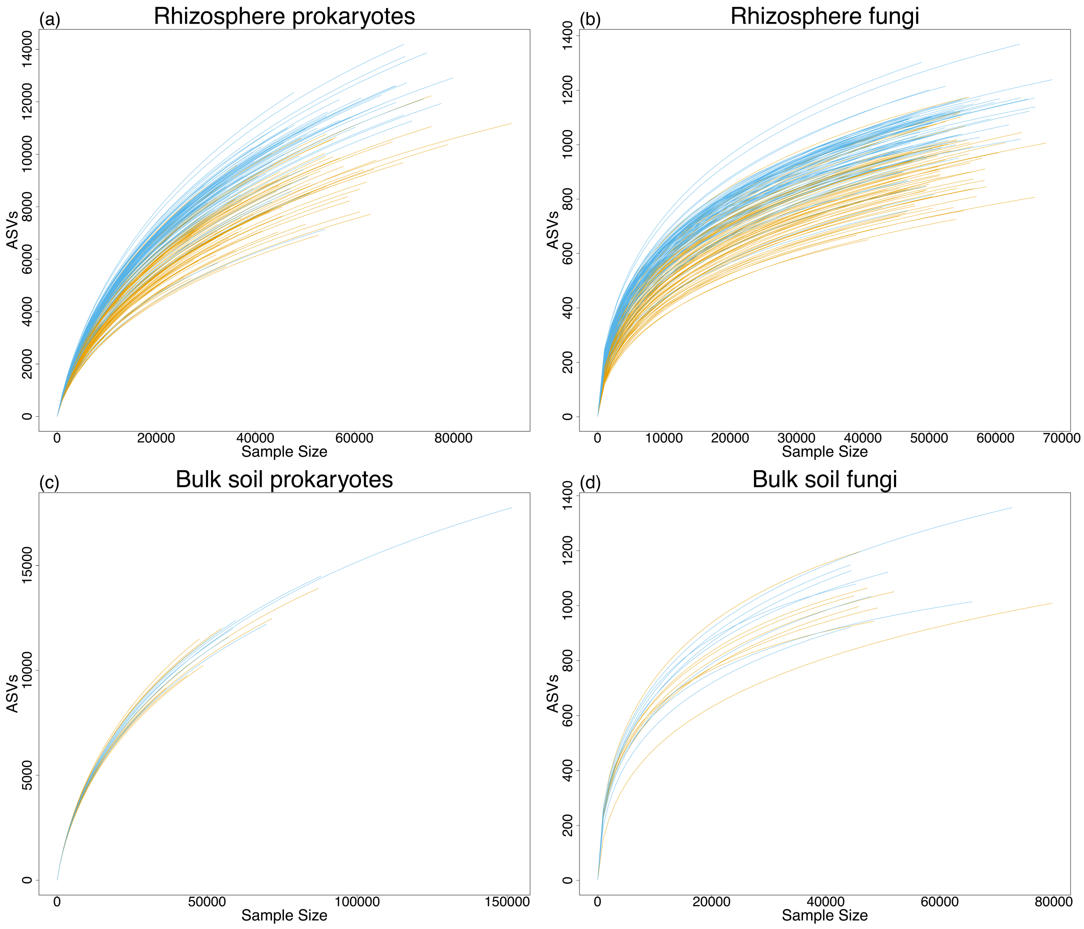


**Fig. S7 Rarefaction curves for rhizosphere and bulk soil prokaryotic and fungal communities.** The plots show the maximum number of ASVs found depending on the sample size (i.e., sequencing depth) for the rhizosphere and bulk soil prokaryotic (a,c) and fungal (b,d) samples under drought (orange) and control conditions (blue).

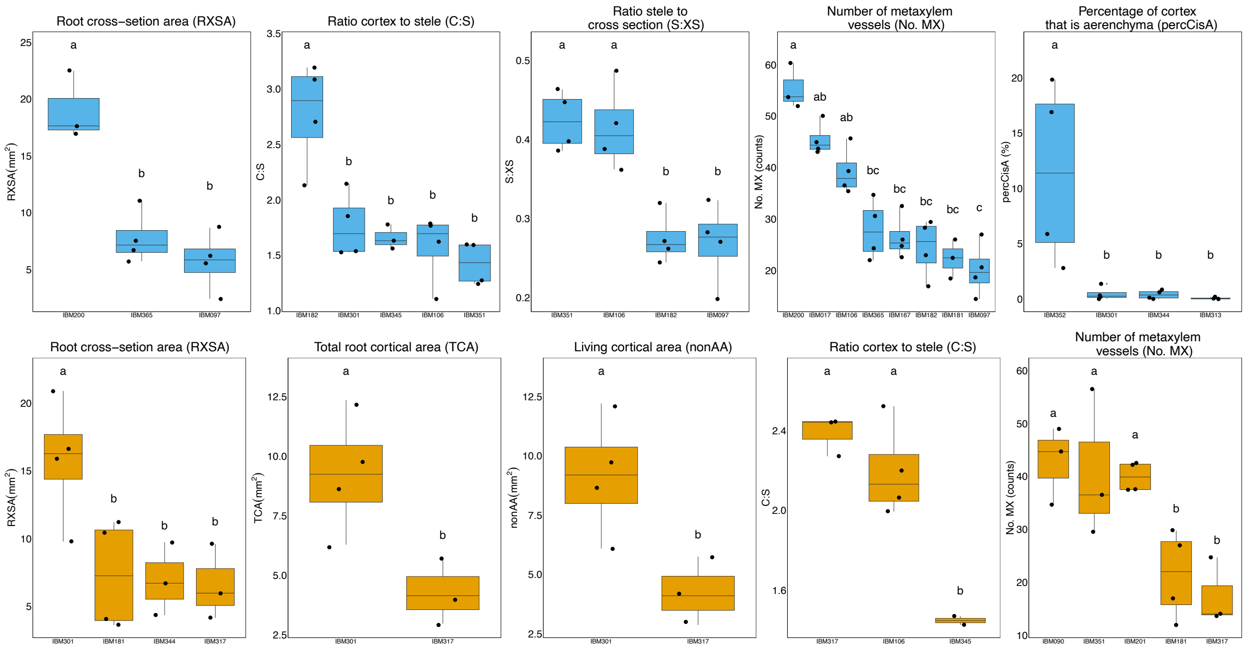


**Fig. S8 Root anatomical phenotypes differentiating across genotypes for each treatment.** Each boxplot reports single measures only for the anatomical phenotypes that resulted to be significantly different between genotypes (pairwise test based on the linear mixed-effects model, Treatment x Genotype, P-adjusted (Tukey) < 0.05) under control conditions (blue) and drought (orange). Total and living cortical area (TCA, nonAA) and cortex to cross-section ratio (C:XS) are not displayed because they had the same results in terms of difference between genotypes as root cross-section area (RXSA) and stele to cross-section ratio (S:XS) under control conditions, respectively.

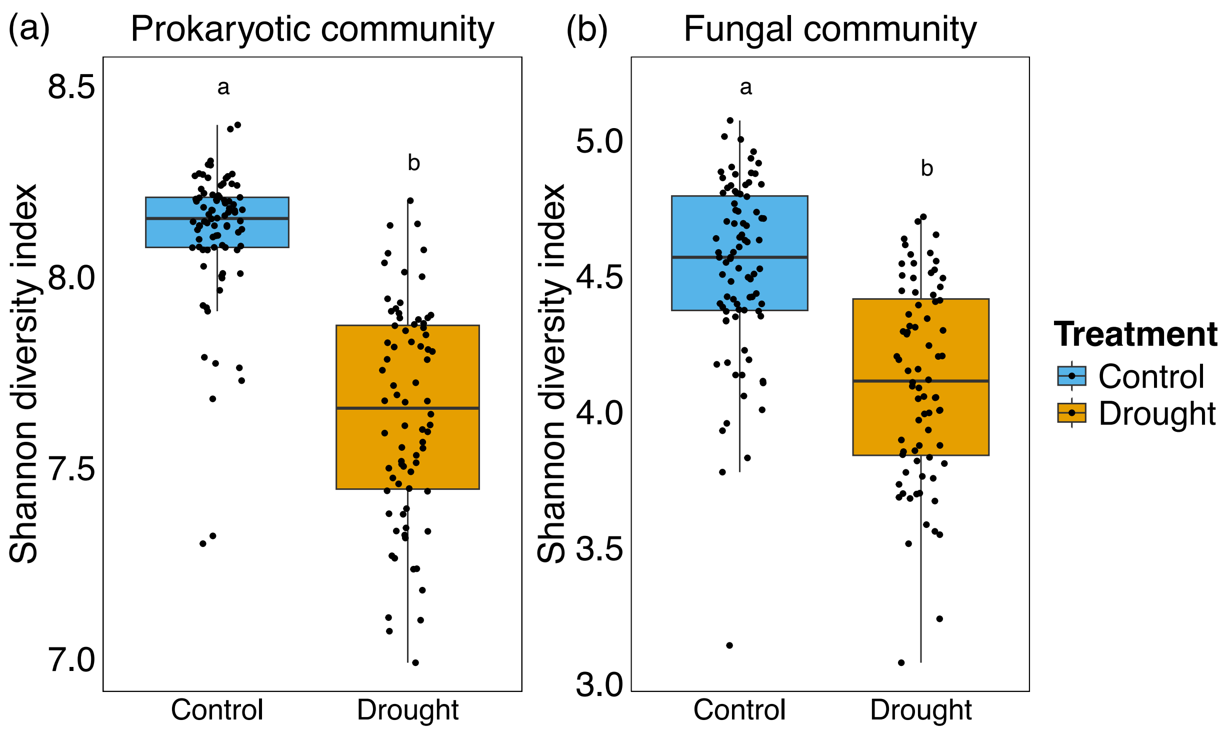


**Fig. S9 Alpha(α)-diversity (Shannon index) of prokaryotic and fungal communities in the rhizosphere.** Shannon index was significantly different (PERMANOVA, P = 0.0001) between treatments for prokaryotes (a) and fungi (b) as indicated by the letters. Heterogeneity of variances by treatment was statistically significant for prokaryotes (PERMDISP, P < 0.001).

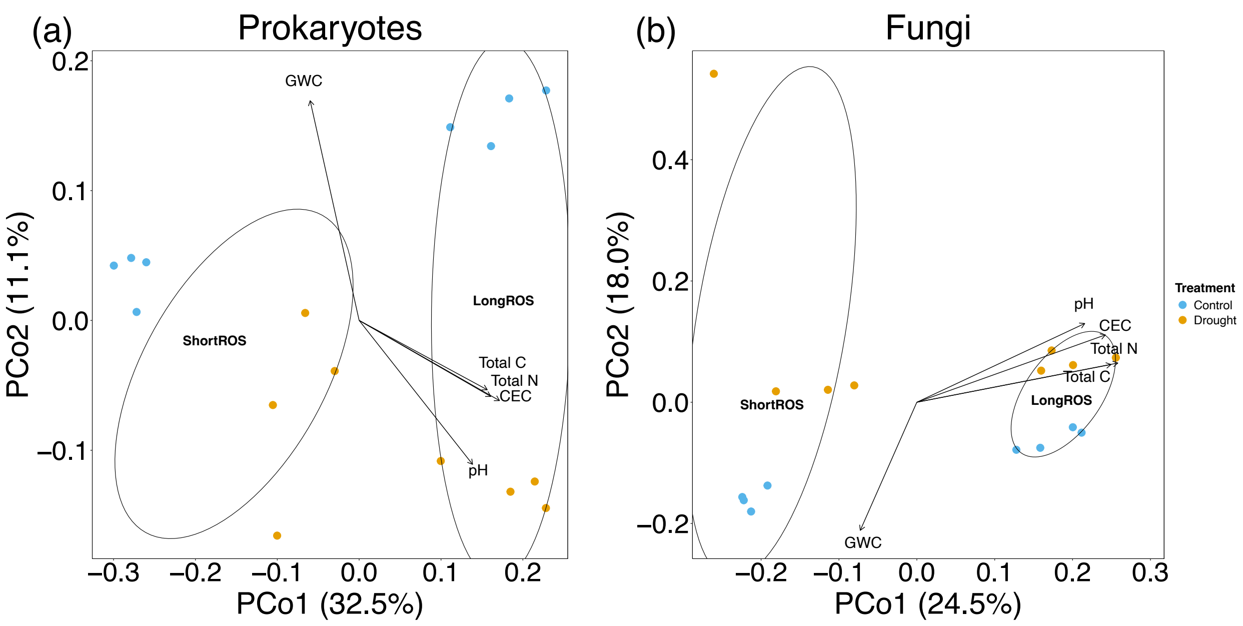


**Fig. S10 Separation of the bulk soil microbial communities by field and treatment.** Community structures (Bray-Curtis dissimilarity) of bulk soil prokaryotes (a) and fungi (b) are reported with Principal Coordinate analysis (PCoA) by treatment and field (Short or Long rain-out shelters ROS). The bulk soil physiochemical properties significantly ($P < 0.05$) affecting the observed separation between communities were identified with the *envfit* function (*vegan* package). They are the following: Gravimetric water content (GWC) and partially pH explaining the separation between treatments, and total nitrogen (Total N), carbon (Total C) and cation exchange capacity (CEC) explaining the dissimilarities between communities driven by the field. The magnitude and direction of the separation due to each property is represented by the respective arrows, which were proportionally scaled to fit the PCo1 and PCo2 value range.

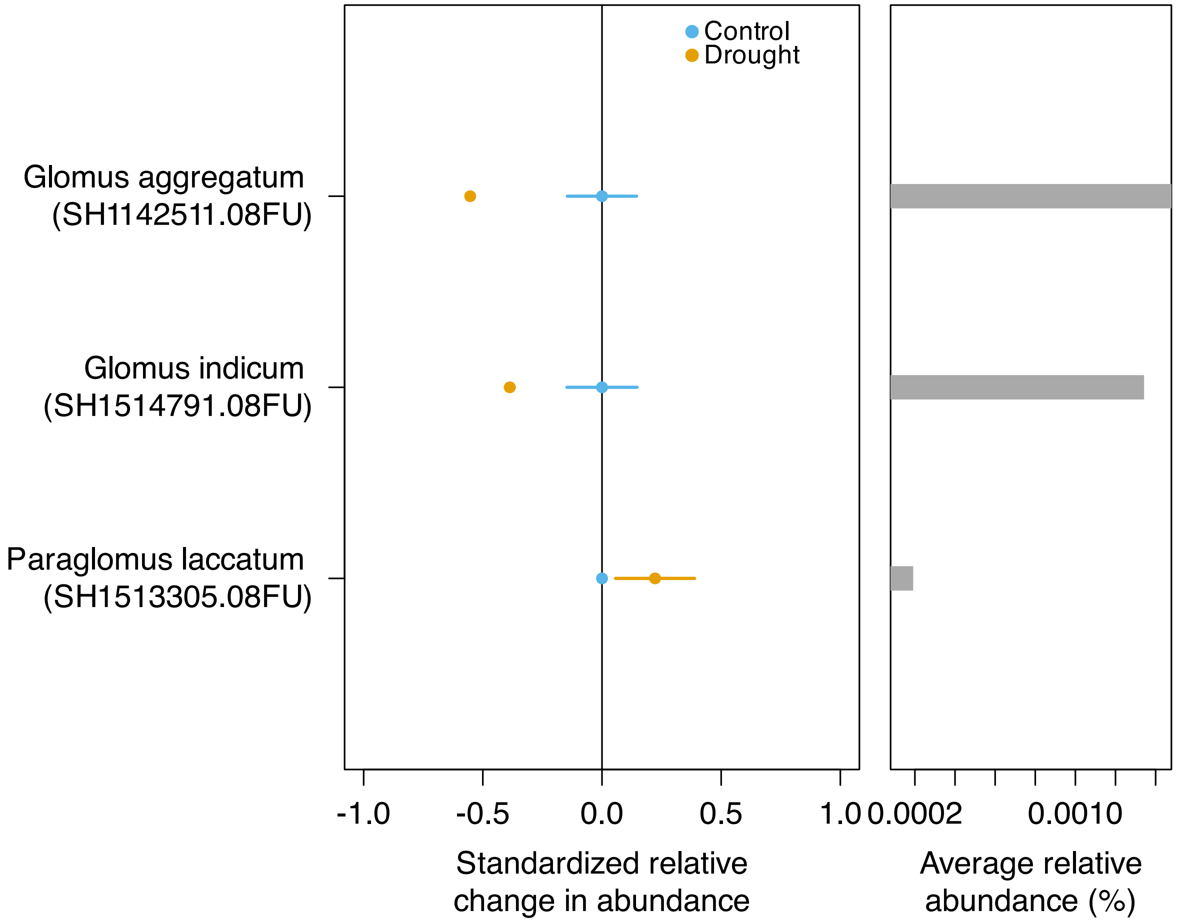


**Fig. S11 Influence of drought on arbuscular mycorrhizal fungi (AMF) species.** Relative change in the abundance of AMF species significantly influenced (PERMANOVA, $P < 0.05$) by drought are reported. Control is the zero line (Control mean – Control mean) and relative abundances (z-scores) under drought were calculated as (value under drought) – (value under control conditions) and reported with their standard error bars. The right side of the graph displays the relative abundance of each species calculated as average across all samples. Each species name is accompanied by the Species Hypothesis (SH) code and the information about the UNITE taxonomic classification dataset used (08FU). The screening of the AMF species was carried out by consulting the AMF database (AMF species list, former AMF-phylogeny) curated by Dr. Arthur Schussler, Dr. Chris Walker, Sidney Stürme, Franck Stefani and Jim Bever (CICG 2026).

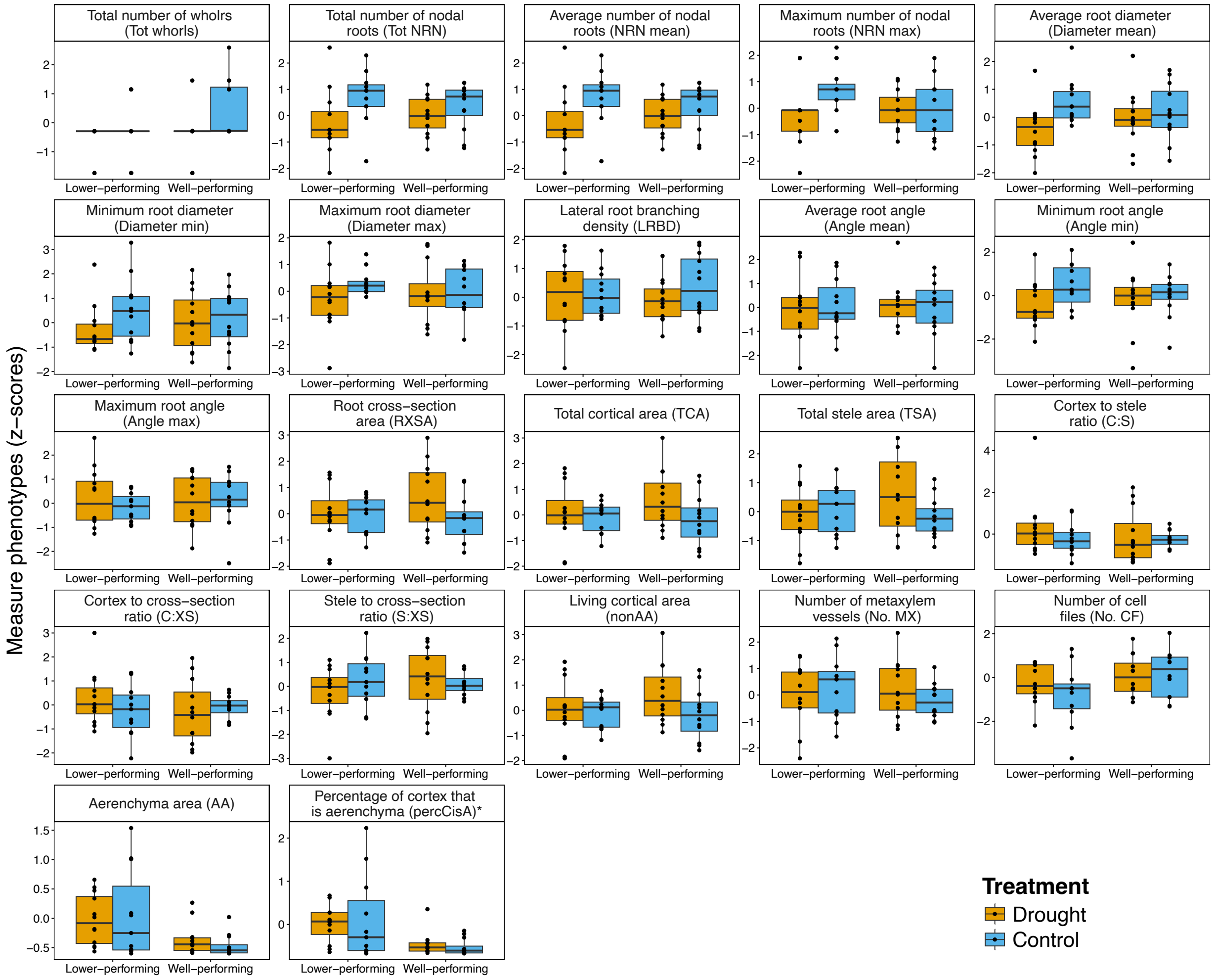


**Fig. S12 Root phenotypic variation across plant performance groups and treatments.** Root phenotypes (z-scores) in well- and lower-performing genotypes under drought and optimal water conditions. The asterisk (*) next to the phenotype name indicate that percCisA was significantly different between performance groups regardless of the treatment.

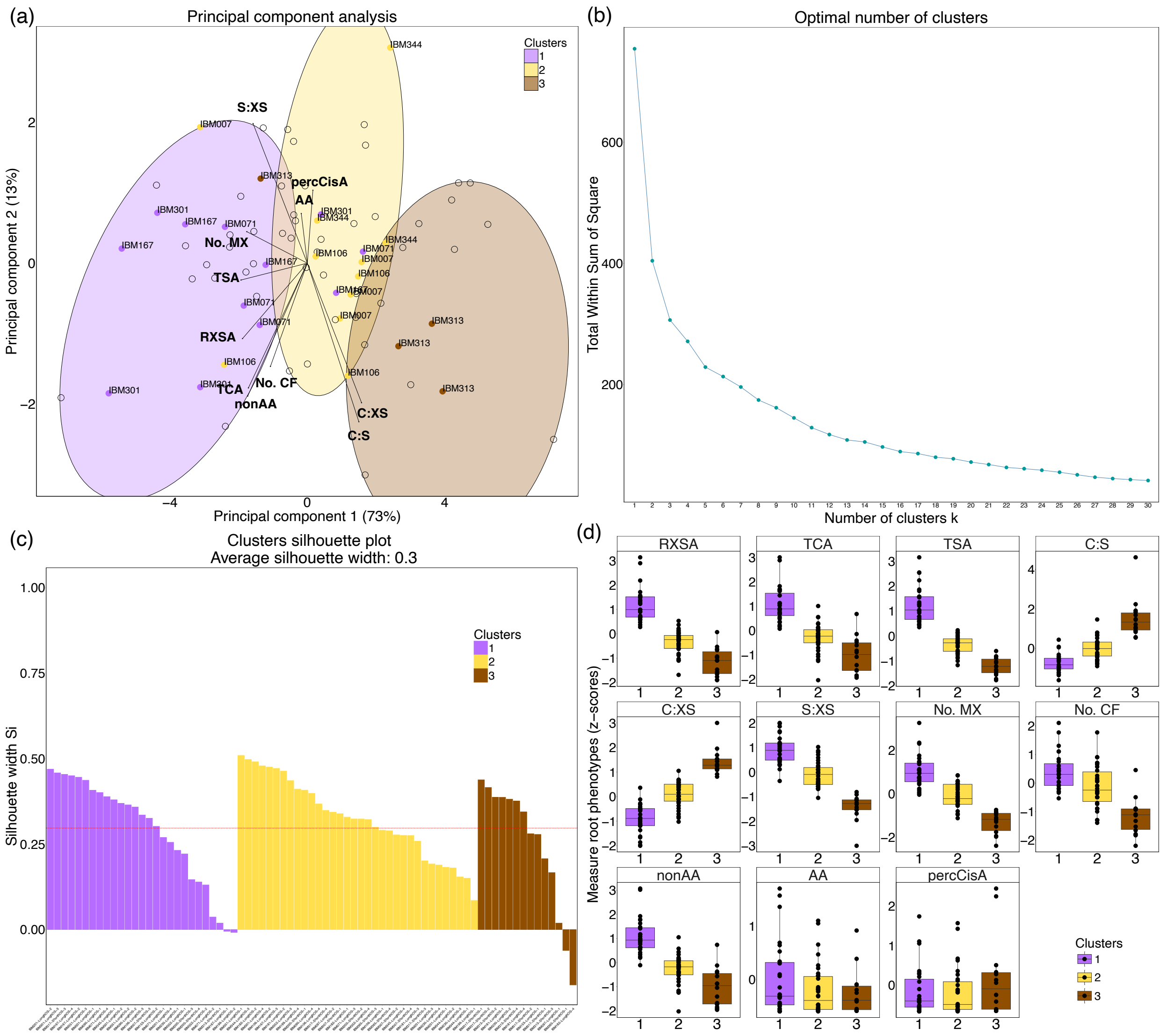


**Fig. S13 Anatomy-based k-medoids clusters under drought.** (a) Principal component analysis (PCA) based on anatomical phenotypes under drought overlapped with the three anatomy-based k-medoids clusters (colored ellipses). Genotypes with four available replicated values for anatomy which displayed more than half (at least 3) of their replicates belonging to one specific cluster were labeled and highlighted with the color of the respective cluster. The proportion of variance explained by each axis is displayed in brackets. (b) Total within-sum-of-squares plotted against the number of clusters used to select the cluster number (n=3) based on the elbow method. (c) Silhouette graph displaying the composition of each cluster and the clustering robustness (average silhouette width). (d) Anatomical phenotypes (z-scores) under drought distinguished by cluster.

The meaning of the root phenotype abbreviations is shown in Table 1. The R script by Klein et al. (2020) was used for these plots, as also applied by Giuliano et al. (2026).

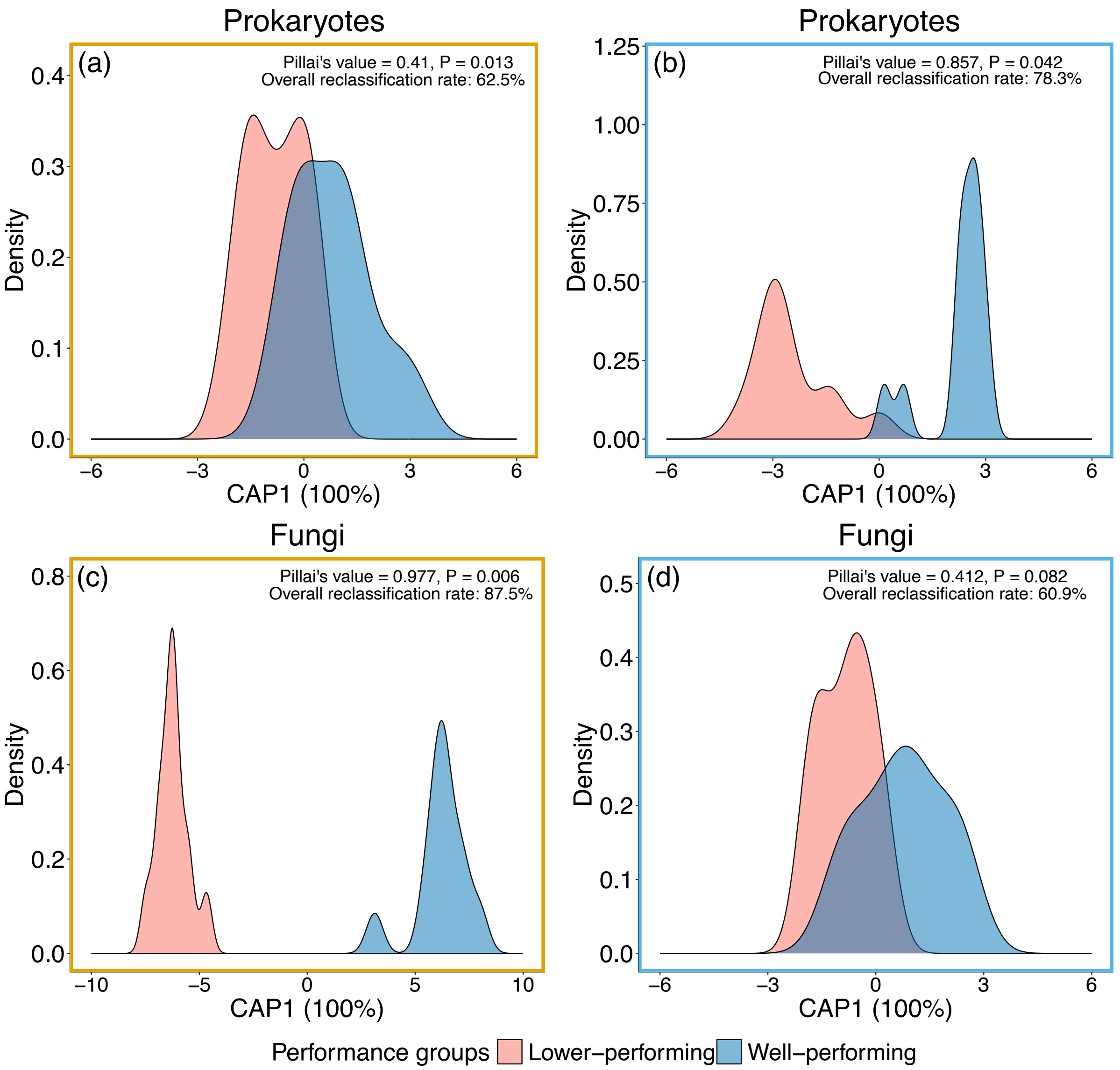


**Fig. S14 Separation between microbial communities belonging to the two plant performance groups.** The sample density in each performance group is plotted against the CAP score obtained through Canonical Analysis of Principal coordinates (CAP) on the Bray-Curtis dissimilarity built on normalized ASV counts for prokaryotes and fungi under drought (a,c) and control conditions (b,d). Only the six genotypes belonging to the two identified performance groups were included in the analysis. The CAP score represents the distance between samples along the only axis identified along which the communities separate the best (100%) based on the performance group. The Pillai's test results indicating the significant separation between groups and the reclassification rate are reported in each plot.

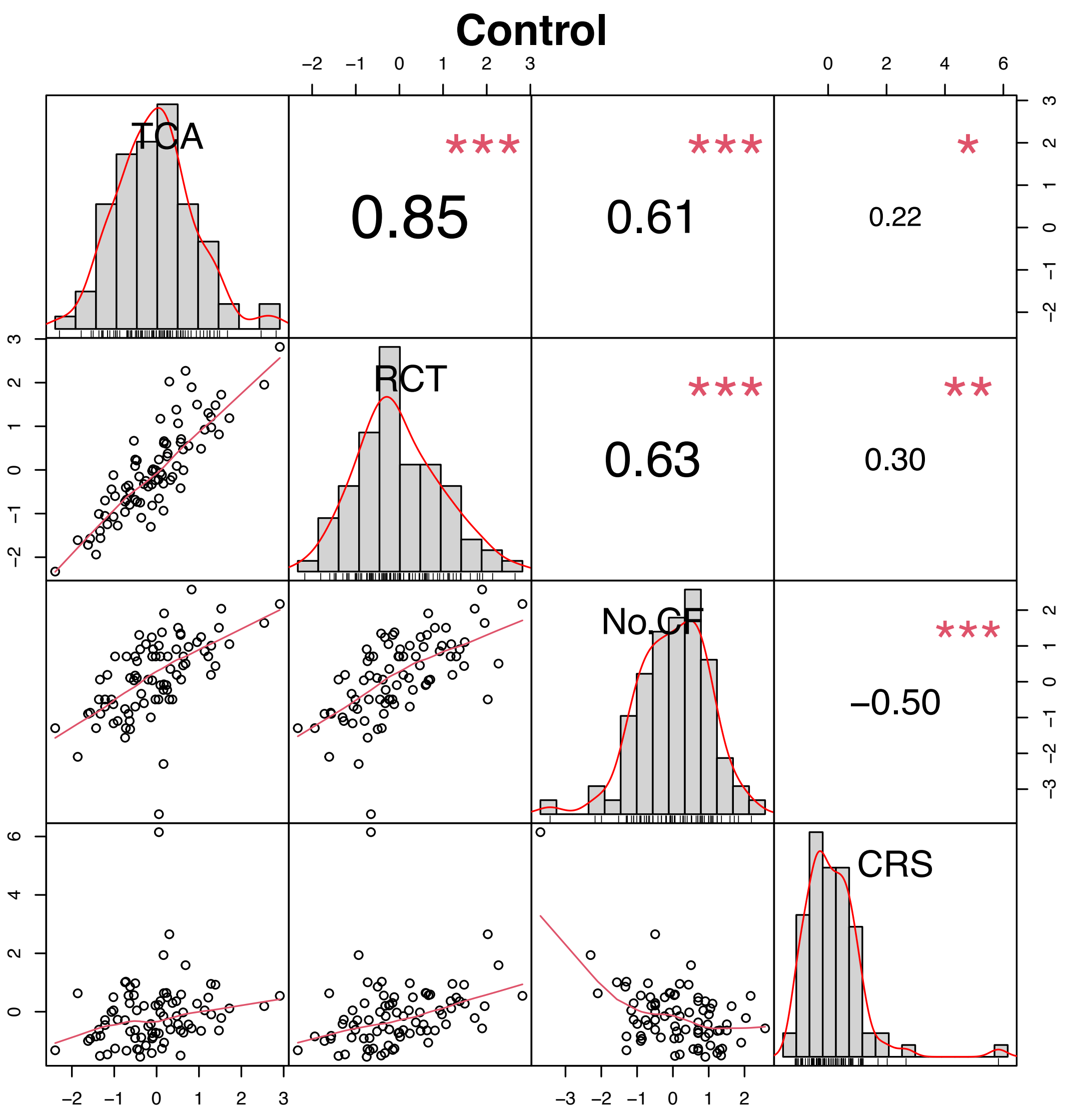


**Fig. S15 Pair-wise correlations among root cortical phenotypes under control conditions.** The scatter plots represent the relation between two of the four analyzed phenotypes (TCA, total cortical area, originally in $mm^2$; RCT, root cortical thickness, originally in mm; No. CF, number of cortical cell files; CRS, cortical cell radial size, originally in mm) for individual samples across the 22 genotypes. The plot axes report the z-scores as these phenotypes were normalized for comparability. The histograms represent the data distribution of each variable. The numbers in the squares are the Pearson's correlation coefficients for each couple of variables, and the red asterisks represent the level of statistically significance produced by the cor.test.

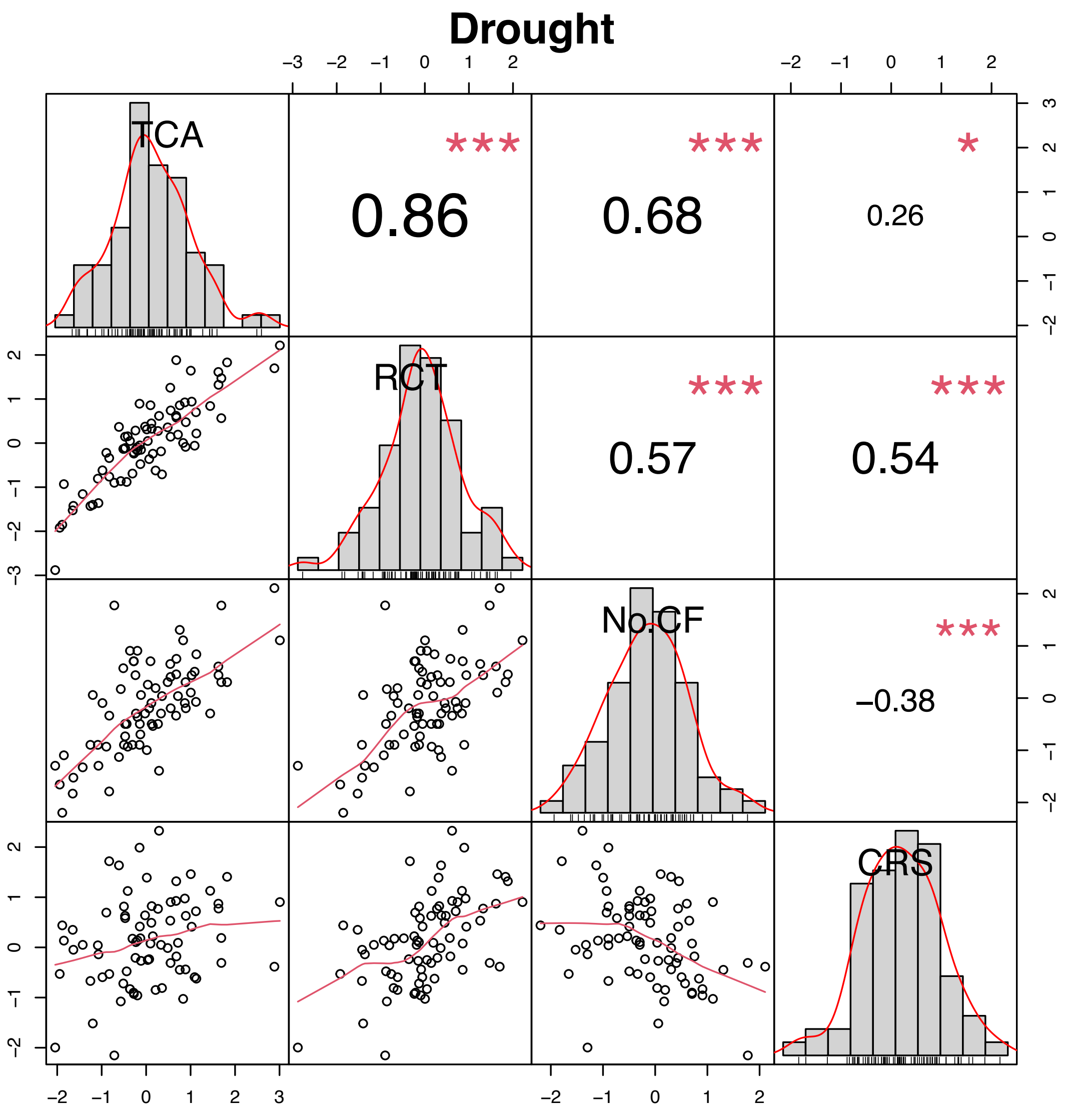


**Fig. S16 Pair-wise correlations among root cortical phenotypes under drought conditions.** The scatter plots represent the relation between two of the four analyzed phenotypes (TCA, total cortical area, originally in $mm^2$; RCT, root cortical thickness, originally in mm; No. CF, number of cortical cell files; CRS, cortical cell radial size, originally in mm) for individual samples across the 22 genotypes. The plot axes report the z-scores as these phenotypes were normalized for comparability. The histograms represent the data distribution of each variable. The numbers in the squares are the Pearson's correlation coefficients for each couple of variables, and the red asterisks represent the level of statistically significance produced by the cor.test.

**Table S1** Content of macro- and micronutrients in bulk soil for each field and treatment.

| | **Drought** | | | | **Control conditions** | | | |
|---|---|---|---|---|---|---|---|---|
| | **Short ROS** | | **Long ROS** | | **Short ROS** | | **Long ROS** | |
| | **Average (mg kg$^{-1}$)** | **SD** | **average (mg kg-$^{1}$)** | **SD** | **average (mg kg$^{-1}$)** | **SD** | **average (mg kg$^{-1}$)** | **SD** |
| Phosphorus (P) | 614.344 | ± 35.636 | 514.472 | ± 238.253 | 501.059 | ± 55.643 | 604.022 | ± 61.003 |
| Potassium (K) | 1466.981 | ± 104.583 | 1273.953 | ± 97.213 | 1285.980 | ± 230.120 | 1432.697 | ± 37.806 |
| Calcium (Ca) | 1551.476 | ± 107.471 | 2136.083 | ± 454.228 | 1180.272 | ± 78.120 | 1788.222 | ± 111.317 |
| Mangnesium (Mg) | 2056.275 | ± 134.981 | 1453.332 | ± 127.827 | 1819.194 | ± 260.677 | 1601.443 | ± 98.1962 |
| Sodium (Na) | 90.050 | ± 4.063 | 61.675 | ± 8.704 | 79.668 | ± 7.620 | 71.952 | ± 2.972 |
| Iron (Fe) | 23861.276 | ± 1035.055 | 20527.822 | ± 4179.822 | 21841.014 | ± 1183.762 | 24350.333 | ± 1181.552 |
| Manganese (Mn) | 1429.994 | ± 169.087 | 1093.095 | ± 184.952 | 1275.912 | ± 213.171 | 1325.021 | ± 98.841 |
| Zinc (Zn) | 94.651 | ± 8.114 | 47.655 | ± 3.874 | 50.009 | ± 3.604 | 50.631 | ± 2.163 |
| Copper (Cu) | 15.543 | ± 1.102 | 16.014 | ± 2.359 | 13.402 | ± 1.597 | 15.963 | ± 0.778 |
| Aluminium (Al) | 18214.647 | ± 742.390 | 15033.614 | ± 1469.054 | 16645.816 | ± 1910.812 | 17002.459 | ± 418.060 |
| Cobalt (Co) | 3.360 | ± 0.639 | 3.105 | ± 1.040 | 3.475 | ± 0.583 | 3.490 | ± 0.566 |
| Cadmium (Cd) | 0.498 | ± 0.003 | 0.621 | ± 0.252 | 0.496 | ± 0.002 | 0.499 | ± 0.002 |
| Lead (Pb) | 19.361 | ± 1.300 | 16.717 | ± 2.422 | 17.457 | ± 1.272 | 18.908 | ± 0.942 |

**Table S2** Number of available samples, missing values and main gap-filling and dataset combination strategies.

| Type of sample | Total number by design | Missing (totally) | Gap-filled | Total available number | Major changes to the final number |
|---|---|---|---|---|---|
| Grain yield | 2 treatments x 4 replicates x 22 genotypes = **176** | 36 | 0 | **140** | Due to limitations in data availability, samples missing for microbiome but present for yield were kept. Slightly adapted to match other datasets for statistics and correlations. |
| Root anatomy (all) | | 14 | 0 | **162** | To perform all main analyses, 4 and 13 samples for anatomy and architecture, respectively, were removed to combine the two datasets based only on the samples available for both and for microbiome analyses (n = **158** total) |
| Root architecture | | 5* | 3* | **171*** | |
| Rhizosphere soil | | 15 | 0 | **161** | Slightly adapted to match other datasets for statistics and correlations |
| Root anatomy (CPW) | 2 treatments x 4 replicates x 6 genotypes = **48** | 9 | 0 | **39** | Slightly adapted to match other datasets for statistics and correlations |

| | | | | | |
|---|---|---|---|---|---|
| Bulk soil | 2 fields x 2 treatments x 4 replicates = **16** | 0 | 0 | **16** | none |

[a]Note that one more sample from the total, corresponding to a failed plot for yield (n.41, IBM365, rep 2, drought) was removed from all datasets. *The indicated architectural missing samples did not have data for all measured variables and no data coming from the other plants sampled in the same plot were available. Regarding the architecture: *the gap-filled samples indicated are only the ones for which DNA samples and plant number information were available, but no architectural data for the same plant were present. Therefore, the average of the other two technical replicates was used; *the total available sample number includes not only the 3 gap-filled mentioned above but all gap-filled samples, regardless of if they were used in the analyses or not (details on gap-filling and samples removed by architectural and anatomical datasets are in Methods S1).

**Table S3** Main variables affecting root architecture and anatomy across 22 maize genotypes.

| **Architecture** | | | **Anatomy** | | |
|---|---|---|---|---|---|
| Treatment | | | Treatment x Genotype | | |
| | **F** | **P** | | **F** | **P** |
| LRBD | 2.29 | 0.1331 | RXSA* | 2.56 | **0.0009** |
| Tot whorls | 9.59 | **0.0025** | TCA* | 2.19 | **0.0050** |
| Tot NRN | 17.09 | **0.0001** | TSA* | 2.68 | **0.0005** |
| NRN mean | 17.06 | **0.0001** | C:S* | 4.07 | **<.0001** |
| NRN max** | 8.89 | **0.0035** | C:XS* | 2.22 | **0.0042** |
| Angle mean* | 1.59 | 0.2096 | S:XS* | 2.22 | **0.0042** |
| Angle min | 3.99 | **0.0484** | AA* | 1.48 | 0.0991 |
| Angle max | 0.02 | 0.8792 | percCisA* | 1.68 | **0.0458** |
| Diameter mean* | 8.69 | **0.0038** | nonAA* | 2.18 | **0.0052** |
| Diameter min | 0.01 | 0.9422 | No. MX* | 2.24 | **0.0037** |
| Diameter max | 17.06 | **0.0001** | No. CF | 1.31 | 0.1808 |

[a]Results of the linear mixed-effects model (LMM) testing the effect of treatment x genotype with field or block as random factor. Statistically significant results ($P < 0.05$) are highlighted. AA and percCisA were transformed with the *bestNormalize* function before running LLM. The single (*) and double (**) asterisks identify the phenotypes that were also overall significantly affected by the genotype or by treatment x genotype, respectively. Statistical model equations are provided in Supplementary Methods S3. The description of the acronyms is in the main text (Table 1).

**Table S4** Effect of genotype and root phenotypes on grain yield under drought according to permutational multivariate analysis of variance (PERMANOVA).

| **PERMANOVA** | | | |
|---|---|---|---|
| | **$R^2$** | **F** | **P** |
| **Genotype** | 0.593 | 5.173 | **0.0036** |
| Total number of nodal roots (Tot NRN) | 0.000 | 0.065 | 0.8078 |
| Lateral root branching density (LRBD) | 0.014 | 2.389 | 0.1403 |
| Total number of whorls (Tot whorls) | 0.000 | 0.047 | 0.8305 |
| Maximum number of nodal roots (NRN max) | 0.000 | 0.005 | 0.9518 |
| Average root angle (Angle mean) | 0.000 | 0.007 | 0.937 |
| Minimum root angle (Angle min) | 0.009 | 1.535 | 0.2344 |
| Maximum root angle (Angle max) | 0.004 | 0.669 | 0.446 |
| Average root diameter (Diameter mean) | 0.007 | 1.229 | 0.2727 |
| Minimum root diameter (Diameter min) | 0.002 | 0.262 | 0.6018 |
| Maximum root diameter (Diameter max) | 0.001 | 0.173 | 0.6874 |
| **Root cross-section area (RXSA)** | 0.038 | 6.301 | **0.0228** |
| Total cortical area (TCA) | 0.002 | 0.411 | 0.5035 |
| **Total stele area (TSA)** | 0.038 | 6.379 | **0.0116** |
| Cortex to stele ratio (C:S) | 0.003 | 0.477 | 0.4697 |
| **Cortex to cross-section ratio (C:XS)** | 0.041 | 6.745 | **0.0178** |
| Stele to cross-section ratio (S:XS) | 0.001 | 0.106 | 0.7429 |
| Cortical aerenchyma area (AA) | 0.009 | 1.479 | 0.2058 |
| Percentage of cortex that is aerenchyma (PercCisA) | 0.006 | 0.970 | 0.2441 |
| **Living cortical area (nonAA)** | 0.019 | 3.215 | **0.041** |
| **Number of metaxylem vessels (No. MX)** | 0.069 | 11.476 | **0.0063** |
| Number of cell files (No. CF) | 0.028 | 4.633 | 0.0575 |

**Table S5** Prokaryotic genera significantly affected (PERMANOVA, permutational multivariate analysis of variance, q < 0.05) by the maize genotype.

| Prokaryotes | | | | |
|---|---|---|---|---|
| | **$R^2$** | **F** | **P** | **q-value** |
| ADurb.Bin063-1 | 0.4781 | 12.4005 | 0.0004 | **0.0255** |
| Aeromicrobium | 0.2356 | 4.5830 | 0.0005 | **0.0266** |
| Aquisphaera | 0.3068 | 5.3010 | 0.0001 | **0.0106** |
| Candidatus Solibacter | 0.4574 | 15.1579 | 0.0001 | **0.0106** |
| Candidatus Xiphinematobacter | 0.2968 | 6.4588 | 0.0001 | **0.0106** |
| Flavitalea | 0.3346 | 7.0773 | 0.0003 | **0.0213** |
| Georgfuchsia | 0.2946 | 4.4821 | 0.0005 | **0.0266** |
| Haliangium | 0.1391 | 3.7243 | 0.0001 | **0.0106** |
| JGI 0001001-H03 | 0.3254 | 5.7091 | 0.0008 | **0.0364** |
| Mycobacterium | 0.2963 | 5.8737 | 0.0002 | **0.0159** |
| Ohtaekwangia | 0.3322 | 4.8024 | 0.0010 | **0.0425** |
| Pajaroellobacter | 0.1876 | 5.8067 | 0.0001 | **0.0106** |
| Panacagrimonas | 0.3855 | 4.5417 | 0.0001 | **0.0106** |
| Phenylobacterium | 0.2649 | 5.6214 | 0.0002 | **0.0159** |
| Thermobifida | 0.3659 | 5.0610 | 0.0006 | **0.0294** |

**Table S6** Effect of the treatment, genotype, their interaction and the root phenotypes on microbial β-diversity (permutational multivariate analysis of variance, PERMANOVA).

| | Total | | | | | | Control conditions | | | | | | Drought | | | | | |
|---|---|---|---|---|---|---|---|---|---|---|---|---|---|---|---|---|---|---|
| | Prokaryotes | | | Fungi | | | Prokaryotes | | | Fungi | | | Prokaryotes | | | Fungi | | |
| | $R^2$ | F | P | $R^2$ | F | P | $R^2$ | F | P | $R^2$ | F | P | $R^2$ | F | P | $R^2$ | F | P |
| *Treatment* | 0.126 | 27.242 | **0.0001** | 0.199 | 45.251 | **0.0001** | | | | | | | | | | | | |
| *Genotype* | 0.219 | 2.243 | **0.0096** | 0.187 | 2.018 | **0.0086** | 0.401 | 2.005 | 0.2935 | 0.421 | 2.214 | **0.004** | 0.389 | 1.557 | 0.7505 | 0.336 | 1.245 | 0.8203 |
| *Treatment x Genotype* | 0.119 | 1.226 | **0.0046** | 0.108 | 1.167 | 0.0673 | | | | | | | | | | | | |
| Total number of nodal roots (Tot NRN) | 0.005 | 1.031 | 0.3377 | 0.005 | 1.029 | 0.3476 | 0.009 | 0.966 | 0.4572 | 0.009 | 1.009 | 0.4299 | 0.011 | 0.954 | 0.4667 | 0.013 | 1.007 | 0.4062 |
| Lateral root branching density (LRBD) | 0.006 | 1.305 | 0.1235 | 0.005 | 1.114 | 0.2798 | 0.012 | 1.222 | 0.1337 | 0.010 | 1.139 | 0.2448 | 0.012 | 1.017 | 0.388 | 0.016 | 1.273 | 0.16 |
| Total number of whorls (Tot whorls) | 0.005 | 1.109 | 0.2451 | 0.005 | 1.111 | 0.2639 | 0.010 | 1.068 | 0.2634 | 0.012 | 1.356 | 0.0786 | 0.011 | 0.958 | 0.5098 | 0.011 | 0.830 | 0.699 |
| Maximum number of nodal roots (NRN max) | 0.004 | 0.908 | 0.509 | 0.003 | 0.666 | 0.8473 | 0.010 | 1.029 | 0.3318 | 0.007 | 0.723 | 0.9073 | 0.013 | 1.051 | 0.2691 | 0.011 | 0.889 | 0.5699 |
| Average root angle (Angle mean) | 0.007 | 1.595 | **0.0445** | 0.004 | 0.978 | 0.4002 | 0.014 | 1.434 | 0.0517 | 0.010 | 1.102 | 0.2885 | 0.011 | 0.958 | 0.4971 | 0.011 | 0.854 | 0.6583 |
| Minimum root angle (Angle min) | 0.004 | 0.819 | 0.749 | 0.003 | 0.780 | 0.7146 | 0.010 | 1.022 | 0.3541 | 0.010 | 1.049 | 0.3902 | 0.009 | 0.763 | 0.9406 | 0.010 | 0.776 | 0.7759 |
| Maximum root angle (Angle max) | 0.006 | 1.253 | 0.148 | 0.006 | 1.254 | 0.1799 | 0.010 | 1.043 | 0.3074 | 0.007 | 0.730 | 0.8977 | 0.008 | 0.665 | 0.9988 | 0.011 | 0.870 | 0.6287 |
| Average root diameter (Diameter mean) | 0.005 | 1.107 | 0.2557 | 0.005 | 1.068 | 0.3137 | 0.008 | 0.845 | 0.8068 | 0.009 | 1.006 | 0.4472 | 0.016 | 1.314 | 0.0777 | 0.015 | 1.180 | 0.2213 |
| Minimum root diameter (Diameter min) | 0.005 | 1.162 | 0.2163 | 0.005 | 1.025 | 0.3701 | 0.011 | 1.195 | 0.1414 | 0.014 | 1.572 | **0.0246** | 0.011 | 0.945 | 0.5839 | 0.012 | 0.947 | 0.5329 |
| Maximum root diameter (Diameter max) | 0.004 | 0.799 | 0.7581 | 0.004 | 0.964 | 0.4145 | 0.008 | 0.849 | 0.7747 | 0.010 | 1.090 | 0.2937 | 0.008 | 0.645 | 0.9983 | 0.010 | 0.745 | 0.808 |
| Root cross-section area (RXSA) | 0.008 | 1.687 | **0.0301** | 0.009 | 1.982 | **0.0341** | 0.012 | 1.309 | 0.0883 | 0.010 | 1.096 | 0.2928 | 0.009 | 0.761 | 0.9358 | 0.012 | 0.927 | 0.5415 |
| Total cortical area (TCA) | 0.004 | 0.958 | 0.45 | 0.004 | 0.896 | 0.5085 | 0.009 | 0.925 | 0.5613 | 0.011 | 1.241 | 0.146 | 0.011 | 0.901 | 0.6644 | 0.011 | 0.862 | 0.659 |
| Total stele area (TSA) | 0.007 | 1.520 | 0.0549 | 0.008 | 1.809 | 0.074 | 0.010 | 1.090 | 0.2551 | 0.013 | 1.446 | 0.0857 | 0.012 | 0.995 | 0.5736 | 0.010 | 0.759 | 0.8506 |
| Cortex to stele ratio (C:S) | 0.004 | 0.896 | 0.5302 | 0.005 | 1.186 | 0.2069 | 0.008 | 0.840 | 0.7624 | 0.011 | 1.174 | 0.2064 | 0.011 | 0.928 | 0.552 | 0.015 | 1.169 | 0.2395 |
| Cortex to cross-section ratio (C:XS) | 0.004 | 0.960 | 0.434 | 0.005 | 1.063 | 0.3113 | 0.011 | 1.168 | 0.1581 | 0.013 | 1.402 | 0.071 | 0.009 | 0.793 | 0.8989 | 0.016 | 1.223 | 0.1963 |
| Stele to cross-section ratio (S:XS) | 0.004 | 0.958 | 0.4458 | 0.004 | 0.945 | 0.4505 | 0.012 | 1.261 | 0.1115 | 0.012 | 1.363 | 0.0769 | 0.009 | 0.757 | 0.9555 | 0.011 | 0.865 | 0.642 |
| Cortical aerenchyma area (AA) | 0.004 | 0.879 | 0.6073 | 0.004 | 0.951 | 0.4526 | 0.009 | 0.987 | 0.4064 | 0.011 | 1.192 | 0.1923 | 0.012 | 1.005 | 0.456 | 0.013 | 0.996 | 0.4675 |
| Percentage of cortex that is aerenchyma (PercCisA) | 0.004 | 0.804 | 0.7351 | 0.004 | 0.926 | 0.4612 | 0.011 | 1.107 | 0.2268 | 0.010 | 1.092 | 0.2975 | 0.011 | 0.957 | 0.6782 | 0.009 | 0.710 | 0.9061 |
| Living cortical area (nonAA) | 0.005 | 1.171 | 0.2258 | 0.004 | 0.919 | 0.5624 | 0.009 | 0.900 | 0.6954 | 0.008 | 0.897 | 0.6463 | 0.010 | 0.819 | 0.9245 | 0.011 | 0.834 | 0.7557 |
| Number of metaxylem vessels (No. MX) | 0.005 | 1.058 | 0.3068 | 0.005 | 1.029 | 0.3512 | 0.021 | 2.158 | **0.0034** | 0.015 | 1.628 | **0.0174** | 0.013 | 1.107 | 0.217 | 0.016 | 1.221 | 0.1951 |
| Number of cell files (No. CF) | 0.007 | 1.404 | 0.084 | 0.005 | 1.144 | 0.2356 | 0.015 | 1.594 | **0.0262** | 0.014 | 1.527 | **0.0295** | 0.012 | 0.973 | 0.4909 | 0.009 | 0.737 | 0.8452 |

[a]PERMANOVA testing the effect of treatment x genotype and all root phenotypes using field as strata. Statistically significant results ($P < 0.05$) are in bold.

**Table S7** List of significant (Spearman, q < 0.1) correlations between root phenotypes and relative abundances of prokaryotic genera under drought and control conditions across all 22 genotypes.

**Prokaryotes**

| Treatment | Root phenotype | Genus | Phylum | Relative abundance under drought comapred to control | Corr (ρ) | P | q-value | Mean relative abundance (%) |
|---|---|---|---|---|---|---|---|---|
| Drought | Diameter mean | Aeribacillus | Firmicutes | not affected | -0.4314 | 0.0002 | 0.0590 | 0.0135 |
| Control | No. CF | Abditibacterium | Abditibacteriota | not affected | -0.1980 | 0.0745 | 0.0940 | 0.0129 |
| | No. CF | Achromobacter | Proteobacteria | increased | -0.2144 | 0.0531 | 0.0811 | 0.0252 |
| | No. CF | Acidicaldus | Proteobacteria | increased | -0.2741 | 0.0127 | 0.0356 | 0.0196 |
| | No. CF | Acidiphilium | Proteobacteria | increased | -0.3120 | 0.0043 | 0.0214 | 0.0086 |
| | No. CF | Acidipila-Silvibacterium | Acidobacteriota | decreased | -0.2025 | 0.0681 | 0.0901 | 0.0250 |
| | No. CF | Actinomycetospora | Actinobacteriota | increased | 0.2832 | 0.0099 | 0.0331 | 0.0233 |
| | No. CF | Actinoplanes | Actinobacteriota | not affected | -0.2737 | 0.0128 | 0.0356 | 0.1141 |
| | No. CF | Adhaeribacter | Bacteroidota | increased | 0.3397 | 0.0018 | 0.0163 | 0.0937 |
| | No. CF | Agromyces | Actinobacteriota | decreased | 0.2651 | 0.0161 | 0.0373 | 0.0532 |
| | No. CF | AKYG587 | Planctomycetota | decreased | 0.2539 | 0.0214 | 0.0430 | 0.0211 |
| | No. CF | Allorhizobium-Neorhizobium-Pararhizobium-Rhizobium | Proteobacteria | increased | -0.2812 | 0.0105 | 0.0331 | 0.7675 |
| | No. CF | Alsobacter | Proteobacteria | not affected | -0.3100 | 0.0046 | 0.0220 | 0.0167 |
| | No. CF | Altererythrobacter | Proteobacteria | increased | -0.2199 | 0.0471 | 0.0752 | 0.0930 |
| | No. CF | Amycolatopsis | Actinobacteriota | increased | -0.2850 | 0.0094 | 0.0331 | 0.2251 |
| | No. CF | Angustibacter | Actinobacteriota | increased | -0.3885 | 0.0003 | 0.0073 | 0.1046 |
| | No. CF | Bauldia | Proteobacteria | decreased | 0.2326 | 0.0355 | 0.0624 | 0.0490 |
| | No. CF | Bradyrhizobium | Proteobacteria | decreased | -0.4136 | 0.0001 | 0.0052 | 0.5587 |
| | No. CF | Bryobacter | Acidobacteriota | increased | -0.2820 | 0.0103 | 0.0331 | 0.8464 |
| | No. CF | Burkholderia-Caballeronia-Paraburkholderia | Proteobacteria | increased | -0.4447 | 0.00003 | 0.0020 | 0.5417 |
| | No. CF | Caenimonas | Proteobacteria | increased | 0.2033 | 0.0670 | 0.0895 | 0.0314 |
| | No. CF | Candidatus Alysiosphaera | Proteobacteria | decreased | 0.3406 | 0.0017 | 0.0163 | 0.1069 |
| | No. CF | Candidatus Nitrososphaera | Crenarchaeota | not affected | 0.2566 | 0.0199 | 0.0414 | 0.0108 |
| | No. CF | Candidatus Protochlamydia | Verrucomicrobiota | decreased | 0.2549 | 0.0208 | 0.0426 | 0.0097 |
| | No. CF | Candidatus Udaeobacter | Verrucomicrobiota | decreased | 0.2701 | 0.0141 | 0.0370 | 0.6298 |
| | No. CF | Candidatus Xiphinematobacter | Verrucomicrobiota | decreased | 0.2389 | 0.0307 | 0.0553 | 0.1304 |
| | No. CF | Catenulispora | Actinobacteriota | not affected | -0.2462 | 0.0258 | 0.0490 | 0.0873 |
| | No. CF | Chitinophaga | Bacteroidota | increased | -0.2282 | 0.0392 | 0.0664 | 0.3404 |
| | No. CF | CL500-29 marine group | Actinobacteriota | decreased | 0.2051 | 0.0645 | 0.0888 | 0.0857 |
| | No. CF | Conexibacter | Actinobacteriota | increased | -0.2342 | 0.0342 | 0.0609 | 0.2483 |
| | No. CF | Deinococcus | Deinococcota | not affected | -0.2104 | 0.0577 | 0.0853 | 0.0279 |
| | No. CF | Devosia | Proteobacteria | not affected | -0.3170 | 0.0037 | 0.0199 | 0.3131 |
| | No. CF | Dokdonella | Proteobacteria | decreased | -0.4884 | 0.000003 | 0.0005 | 0.0744 |
| | No. CF | Dyella | Proteobacteria | decreased | -0.2822 | 0.0102 | 0.0331 | 0.0277 |
| | No. CF | Ellin516 | Verrucomicrobiota | decreased | -0.2947 | 0.0072 | 0.0270 | 0.0207 |
| | No. CF | Ferruginibacter | Bacteroidota | decreased | 0.2741 | 0.0127 | 0.0356 | 0.1592 |
| | No. CF | Filomicrobium | Proteobacteria | not affected | 0.2423 | 0.0283 | 0.0517 | 0.0202 |
| | No. CF | Flavitalea | Bacteroidota | increased | 0.2473 | 0.0251 | 0.0484 | 0.1210 |
| | No. CF | Geminicoccus | Proteobacteria | increased | 0.2069 | 0.0622 | 0.0872 | 0.0191 |
| | No. CF | Gemmatimonas | Gemmatimonadota | decreased | -0.4032 | 0.0002 | 0.0060 | 1.2330 |
| | No. CF | Geodermatophilus | Actinobacteriota | increased | -0.3361 | 0.0020 | 0.0163 | 0.2207 |
| | No. CF | Granulicella | Acidobacteriota | decreased | -0.2998 | 0.0062 | 0.0239 | 0.0180 |
| | No. CF | Iamia | Actinobacteriota | not affected | 0.2717 | 0.0135 | 0.0362 | 0.1061 |
| | No. CF | Ideonella | Proteobacteria | decreased | 0.2589 | 0.0188 | 0.0407 | 0.0183 |
| | No. CF | Ilumatobacter | Actinobacteriota | decreased | 0.2874 | 0.0089 | 0.0324 | 0.0641 |
| | No. CF | IMCC26207 | Actinobacteriota | not affected | 0.2660 | 0.0157 | 0.0373 | 0.0071 |
| | No. CF | Intrasporangium | Actinobacteriota | not affected | -0.2213 | 0.0457 | 0.0738 | 0.0092 |
| | No. CF | Jatrophihabitans | Actinobacteriota | not affected | -0.3348 | 0.0021 | 0.0163 | 0.1307 |
| | No. CF | JGI 0001001-H03 | Acidobacteriota | not affected | 0.2652 | 0.0160 | 0.0373 | 0.0513 |
| | No. CF | Kitasatospora | Actinobacteriota | decreased | -0.3300 | 0.0025 | 0.0175 | 0.0437 |
| | No. CF | Kribbella | Actinobacteriota | increased | -0.2001 | 0.0714 | 0.0916 | 0.4124 |
| | No. CF | Legionella | Proteobacteria | decreased | -0.2043 | 0.0656 | 0.0888 | 0.0640 |
| | No. CF | Leifsonia | Actinobacteriota | not affected | -0.2480 | 0.0247 | 0.0483 | 0.0082 |
| | No. CF | Litorilinea | Chloroflexi | decreased | 0.3401 | 0.0018 | 0.0163 | 0.0387 |
| | No. CF | Longispora | Actinobacteriota | not affected | 0.2077 | 0.0612 | 0.0867 | 0.0106 |
| | No. CF | Luteimonas | Proteobacteria | increased | -0.2091 | 0.0594 | 0.0859 | 0.0480 |
| | No. CF | Luteitalea | Acidobacteriota | not affected | 0.2680 | 0.0149 | 0.0373 | 0.0346 |
| | No. CF | Luteolibacter | Verrucomicrobiota | not affected | 0.2120 | 0.0559 | 0.0834 | 0.1819 |
| | No. CF | Lysinimonas | Actinobacteriota | decreased | -0.3881 | 0.0003 | 0.0073 | 0.0167 |
| | No. CF | Massilia | Proteobacteria | increased | -0.3416 | 0.0017 | 0.0163 | 1.5206 |
| | No. CF | Mesorhizobium | Proteobacteria | increased | -0.3192 | 0.0035 | 0.0199 | 0.6559 |
| | No. CF | Methylobacterium-Methylorubrum | Proteobacteria | not affected | -0.2270 | 0.0402 | 0.0665 | 0.0843 |
| | No. CF | Microvirga | Proteobacteria | not affected | 0.2773 | 0.0117 | 0.0345 | 0.2388 |
| | No. CF | Modestobacter | Actinobacteriota | increased | -0.3169 | 0.0037 | 0.0199 | 0.0115 |
| | No. CF | Mucilaginibacter | Bacteroidota | decreased | -0.3008 | 0.0060 | 0.0239 | 0.1646 |
| | No. CF | Nannocystis | Myxococcota | decreased | 0.2190 | 0.0480 | 0.0754 | 0.0229 |
| | No. CF | Neochlamydia | Verrucomicrobiota | decreased | 0.3123 | 0.0043 | 0.0214 | 0.0090 |
| | No. CF | Nitrolancea | Chloroflexi | increased | -0.2444 | 0.0269 | 0.0498 | 0.2224 |
| | No. CF | Nocardia | Actinobacteriota | increased | -0.2017 | 0.0692 | 0.0906 | 0.0912 |
| | No. CF | Nocardioides | Actinobacteriota | increased | -0.2658 | 0.0158 | 0.0373 | 1.2082 |
| | No. CF | Nordella | Proteobacteria | decreased | 0.3011 | 0.0060 | 0.0239 | 0.1393 |
| | No. CF | Novosphingobium | Proteobacteria | decreased | -0.2312 | 0.0366 | 0.0635 | 0.0542 |

| | | | | | | | | |
|---|---|---|---|---|---|---|---|---|
| | No. CF | Ohtaekwangia | Bacteroidota | increased | 0.2304 | 0.0373 | 0.0640 | 0.0606 |
| | No. CF | Ornithinibacillus | Firmicutes | not affected | -0.2217 | 0.0454 | 0.0738 | 0.0266 |
| | No. CF | Panacagrimonas | Proteobacteria | not affected | -0.2684 | 0.0148 | 0.0373 | 0.0062 |
| | No. CF | Paraclostridium | Firmicutes | decreased | 0.2086 | 0.0600 | 0.0859 | 0.0231 |
| | No. CF | Paucisalibacillus | Firmicutes | increased | -0.1999 | 0.0718 | 0.0916 | 0.0108 |
| | No. CF | Pedococcus-Phycicoccus | Actinobacteriota | increased | -0.3044 | 0.0054 | 0.0235 | 0.0681 |
| | No. CF | Pedomicrobium | Proteobacteria | decreased | 0.2585 | 0.0191 | 0.0407 | 0.2147 |
| | No. CF | Phaselicystis | Myxococcota | decreased | 0.2041 | 0.0659 | 0.0888 | 0.0679 |
| | No. CF | Phenylobacterium | Proteobacteria | decreased | -0.3549 | 0.0011 | 0.0163 | 0.1095 |
| | No. CF | Pir4 lineage | Planctomycetota | decreased | 0.3004 | 0.0061 | 0.0239 | 0.1831 |
| | No. CF | Pirellula | Planctomycetota | decreased | 0.3055 | 0.0052 | 0.0235 | 0.3955 |
| | No. CF | possible genus 04 | Fibrobacterota | decreased | -0.3623 | 0.0008 | 0.0163 | 0.0454 |
| | No. CF | Pseudolabrys | Proteobacteria | not affected | -0.3213 | 0.0032 | 0.0199 | 0.2504 |
| | No. CF | RB41 | Acidobacteriota | decreased | 0.3293 | 0.0025 | 0.0175 | 0.6379 |
| | No. CF | Rhizobacter | Proteobacteria | decreased | 0.2511 | 0.0229 | 0.0454 | 0.0822 |
| | No. CF | Rhodanobacter | Proteobacteria | decreased | -0.3374 | 0.0019 | 0.0163 | 0.2077 |
| | No. CF | Rhodomicrobium | Proteobacteria | decreased | 0.2449 | 0.0266 | 0.0498 | 0.0053 |
| | No. CF | Rhodoplanes | Proteobacteria | decreased | 0.2639 | 0.0166 | 0.0378 | 0.1953 |
| | No. CF | Roseimicrobium | Verrucomicrobiota | not affected | 0.2565 | 0.0200 | 0.0414 | 0.1398 |
| | No. CF | Roseisolibacter | Gemmatimonadota | not affected | -0.2604 | 0.0181 | 0.0406 | 0.0137 |
| Control | No. CF | Rubellimicrobium | Proteobacteria | increased | -0.1965 | 0.0768 | 0.0961 | 0.0992 |
| | No. CF | Ruminiclostridium | Firmicutes | not affected | 0.2717 | 0.0135 | 0.0362 | 0.0094 |
| | No. CF | Segetibacter | Bacteroidota | increased | -0.2798 | 0.0109 | 0.0334 | 0.0776 |
| | No. CF | Singulisphaera | Planctomycetota | decreased | -0.3076 | 0.0049 | 0.0229 | 0.2138 |
| | No. CF | Solirubrobacter | Actinobacteriota | not affected | 0.2090 | 0.0595 | 0.0859 | 0.4549 |
| | No. CF | Sorangium | Myxococcota | decreased | 0.3506 | 0.0012 | 0.0163 | 0.0202 |
| | No. CF | Sphaerimonospora | Actinobacteriota | decreased | 0.1998 | 0.0719 | 0.0916 | 0.0112 |
| | No. CF | Sphaerisporangium | Actinobacteriota | decreased | 0.2792 | 0.0111 | 0.0334 | 0.0152 |
| | No. CF | Sphingobium | Proteobacteria | increased | -0.2585 | 0.0190 | 0.0407 | 1.1848 |
| | No. CF | Sphingomonas | Proteobacteria | increased | -0.2838 | 0.0098 | 0.0331 | 1.9436 |
| | No. CF | Stenotrophobacter | Acidobacteriota | increased | 0.1944 | 0.0801 | 0.0993 | 0.0201 |
| | No. CF | Subgroup 10 | Acidobacteriota | decreased | 0.3192 | 0.0035 | 0.0199 | 0.1594 |
| | No. CF | Tardiphaga | Proteobacteria | not affected | -0.2273 | 0.0400 | 0.0665 | 0.0112 |
| | No. CF | Tepidisphaera | Planctomycetota | decreased | -0.3385 | 0.0019 | 0.0163 | 0.0628 |
| | No. CF | Terrabacter | Actinobacteriota | increased | -0.2133 | 0.0544 | 0.0821 | 0.2741 |
| | No. CF | Terrimonas | Bacteroidota | decreased | 0.3350 | 0.0021 | 0.0163 | 0.0868 |
| | No. CF | Thermobifida | Actinobacteriota | increased | 0.2186 | 0.0485 | 0.0754 | 0.0088 |
| | No. CF | TM7a | Patescibacteria | not affected | -0.2183 | 0.0488 | 0.0754 | 0.0176 |
| | No. CF | Tundrisphaera | Planctomycetota | decreased | -0.2673 | 0.0152 | 0.0373 | 0.1125 |
| | No. CF | Vicinamibacter | Acidobacteriota | not affected | 0.3226 | 0.0031 | 0.0199 | 0.0110 |
| | No. CF | YC-ZSS-LKJ147 | Gemmatimonadota | not affected | 0.2045 | 0.0653 | 0.0888 | 0.0258 |

[a]Mean relative abundance (%) was calculated across all samples for the two treatments combined. Explanation of the acronyms of the root phenotypes is reported in Table 1 in the main text.

**Table S8** List of significant (Spearman, q < 0.1) correlations between root phenotypes and relative abundances of fungal genera under drought and control conditions across all 22 genotypes.

| Fungi | | | | | | | | |
|---|---|---|---|---|---|---|---|---|
| **Treatment** | **Root phenotype** | **Genus** | **Phylum** | **Relative abundance under drought comapred to control** | **Corr ($\rho$)** | **P** | **q-value** | **Mean relative abundance (%)** |
| Drought | No. CF | Gliomastix | Ascomycota | increased | 0.3576 | 0.0016 | 0.0867 | 0.0543 |
| | No. CF | Triparticalcar | Chytridiomycota | not affected | 0.3511 | 0.0020 | 0.0867 | 0.0252 |
| Control | RXSA | Acremonium | Ascomycota | increased | 0.3637 | 0.0008 | 0.0414 | 2.9136 |
| | RXSA | Conocybe | Basidiomycota | not affected | -0.3778 | 0.0005 | 0.0414 | 0.1942 |
| | TCA | Acremonium | Ascomycota | increased | 0.3600 | 0.0009 | 0.0359 | 2.9136 |
| | TCA | Conocybe | Basidiomycota | not affected | -0.3833 | 0.0004 | 0.0304 | 0.1942 |
| | percCisA | Metacordyceps | Ascomycota | increased | -0.3324 | 0.0023 | 0.0750 | 0.2427 |
| | percCisA | Saitozyma | Basidiomycota | decreased | -0.3637 | 0.0008 | 0.0514 | 0.0090 |
| | nonAA | Acremonium | Ascomycota | increased | 0.3422 | 0.0017 | 0.0574 | 2.9136 |
| | nonAA | Conocybe | Basidiomycota | not affected | -0.3915 | 0.0003 | 0.0192 | 0.1942 |
| | No. MX | Acremonium | Ascomycota | increased | 0.3717 | 0.0006 | 0.0422 | 2.9136 |
| | No. MX | Entoloma | Basidiomycota | increased | 0.3337 | 0.0022 | 0.0789 | 0.2277 |
| | No. CF | Acremonium | Ascomycota | increased | 0.3429 | 0.0016 | 0.0259 | 2.9136 |
| | No. CF | Conlarium | Ascomycota | decreased | -0.2484 | 0.0244 | 0.0799 | 0.2665 |
| | No. CF | Conocybe | Basidiomycota | not affected | -0.3593 | 0.0009 | 0.0259 | 0.1942 |
| | No. CF | Devriesia | Ascomycota | increased | -0.2872 | 0.0089 | 0.0471 | 0.0248 |
| | No. CF | Exophiala | Ascomycota | not affected | -0.3472 | 0.0014 | 0.0259 | 3.0619 |
| | No. CF | Gongronella | Mucoromycota | increased | -0.2573 | 0.0196 | 0.0787 | 0.2033 |
| | No. CF | Hyaloscypha | Ascomycota | not affected | -0.2362 | 0.0326 | 0.0891 | 0.0091 |
| | No. CF | Lectera | Ascomycota | not affected | 0.2476 | 0.0249 | 0.0799 | 0.1859 |
| | No. CF | Lycoperdon | Basidiomycota | not affected | 0.2508 | 0.0230 | 0.0799 | 0.0305 |
| | No. CF | Phallus | Basidiomycota | decreased | 0.2771 | 0.0117 | 0.0514 | 0.7397 |
| | No. CF | Podospora | Ascomycota | decreased | -0.2432 | 0.0277 | 0.0834 | 0.7179 |
| | No. CF | Psathyrella | Basidiomycota | not affected | 0.2874 | 0.0088 | 0.0471 | 0.0636 |
| | No. CF | Rhizophagus | Glomeromycota | not affected | -0.2900 | 0.0082 | 0.0471 | 0.0974 |
| | No. CF | Rhodotorula | Basidiomycota | increased | 0.3051 | 0.0053 | 0.0427 | 0.0305 |
| | No. CF | Scytalidium | Ascomycota | decreased | 0.2353 | 0.0333 | 0.0891 | 0.0351 |
| | No. CF | Solicoccozyma | Basidiomycota | decreased | -0.3302 | 0.0025 | 0.0295 | 0.2438 |
| | No. CF | Stachybotrys | Ascomycota | decreased | 0.3056 | 0.0052 | 0.0427 | 0.0526 |
| | No. CF | Thielavia | Ascomycota | not affected | 0.2838 | 0.0098 | 0.0471 | 0.0115 |

[a]Mean relative abundance (%) was calculated across all samples for the two treatments combined. Explanation of the acronyms of the root phenotypes is reported in Table 1 in the main text.

**Table S9** Complete list reporting the significant (Spearman, q < 0.1) correlations found between root phenotypes and relative abundances of prokaryotic genera for well- and lower-performing genotypes under drought and control conditions.

| Prokaryotes | | | | | | | | | |
|---|---|---|---|---|---|---|---|---|---|
| **Treatment** | **Group** | **Root phenotype** | **Genus** | **Phylum** | **Relative abundance under drought comapred to control** | **Corr (ρ)** | **P** | **q-value** | **Mean relative abundance (%)** |
| Drought | Lower-performing | Diameter mean | Chryseolinea | Bacteroidota | decreased | -0.8881 | 0.0001 | 0.0213 | 0.0305 |
| | | Diameter max | Actinoplanes | Actinobacteriota | not affected | 0.7902 | 0.0022 | 0.0648 | 0.1141 |
| | | Diameter max | Chryseolinea | Bacteroidota | decreased | -0.7692 | 0.0034 | 0.0754 | 0.0305 |
| | | Diameter max | Clostridium sensu stricto 10 | Firmicutes | not affected | -0.7413 | 0.0058 | 0.0846 | 0.0327 |
| | | Diameter max | Dethiobacter | Firmicutes | not affected | -0.7531 | 0.0047 | 0.0846 | 0.0070 |
| | | Diameter max | Ferruginibacter | Bacteroidota | decreased | -0.8252 | 0.0010 | 0.0416 | 0.1592 |
| | | Diameter max | Fimbriiglobus | Planctomycetota | decreased | -0.8462 | 0.0005 | 0.0416 | 0.0476 |
| | | Diameter max | Herpetosiphon | Chloroflexi | not affected | -0.7692 | 0.0034 | 0.0754 | 0.0948 |
| | | Diameter max | Iamia | Actinobacteriota | not affected | -0.7273 | 0.0074 | 0.0990 | 0.1061 |
| | | Diameter max | Luteitalea | Acidobacteriota | not affected | -0.7972 | 0.0019 | 0.0648 | 0.0346 |
| | | Diameter max | Opitutus | Verrucomicrobiota | increased | -0.8322 | 0.0008 | 0.0416 | 0.1073 |
| | | Diameter max | Parasegetibacter | Bacteroidota | increased | -0.8252 | 0.0010 | 0.0416 | 0.0900 |
| | | Diameter max | Rhodanobacter | Proteobacteria | decreased | 0.7483 | 0.0051 | 0.0846 | 0.2077 |
| | | Diameter max | Roseimicrobium | Verrucomicrobiota | not affected | -0.7413 | 0.0058 | 0.0846 | 0.1398 |
| | | RXSA | Lacunisphaera | Verrucomicrobiota | decreased | -0.8521 | 0.0004 | 0.0622 | 0.0289 |
| | | RXSA | OM27 clade | Bdellovibrionota | decreased | -0.8881 | 0.0001 | 0.0329 | 0.1294 |
| | | TSA | Lacunisphaera | Verrucomicrobiota | decreased | -0.9296 | 0.00001 | 0.0025 | 0.0289 |
| | | TSA | OM27 clade | Bdellovibrionota | decreased | -0.8811 | 0.0002 | 0.0156 | 0.1294 |
| | | C:XS | Acidibacter | Proteobacteria | decreased | 0.8182 | 0.0011 | 0.1000 | 0.4532 |
| | | C:XS | AKYG587 | Planctomycetota | decreased | 0.8182 | 0.0011 | 0.1000 | 0.0211 |
| | | C:XS | Nakamurella | Actinobacteriota | decreased | -0.8741 | 0.0002 | 0.0527 | 0.1551 |
| | | S:XS | Acidibacter | Proteobacteria | decreased | -0.8182 | 0.0011 | 0.1000 | 0.4532 |
| | | S:XS | AKYG587 | Planctomycetota | decreased | -0.8182 | 0.0011 | 0.1000 | 0.0211 |
| | | S:XS | Nakamurella | Actinobacteriota | decreased | 0.8741 | 0.0002 | 0.0527 | 0.1551 |
| | | No. MX | Abditibacterium | Abditibacteriota | not affected | 0.7649 | 0.0038 | 0.0916 | 0.0129 |
| | | No. MX | Acidibacter | Proteobacteria | decreased | -0.7439 | 0.0055 | 0.0945 | 0.4532 |
| | | No. MX | Aridibacter | Acidobacteriota | decreased | 0.7439 | 0.0055 | 0.0945 | 0.0175 |
| | | No. MX | Deinococcus | Deinococcota | not affected | 0.8386 | 0.0007 | 0.0318 | 0.0279 |
| | | No. MX | Dongia | Proteobacteria | decreased | -0.8351 | 0.0007 | 0.0318 | 0.0907 |
| | | No. MX | Iamia | Actinobacteriota | not affected | -0.7368 | 0.0063 | 0.0974 | 0.1061 |
| | | No. MX | Lacunisphaera | Verrucomicrobiota | decreased | -0.9329 | 0.00001 | 0.0021 | 0.0289 |
| | | No. MX | Microbispora | Actinobacteriota | decreased | -0.7434 | 0.0056 | 0.0945 | 0.0170 |
| | | No. MX | MND1 | Proteobacteria | not affected | -0.8526 | 0.0004 | 0.0311 | 0.4283 |
| | | No. MX | OM27 clade | Bdellovibrionota | decreased | -0.8807 | 0.0002 | 0.0171 | 0.1294 |
| | | No. MX | Opitutus | Verrucomicrobiota | increased | -0.7474 | 0.0052 | 0.0945 | 0.1073 |
| | | No. MX | Pedococcus-Phycicoccus | Actinobacteriota | increased | 0.7684 | 0.0035 | 0.0916 | 0.0681 |
| | | No. MX | Pedomicrobium | Proteobacteria | decreased | -0.8140 | 0.0013 | 0.0465 | 0.2147 |
| | | No. MX | Rhizocola | Actinobacteriota | decreased | -0.7333 | 0.0066 | 0.0974 | 0.0423 |
| | | No. MX | Terrimonas | Bacteroidota | decreased | -0.7860 | 0.0024 | 0.0766 | 0.0868 |
| | Well-performing | Angle mean | Luedemannella | Actinobacteriota | decreased | 0.8811 | 0.0002 | 0.0366 | 0.0322 |
| | | Diameter mean | Actinoallomurus | Actinobacteriota | increased | 0.8671 | 0.0003 | 0.0331 | 0.0960 |
| | | Diameter mean | Stenotrophobacter | Acidobacteriota | increased | -0.8671 | 0.0003 | 0.0331 | 0.0201 |
| | | Diameter min | Puia | Bacteroidota | decreased | 0.8881 | 0.0001 | 0.0340 | 0.1016 |
| | | [illegible] | [illegible] | [illegible] | | [illegible] | [illegible] | [illegible] | [illegible] |

| | | | | | | | | | |
|---|---|---|---|---|---|---|---|---|---|
| | | Diameter max | Ammoniphilus | Firmicutes | not affected | 0.8636 | 0.0006 | 0.0615 | 0.2539 |
| | | Diameter max | Brevibacillus | Firmicutes | not affected | 0.8182 | 0.0021 | 0.0698 | 0.0551 |
| | | Diameter max | Dactylosporangium | Actinobacteriota | decreased | 0.8182 | 0.0021 | 0.0698 | 0.1260 |
| | | Diameter max | Desulfitibacter | Firmicutes | not affected | 0.8818 | 0.0003 | 0.0615 | 0.0202 |
| | Lower-performing | Diameter max | Thermobacillus | Firmicutes | not affected | 0.8182 | 0.0021 | 0.0698 | 0.5233 |
| | | Diameter max | Thermopolyspora | Actinobacteriota | increased | 0.8182 | 0.0021 | 0.0698 | 0.0298 |
| | | C:S | Clostridium sensu stricto 12 | Firmicutes | not affected | 0.8545 | 0.0008 | 0.0828 | 0.0149 |
| | | C:S | IS-44 | Proteobacteria | decreased | -0.8545 | 0.0008 | 0.0828 | 0.0178 |
| | | No. MX | Edaphobacter | Acidobacteriota | decreased | -0.8676 | 0.0005 | 0.0884 | 0.0090 |
| | | No. MX | IS-44 | Proteobacteria | decreased | 0.9522 | 0.00001 | 0.0020 | 0.0178 |
| | | Angle mean | Acidipila-Silvibacterium | Acidobacteriota | decreased | -0.6993 | 0.0114 | 0.0949 | 0.0250 |
| | | Angle mean | Bradyrhizobium | Proteobacteria | decreased | -0.7902 | 0.0022 | 0.0574 | 0.5587 |
| | | Angle mean | Dyella | Proteobacteria | decreased | -0.7273 | 0.0074 | 0.0743 | 0.0277 |
| | | Angle mean | Ellin516 | Verrucomicrobiota | decreased | -0.7762 | 0.0030 | 0.0574 | 0.0207 |
| | | Angle mean | Filomicrobium | Proteobacteria | not affected | 0.8112 | 0.0014 | 0.0574 | 0.0202 |
| | | Angle mean | Jatrophihabitans | Actinobacteriota | not affected | -0.7413 | 0.0058 | 0.0655 | 0.1307 |
| | | Angle mean | Luteitalea | Acidobacteriota | not affected | 0.8671 | 0.0003 | 0.0498 | 0.0346 |
| | | Angle mean | Lysobacter | Proteobacteria | increased | 0.7622 | 0.0040 | 0.0655 | 0.2840 |
| | | Angle mean | Micromonospora | Actinobacteriota | decreased | 0.7413 | 0.0058 | 0.0655 | 0.0372 |
| | | Angle mean | MND1 | Proteobacteria | not affected | 0.7832 | 0.0026 | 0.0574 | 0.4283 |
| | | Angle mean | Nakamurella | Actinobacteriota | decreased | -0.7413 | 0.0058 | 0.0655 | 0.1551 |
| | | Angle mean | Nitrolancea | Chloroflexi | increased | -0.7203 | 0.0082 | 0.0753 | 0.2224 |
| | | Angle mean | Novosphingobium | Proteobacteria | decreased | -0.7203 | 0.0082 | 0.0753 | 0.0542 |
| | | Angle mean | Ohtaekwangia | Bacteroidota | increased | 0.7832 | 0.0026 | 0.0574 | 0.0606 |
| | | Angle mean | Phenylobacterium | Proteobacteria | decreased | -0.8182 | 0.0011 | 0.0574 | 0.1095 |
| | | Angle mean | Pirellula | Planctomycetota | decreased | 0.8252 | 0.0010 | 0.0574 | 0.3955 |
| | | Angle mean | Pseudomonas | Proteobacteria | increased | 0.7343 | 0.0065 | 0.0697 | 0.2531 |
| | | Angle mean | RB41 | Acidobacteriota | decreased | 0.7762 | 0.0030 | 0.0574 | 0.6379 |
| | | Angle mean | Rhizocola | Actinobacteriota | decreased | 0.7510 | 0.0049 | 0.0655 | 0.0423 |
| | | Angle mean | Rhodanobacter | Proteobacteria | decreased | -0.7762 | 0.0030 | 0.0574 | 0.2077 |
| | | Angle mean | Solirubrobacter | Actinobacteriota | not affected | 0.7133 | 0.0092 | 0.0803 | 0.4549 |
| | | Angle mean | Sorangium | Myxococcota | decreased | 0.7483 | 0.0051 | 0.0655 | 0.0202 |
| | | Angle mean | Tundrisphaera | Planctomycetota | decreased | -0.7413 | 0.0058 | 0.0655 | 0.1125 |
| | | Angle max | Acidipila-Silvibacterium | Acidobacteriota | decreased | -0.8392 | 0.0006 | 0.0531 | 0.0250 |
| | | Angle max | Ellin516 | Verrucomicrobiota | decreased | -0.8042 | 0.0016 | 0.0801 | 0.0207 |
| | | Angle max | Luteitalea | Acidobacteriota | not affected | 0.8042 | 0.0016 | 0.0801 | 0.0346 |
| | | Angle max | Nannocystis | Myxococcota | decreased | 0.7832 | 0.0026 | 0.0916 | 0.0229 |
| | | Angle max | Pirellula | Planctomycetota | decreased | 0.7832 | 0.0026 | 0.0916 | 0.3955 |
| | | Angle max | Pseudomonas | Proteobacteria | increased | 0.8462 | 0.0005 | 0.0531 | 0.2531 |
| | | Angle max | Rhodanobacter | Proteobacteria | decreased | -0.8392 | 0.0006 | 0.0531 | 0.2077 |
| | | RXSA | Acidicaldus | Proteobacteria | increased | -0.6853 | 0.0139 | 0.0978 | 0.0196 |
| | | RXSA | Angustibacter | Actinobacteriota | increased | -0.7133 | 0.0092 | 0.0809 | 0.1046 |
| | | RXSA | Bradyrhizobium | Proteobacteria | decreased | -0.7552 | 0.0045 | 0.0661 | 0.5587 |
| | | RXSA | Candidatus Alysiosphaera | Proteobacteria | decreased | 0.8462 | 0.0005 | 0.0599 | 0.1069 |
| | | RXSA | CL500-29 marine group | Actinobacteriota | decreased | 0.7692 | 0.0034 | 0.0661 | 0.0857 |
| | | RXSA | Coxiella | Proteobacteria | decreased | 0.6993 | 0.0114 | 0.0870 | 0.0139 |
| | | RXSA | Deinococcus | Deinococcota | not affected | -0.8182 | 0.0011 | 0.0599 | 0.0279 |
| | | RXSA | Devosia | Proteobacteria | not affected | -0.7413 | 0.0058 | 0.0719 | 0.3131 |
| | | RXSA | Granulicella | Acidobacteriota | decreased | -0.7692 | 0.0034 | 0.0661 | 0.0180 |
| | | RXSA | Iamia | Actinobacteriota | not affected | 0.7622 | 0.0040 | 0.0661 | 0.1061 |
| | | RXSA | Intrasporangium | Actinobacteriota | not affected | -0.8112 | 0.0014 | 0.0599 | 0.0092 |
| | | RXSA | Kribbella | Actinobacteriota | increased | -0.7273 | 0.0074 | 0.0719 | 0.4124 |
| | | RXSA | Leifsonia | Actinobacteriota | not affected | -0.7902 | 0.0022 | 0.0661 | 0.0082 |
| | | RXSA | Litorilinea | Chloroflexi | decreased | 0.7063 | 0.0102 | 0.0819 | 0.0387 |
| | | RXSA | Mucilaginibacter | Bacteroidota | decreased | -0.6923 | 0.0126 | 0.0923 | 0.1646 |
| | | RXSA | Nocardia | Actinobacteriota | increased | -0.7552 | 0.0045 | 0.0661 | 0.0912 |
| | | RXSA | Pantoea | Proteobacteria | increased | -0.7461 | 0.0053 | 0.0719 | 0.0759 |
| Control | | RXSA | Pedomicrobium | Proteobacteria | decreased | 0.7273 | 0.0074 | 0.0719 | 0.2147 |
| | | RXSA | Phenylobacterium | Proteobacteria | decreased | -0.7622 | 0.0040 | 0.0661 | 0.1095 |
| | | RXSA | Pir4 lineage | Planctomycetota | decreased | 0.7273 | 0.0074 | 0.0719 | 0.1831 |
| | | RXSA | Pirellula | Planctomycetota | decreased | 0.7133 | 0.0092 | 0.0809 | 0.3955 |
| | | RXSA | possible genus 04 | Fibrobacterota | decreased | -0.7762 | 0.0030 | 0.0661 | 0.0454 |
| | Well-performing | RXSA | Pseudolabrys | Proteobacteria | not affected | -0.8322 | 0.0008 | 0.0599 | 0.2504 |
| | | RXSA | Rhodomicrobium | Proteobacteria | decreased | 0.7273 | 0.0074 | 0.0719 | 0.0053 |
| | | RXSA | Roseimicrobium | Verrucomicrobiota | not affected | 0.7063 | 0.0102 | 0.0819 | 0.1398 |
| | | TCA | Angustibacter | Actinobacteriota | increased | -0.7203 | 0.0082 | 0.0964 | 0.1046 |
| | | TCA | Bradyrhizobium | Proteobacteria | decreased | -0.7203 | 0.0082 | 0.0964 | 0.5587 |
| | | TCA | Candidatus Alysiosphaera | Proteobacteria | decreased | 0.8811 | 0.0002 | 0.0305 | 0.1069 |
| | | TCA | Candidatus Protochlamydia | Verrucomicrobiota | decreased | 0.7040 | 0.0106 | 0.0964 | 0.0097 |
| | | TCA | Candidatus Xiphinematobacter | Verrucomicrobiota | decreased | 0.7063 | 0.0102 | 0.0964 | 0.1304 |
| | | TCA | CL500-29 marine group | Actinobacteriota | decreased | 0.7622 | 0.0040 | 0.0931 | 0.0857 |
| | | TCA | Coxiella | Proteobacteria | decreased | 0.7133 | 0.0092 | 0.0964 | 0.0139 |
| | | TCA | Deinococcus | Deinococcota | not affected | -0.7902 | 0.0022 | 0.0862 | 0.0279 |
| | | TCA | Devosia | Proteobacteria | not affected | -0.7483 | 0.0051 | 0.0931 | 0.3131 |
| | | TCA | Granulicella | Acidobacteriota | decreased | -0.8322 | 0.0008 | 0.0634 | 0.0180 |
| | | TCA | Iamia | Actinobacteriota | not affected | 0.7273 | 0.0074 | 0.0964 | 0.1061 |
| | | TCA | Intrasporangium | Actinobacteriota | not affected | -0.7832 | 0.0026 | 0.0862 | 0.0092 |
| | | TCA | Kribbella | Actinobacteriota | increased | -0.7483 | 0.0051 | 0.0931 | 0.4124 |
| | | TCA | Leifsonia | Actinobacteriota | not affected | -0.7972 | 0.0019 | 0.0862 | 0.0082 |
| | | TCA | Nocardia | Actinobacteriota | increased | -0.7133 | 0.0092 | 0.0964 | 0.0912 |
| | | TCA | Pantoea | Proteobacteria | increased | -0.7215 | 0.0081 | 0.0964 | 0.0759 |
| | | TCA | Phenylobacterium | Proteobacteria | decreased | -0.7622 | 0.0040 | 0.0931 | 0.1095 |
| | | TCA | Pir4 lineage | Planctomycetota | decreased | 0.7063 | 0.0102 | 0.0964 | 0.1831 |
| | | TCA | Pirellula | Planctomycetota | decreased | 0.7343 | 0.0065 | 0.0964 | 0.3955 |
| | | TCA | possible genus 04 | Fibrobacterota | decreased | -0.7483 | 0.0051 | 0.0931 | 0.0454 |
| | | TCA | Pseudolabrys | Proteobacteria | not affected | -0.8252 | 0.0010 | 0.0634 | 0.2504 |
| | | TCA | Tepidisphaera | Planctomycetota | decreased | -0.7203 | 0.0082 | 0.0964 | 0.0628 |
| | | TCA | Tundrisphaera | Planctomycetota | decreased | -0.6993 | 0.0114 | 0.0989 | 0.1125 |
| | | TSA | Nocardia | Actinobacteriota | increased | -0.8252 | 0.0010 | 0.0992 | 0.0912 |
| | | TSA | possible genus 04 | Fibrobacterota | decreased | -0.8392 | 0.0006 | 0.0992 | 0.0454 |
| | | nonAA | Angustibacter | Actinobacteriota | increased | -0.7203 | 0.0082 | 0.0964 | 0.1046 |
| | | nonAA | Bradyrhizobium | Proteobacteria | decreased | -0.7203 | 0.0082 | 0.0964 | 0.5587 |
| | | nonAA | Candidatus Alysiosphaera | Proteobacteria | decreased | 0.8811 | 0.0002 | 0.0305 | 0.1069 |
| | | nonAA | Candidatus Protochlamydia | Verrucomicrobiota | decreased | 0.7040 | 0.0106 | 0.0964 | 0.0097 |
| | | nonAA | Candidatus Xiphinematobacter | Verrucomicrobiota | decreased | 0.7063 | 0.0102 | 0.0964 | 0.1304 |
| | | nonAA | CL500-29 marine group | Actinobacteriota | decreased | 0.7622 | 0.0040 | 0.0931 | 0.0857 |
| | | nonAA | Coxiella | Proteobacteria | decreased | 0.7133 | 0.0092 | 0.0964 | 0.0139 |
| | | nonAA | Deinococcus | Deinococcota | not affected | -0.7902 | 0.0022 | 0.0862 | 0.0279 |
| | | nonAA | Devosia | Proteobacteria | not affected | -0.7483 | 0.0051 | 0.0931 | 0.3131 |
| | | nonAA | Granulicella | Acidobacteriota | decreased | -0.8322 | 0.0008 | 0.0634 | 0.0180 |
| | | nonAA | Iamia | Actinobacteriota | not affected | 0.7273 | 0.0074 | 0.0964 | 0.1061 |
| | | nonAA | Intrasporangium | Actinobacteriota | not affected | -0.7832 | 0.0026 | 0.0862 | 0.0092 |
| | | nonAA | Kribbella | Actinobacteriota | increased | -0.7483 | 0.0051 | 0.0931 | 0.4124 |
| | | nonAA | Leifsonia | Actinobacteriota | not affected | -0.7972 | 0.0019 | 0.0862 | 0.0082 |
| | | nonAA | Nocardia | Actinobacteriota | increased | -0.7133 | 0.0092 | 0.0964 | 0.0912 |
| | | nonAA | Pantoea | Proteobacteria | increased | -0.7215 | 0.0081 | 0.0964 | 0.0759 |
| | | nonAA | Phenylobacterium | Proteobacteria | decreased | -0.7622 | 0.0040 | 0.0931 | 0.1095 |
| | | nonAA | Pir4 lineage | Planctomycetota | decreased | 0.7063 | 0.0102 | 0.0964 | 0.1831 |
| | | nonAA | Pirellula | Planctomycetota | decreased | 0.7343 | 0.0065 | 0.0964 | 0.3955 |
| | | nonAA | possible genus 04 | Fibrobacterota | decreased | -0.7483 | 0.0051 | 0.0931 | 0.0454 |
| | | nonAA | Pseudolabrys | Proteobacteria | not affected | -0.8252 | 0.0010 | 0.0634 | 0.2504 |
| | | nonAA | Tepidisphaera | Planctomycetota | decreased | -0.7203 | 0.0082 | 0.0964 | 0.0628 |
| | | nonAA | Tundrisphaera | Planctomycetota | decreased | -0.6993 | 0.0114 | 0.0989 | 0.1125 |
| | | No. MX | Candidatus Alysiosphaera | Proteobacteria | decreased | 0.8531 | 0.0004 | 0.0395 | 0.1069 |
| | | No. MX | Leifsonia | Actinobacteriota | not affected | -0.8741 | 0.0002 | 0.0379 | 0.0082 |

[a]Mean relative abundance (%) was calculated across all samples for the two treatments and performance groups combined. The complete description of the acronyms of the root phenotypes is in the main text (Table 1).

**Table S10** Complete list reporting the significant (Spearman, q < 0.1) correlations found between root phenotypes and relative abundances of fungal genera for well- and lower-performing genotypes under drought and control conditions.

| Fungi | | | | | | | | | |
|---|---|---|---|---|---|---|---|---|---|
| Treatment | Group | Root phenotype | Genus | Phylum | Relative abundance under drought comapred to control | Corr ($\rho$) | P | q-value | Mean relative abundance (%) |
| Drought | Lower-performing | NRN max | Lecythophora | Ascomycota | decreased | 0.8334 | 0.0008 | 0.0804 | 0.2449 |
| | | Angle min | Trichaptum | Basidiomycota | decreased | 0.8231 | 0.0010 | 0.0994 | 0.0056 |
| | | TSA | Sporobolomyces | Basidiomycota | increased | 0.8322 | 0.0008 | 0.0833 | 0.0371 |
| | | No. MX | Sporobolomyces | Basidiomycota | increased | 0.8912 | 0.0001 | 0.0106 | 0.0371 |
| Control | Well-performing | LRBD | Sarocladium | Ascomycota | not affected | 0.9161 | 0.00003 | 0.0018 | 0.1408 |

[a]Mean relative abundance (%) was calculated across all samples for the two treatments and performance groups combined. The complete description of the acronyms of the root phenotypes is in the main text (Table 1).

**Notes S1** Effects of drought on grain yield and identification of plant performance groups.

Relative change in grain yield was used to evaluate the impact of drought stress on each genotype (Fig. S5). It was calculated for 17 out of 22 genotypes based on the availability of corresponding replicates of the same genotype coupled to perform this calculation (see Materials and Methods). Twelve out of 17 genotypes were considered significantly different from zero (control baseline) based on the 95% confidence intervals of the estimated marginal means (EMMs). EMMs were calculated on the output of the linear mixed-effects model testing the effect of the genotype on relative yield change.

**Notes S2** Grain yield performance-based differences in root phenotypes and microbiomes.

Patterns in expression of anatomical and architectural phenotypes did not differentiate clearly between the two yield-based performance groups under either drought or control conditions (Fig. S12). This was further confirmed by the fact that the anatomy-based k-medoid clusters under drought did not overlap with the performance groups (Fig. S13). Only the two well-performing IBM167 and IBM301 planted in the LongROS were characterized by a similar greater root cross-section and stele areas, and number of metaxylem vessels. However, the opposite trend was

observed for the well-performing IBM313 belonging to the ShortROS (Fig. S8). Nevertheless, the percentage of cortex that is aerenchyma (percCisA) significantly (PERMANOVA, $P < 0.05$) increased in lower performing compared to well-performing genotypes (Fig. S12).
Moreover, the well- and lower-performing genotypes differed in their rhizosphere prokaryotic and fungal communities when constraining the entire variability by performance groups (CAP, Fig. S14). A greater separation was observed for the fungal communities under drought.

**Methods S1** Data curation for plant performance, root phenotypes and bulk soil properties.

After thorough review and pairing with the respective design information, shoot biomass, vigor and root phenotypic data were gap filled where possible. Missing values (NAs) for the specific plant analyzed were replaced by single or average values of one, two or three plants grown in the same plot. We assumed that the difference between these plants was not great since they belonged to the same genotype, plot, block and treatment. NAs were not imputed for the yield data to avoid introducing biases. Thirteen samples for architecture (control: IBM200 rep2, IBM181 rep2, IBM317 rep2, IBM345 rep2; drought: IBM351 rep4, IBM345 rep3, IBM344 rep4, IBM199 rep3, IBM090 rep3, IBM097 rep3, IBM345 rep2, IBM153 rep3, IBM097 rep4) and four for anatomy (control: IBM317 rep2, IBM345 rep2; drought: IBM345 rep2, IBM153 rep3) were removed to match the length of the two datasets and to keep only the samples with complete entries for all measured variables and available for DNA metabarcoding (see Table S2). A sample showing too little yield compared to the others (IBM365, Short ROS, rep 2) was removed from all plant and microbial datasets as well. Regarding root architecture, the total number of whorls corresponded to the last possible whorl for which architectural data were available. Minimum, maximum and average number of nodal roots, diameter and angle were calculated across all available whorls. Relative yield change, root phenotypic and bulk soil data were z-transformed (*scale* function) to perform statistics, correlations and clustering. The effect of the main experimental factors (drought and genotypes) on the measured variables was not completely masked by the differences between the two field locations (Short and Long ROS), which were mostly based on pH, cation exchange capacity (CEC), total nitrogen and carbon and macro- and micronutrient content (Fig. S1, Table S1). We always included the field as a random factor or constraining factor for permutations in the statistical models and normalized root phenotypes and relative yield change by field to minimize biases.

**Methods S2** Protocols used to characterize physiochemical properties, and microbial communities of bulk soil.

The following methods were already described in a previous work (Giuliano et al. 2026). Briefly, after -80°C storage and overnight thawing (4°C), bulk soil samples were sieved (2 mm). Dried soil (10 g, 105°C, 72 h) was used to measure gravimetric water content (GWC). Moist soil (10 g) was used to measure ammonium ($NH_4^+$) and nitrate ($NO_3^-$) content by measuring the absorbance at 650 nm and 540 nm with a visible spectrophotometer (V-1200, VWR International), respectively, after 1 hour shaking in 2 M KCl and filtering through Whatman® 42 mm filters (GE Healthcare Life Science Whatman™, Chicago, USA). A C/N analyzer (CN628, LECO Corporation, St. Joseph, USA) was used to quantify total nitrogen (N) and carbon (C) on 200 mg of dry (60°C, 72 h) and milled soil. A multi-parameter meter (pHenomonal® MU6100L, VWR Int., Radnor, USA) was used to measure pH after suspending dried soil (10 g, 40°C, 5 days) in 25 ml of $ddH_2O$, shaking for 24 hours and settling down for 24 hours. Milled and dried soil (2.5 g, 60°C, 72 h) was shaken in 0.1 M $BaCl_2$ for 2 hours, centrifuged (2500 rpm, 10 minutes) and filtered (Whatman® 41 mm filter papers, GE Healthcare Life Science Whatman™) to measure effective cation exchange ($CEC_{eff}$) capacity following Hendershot and Duquette (1986). $CEC_{eff}$ calculation considered the concentrations of potassium, calcium, magnesium, sodium, hydrogen, aluminium, manganese and iron. Dried (1 g, 60°C, 72 h) and milled soil dissolved first in 2 mL $ddH_2O$, and then in 2 mL $HNO_3$ 70% and 6 mL HCl 37% was digested (120°C, 90 min, DigiPREP MS, Baie-D'Urfé, Canada) and filtered (Whatman® 41 mm filter, GE Healthcare Life Science Whatman™) to quantify soil macro- and micronutrients. The filtrates for CEC and total nutrients were analyzed with ICP-OES (SVDV, 5100 ICP-OES, Agilent, US). Soil was always milled with the MM200 mill (Retsch GmbH, Haan, Germany). These bulk soil samples were used to characterize differences in microbial community structure due to treatment or field. The same rhizosphere soil protocol was used, but in this case 3 g of sieved bulk soil were dissolved in autoclaved phosphate buffer (8.75 g $K_2HPO_4$, 6.75 g of $KH_2PO_4$ and 200 µl of Tween 20 in 1 l $H_2O$) to have an amount of final pellet comparable to the rhizosphere.

**Methods S3** Statistical models applied on plant grain yield, root architectural and anatomical phenotypes, bulk soil physiochemical properties and microbial diversity (main analyses).

As mentioned in the main text, similar statistical approaches were already used and described by

Giuliano et al. (2026), and here they are adapted to the current study.

1. *Plant yield and root phenotypes:* the effect of treatment (T), maize genotype (G) and field (F) were tested with linear mixed-effect models (LMMs). Field or blocks (B) were included in the models as random factors depending on the model quality. First, models with different random but same fixed factors were compared with the restricted maximum likelihood method (RELM). Secondly, models with different fixed but same random factors were compared with the maximum likelihood method (LM). In some cases, weights = *varIdent(form = ~1 | G)* was included in the model to account for genotype-dependent heterogeneity of variance. Plots of normalized residuals, Shapiro-Wilk test (function *shapiro.test*, package *stats* v4.4.2) and Levene's test (function *leveneTest*, package *car* v3.1.3) were used to check for normality and homoscedasticity. The main model equations are the following:
   - lme(yield ~ T * G, random = ~1 | F, method = "REML", weights = varIdent(form = ~1 | G), data = yield_data)
   - lme(relativeChange_grain_yield ~ G, random = ~1 | B, method = "REML", data = yield_data)
   - lme(individual_phenotype ~ T * or + G, random = ~1 | F or B, method = "REML", data = phen_data)
2. *Bulk soil properties:* the effect of treatment and field were tested with permutational multivariate analysis of variance (PERMANOVA). The model equation used is the following:
   - mclapply(**bulksoil_data.list***, function(x) adonis2(vegdist(x, method = "euclidean") ~ T * F, data = soil_data, by = "terms", permutations = 9999))

   ***bulksoil_data.list**: list of z-transformed bulk soil properties
3. *Microbial α- and β- diversity and taxon level composition:* the effects of treatment x genotype and root phenotypes were tested with PERMANOVA. After frequently observing a field effect on microbiomes, field was always included as *strata*. If necessary, the datasets were further split by treatment. The model equations used for both treatments combined are the following:
   - adonis2(**distance*** ~ T * G, strata = design$F, data = design, by = "terms", permutations = 9999)
   - adonis2(**distance*** ~ T * G + phen1 + phen2 + phen3 etc., strata = phen_data$F, data = phen_data, by = "terms", permutations = 9999)

- lapply(**genera_data.list***, function(x) adonis2(vegdist(x, method = "euclidean") ~ T * G, strata = design$F, data = design, by = "terms", permutations = 9999))

***distance**: Euclidean distance (α-diversity) or Bray-Curtis dissimilarity (β-diversity)

***genera_data.list**: list of microbial genera or species relative abundances (Euclidean distances calculated for each component)

**Methods S4** Approaches to identify associations between plant grain yield, root architecture and anatomy and microbial diversity (additional analyses).

Additional analyses (in bold if reported in the main text) and examples of the statistical models used are listed below. The treatment factor was removed when running models on the data split by treatment. Abbreviations: treatment (T), genotype (G), field (F). Similar approaches were previously used for a different experiment and described by Giuliano et al. (2026).

1. *Root phenotypes – yield:*
    - **Effect of all phenotypes and CPW on yield by treatment (PERMANOVA)**
    - **Spearman correlations between all phenotypes and yield by treatment**
    - **Effect of plant performance groups on root phenotypes (PERMANOVA)**
        - adonis2(yield_Euclidean_dist ~ G + phen1 + phen2 + phen3 + etc., strata = phen_data$F, data = phen_data, by = "terms", permutations = 9999))
        - mclapply(***phenotype.data.list**, function(x) adonis2(vegdist(x, method = "euclidean") ~ T * (perf_groups OR raw_yield) + G, strata = yield_data$F, data = yield_data, by = "terms", permutations = 9999))

    ***phenotype_data.list**: list of all z-transformed root phenotypes
2. *Microbiomes – yield:*
    - **Effect of plant performance groups on** α-diversity and **β-diversity overall and by treatment (PERMANOVA)**
        - adonis2(**distance*** ~ perf_groups + G, strata = design$F, data = design, by = "terms", permutations = 9999))
    - Correlation between overall patterns of yield and β-diversity by treatment (Mantel test)
    - Spearman correlations between yield and genera relative abundances by treatment

    ***distance**: Euclidean distance (α-diversity) or Bray-Curtis dissimilarity (β-diversity)

3. *Root phenotypes – microbiomes:*

- **Effect of individual phenotypes on microbial β-diversity (PERMANOVA)**: see Methods S3, point 3
- **Correlation between overall patterns based on root phenotypes (Euclidean distance) and β-diversity (Bray-Curtis dissimilarity) by treatment (Mantel test)**

### References

CICG (2026) International Culture Collection of Glomeromycota. Available in: https://sites.google.com/view/cicg-furb-english/home

Giuliano E, Singh Sidhu J, Lopez-Valdivia I, et al (2026) Shovelomics meets microbiomics: root phenotype-microbiome associations and links with maize yield under nitrogen limitation. Rhizosphere 101241. https://doi.org/10.1016/J.RHISPH.2025.101241

Hendershot WH, Duquette M (1986) A Simple Barium Chloride Method for Determining Cation Exchange Capacity and Exchangeable Cations. Soil Science Society of America Journal 50:605–608. https://doi.org/10.2136/SSSAJ1986.03615995005000030013X

Klein SP, Schneider HM, Perkins AC, et al (2020) Multiple Integrated Root Phenotypes Are Associated with Improved Drought Tolerance. Plant Physiol 183:1011–1025. https://doi.org/10.1104/PP.20.00211
